# Mapping the nuclear landscape with multiplexed super-resolution fluorescence microscopy

**DOI:** 10.1101/2024.07.27.605159

**Authors:** Fariha Rahman, Victoria Augoustides, Emma Tyler, Timothy A. Daugird, Christian Arthur, Wesley R. Legant

## Abstract

The nucleus coordinates many different processes. Visualizing how these are spatially organized requires imaging protein complexes, epigenetic marks, and DNA across scales from single molecules to the whole nucleus. To accomplish this, we developed a multiplexed imaging protocol to localize 13 different nuclear targets with nanometer precision in single cells. We show that nuclear specification into active and repressive states exists along a spectrum of length scales, emerging below one micron and becoming strengthened at the nanoscale with unique organizational principles in both heterochromatin and euchromatin. HP1-α was positively correlated with DNA at the microscale but uncorrelated at the nanoscale. RNA Polymerase II, p300, and CDK9 were positively correlated at the microscale but became partitioned below 300 nm. Perturbing histone acetylation or transcription disrupted nanoscale organization but had less effect at the microscale. We envision that our imaging and analysis pipeline will be useful to reveal the organizational principles not only of the cell nucleus but also other cellular compartments.

## Introduction

Over 6000 proteins (>30 percent of the human proteome) localize to the nucleus. Of these, only 8% localize to clearly defined structures such as the nucleolus or nuclear speckles. The other 92% are diffusely distributed within the nucleoplasm where they bind to a dynamic meshwork of DNA and RNA^1^. Given the lack of partitioning membranes, there is immense interest in how the nucleus organizes different and often opposing biological functions^2,3^. Biochemical approaches such as chromatin immunoprecipitation and sequencing (ChIP-seq) and chromatin conformation capture have revealed that genomic sequences can be broadly classified as corresponding to active euchromatin (A-Type) or repressive heterochromatin (B-Type) regions. These regions are defined by their transcriptional output and the combinatorial enrichment of transcriptionally active (or repressive) proteins and histone post-translational modifications (PTMs) and self-associate into topologically associated domains (TADs) of enriched contact frequency^4,5^. However, genomic studies typically average the signal from millions of cells and there are conflicting studies as to whether these regions correspond to real physical compartments within the nuclei of single cells^6,7^ or if they are emergent phenomena from population averaging^8^. Fluorescence microscopy and chromosome tracing have revealed that TADs exhibit a high degree of heterogeneity and mixing in single cells^9^, while individual genomic loci tend to be consistently organized in proximity (≤ 300 nm) to different combinations of activation or repression-associated histone PTMs and larger nuclear landmarks such as speckles^10,11^. However, it is less clear how changes in proximity, at the scales measured via diffraction limited imaging, relate to the molecular organization of an active or repressive genomic region.

At the nanoscale, super-resolution immunofluorescence imaging has revealed that nucleosomes, epigenetic marks, and transcription-associated proteins all display unique organizations in cells. Nucleosomes coalesce into irregular groups or clusters at the scale of tens of nanometers^12,13^. Activation associated histone PTMs (H3K27Ac, H3K4me1) form dispersed nanoclusters (typically less than 100 nm) whereas repressive marks (H3K9me3) form larger aggregates that are up to several hundred nm in size^14^. Finally, transcription associated proteins such as RNA Polymerase II (RNAPII) form transient clusters that are 100-200 nm in size^15^. However, connecting the nanoscale distributions of chromatin, histone PTMs, and transcriptional machinery to the organization of active or repressive chromatin states in the nucleus requires combinatorial imaging of many targets simultaneously.

To accomplish this, we used multiplexed Exchange-PAINT labeling^16^ and highly inclined swept tile (HIST) imaging^17^ to localize 12 different antibody targets and DNA in the nuclei of individual cells. We found that at larger length scales, most of our selected nuclear proteins and epigenetic marks were nearly randomly distributed within the nucleus and share similar scaling properties to DNA. However, these same targets formed small self-associated clusters below ∼200 nm in a manner that is distinct from the chromatin scaffold. Nuclear specification of multiple targets into active or repressive regions was not delineated by sharp boundaries, but rather, emerged below one micron and became progressively strengthened at short length scales below 500 nm. Nanoscale distance measurements between individual protein pairs correlated with known protein-protein interactions and revealed higher order organizational principles in cells. For example, we found that transcription-associated targets were all enriched together at approximately one-micron length scale, but partitioned into unique environments below the diffraction limit. Finally, we studied how perturbations to histone acetylation and transcription altered nuclear organization. Together, our results reveal the multiple length scales of nuclear organization from the micro to the nanoscale and provide a quantitative platform to study the emergence of functionally specified environments in the cell nucleus.

## Results

### Thirteen-plex single molecule imaging of cell nuclei over a large field-of-view using Highly Inclined Swept Tile microscopy and multiplexed Exchange PAINT

To label nuclear targets and DNA together in chemically fixed cells, we implemented a modified indirect immunofluorescence pipeline. Primary antibodies were pre-associated to species-specific secondary nanobodies covalently modified with orthogonal oligonucleotide docking strands. Pre- associated antibody/nanobody complexes against each target were then combined and incubated together with the sample. After labeling, individual targets were imaged sequentially using dye- conjugated imager strands that were complementary to each docking strand (**Fig. 1A**). To ensure high-specificity, both docking and imager strand sequences were optimized for orthogonality and low nuclear background (**Table S1, Fig. S1-S3**). To enable high-precision, we used a custom- developed variant of HIST microscopy^17^ that leveraged a high numerical aperture (NA = 1.27) water-immersion objective for optically sectioned illumination and astigmatic detection for 3D localization (**Fig. S4, S5**). Together, this strategy allowed us to localize 13 total targets within an approximately one-micron-thick plane midway through the nuclei of many cells within a single field of view. The 3D localization precision for each label was on average 9.9/35 nm (Cramer-Rao Lower Bound, CRLB, laterally/axially) for Cy3B targets, 11.5/52.2 nm for ATTO655 targets, and 9/49.4 nm for Hoechst (**Figures S6-S17**). Overall, this represents approximately a 35,000-fold volumetric resolution improvement over confocal imaging^10^ (220/660 nm FWHM) and a 4,000- fold improvement over structured illumination microscopy^18^ (110/330 nm FWHM) (**Fig. S18-S20**) of nuclear features. Our imaging approach improves upon the precision of prior localization microscopy and stimulated emission depletion imaging studies of nuclear targets^19,20^ and extends these imaging approaches to 3D while simultaneously permitting multiplexing and quantitative evaluation of target localization via point statistics. With this approach, we targeted post- translational modifications to the histone H3 tail (H3K4me1, H3K27ac, H3K27me3, and H3K9me3) commonly associated with both gene activation and repression, RNA Polymerase II and CTD-tail phosphorylation variants S2 and S5, the transcription cofactor CDK9, chromatin “readers” and “writers” p300 and HP1α, and nuclear landmarks including nuclear speckles (SC-35) and the nuclear envelope (lamin A/C), as well as DNA (**Fig. 1B-F, Movie S1**).

**Figure 1.**
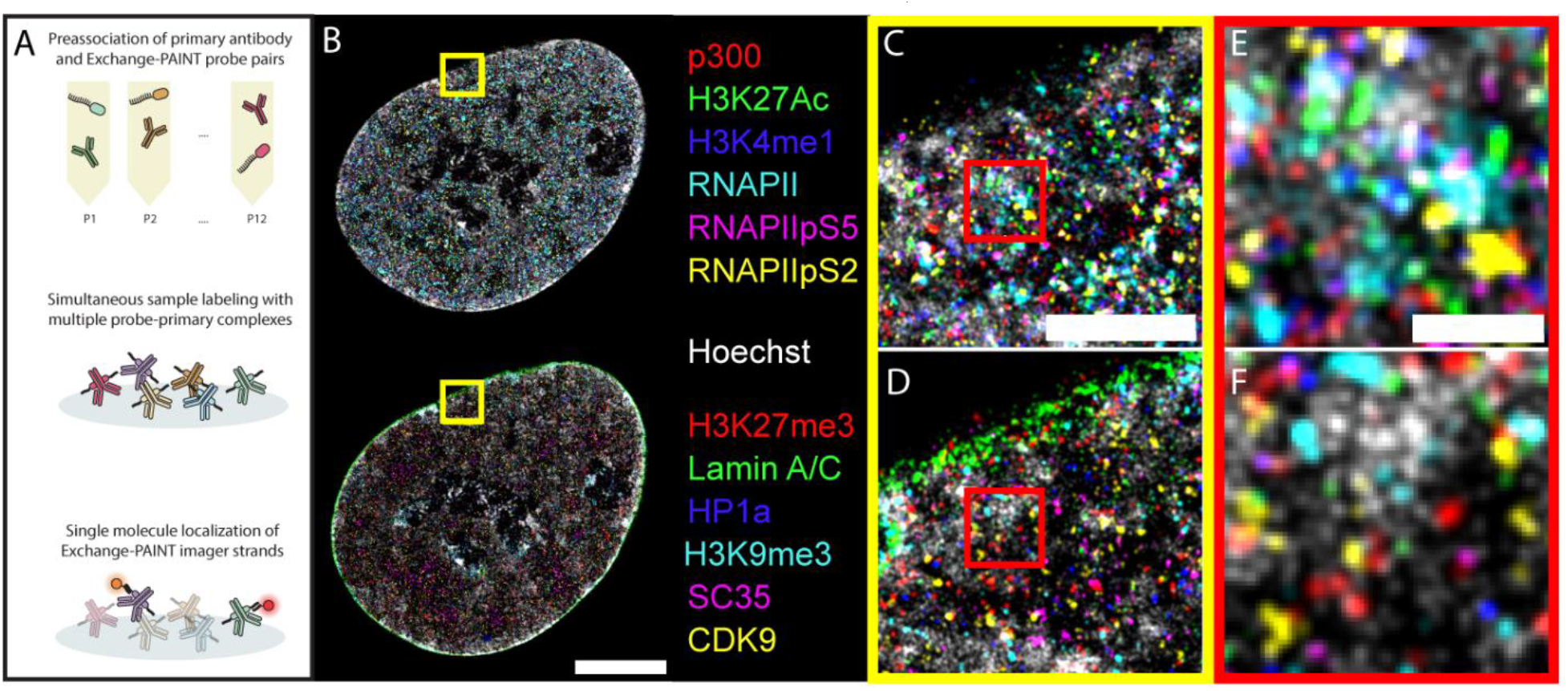
Overview of Exchange-PAINT protocol and representative super-resolution rendering of the nucleus with 13 labels. (A) Schematic of the Exchange-PAINT sample preparation and imaging pipeline. **(B)** 13- plex rendering of a representative unperturbed nucleus, scale bar 5 μm. **(C-D)** Top and bottom insets of B, respectively, scale bar 1 μm. **(E-F)** Insets of C and D, respectively, scale bar 200 nm.

### Most nuclear targets display two length scales of organization with a transition between spatial regimes at approximately 100 - 200 nm

To measure the spatial distribution of each target within the nucleus, we computed the normalized 3D pair correlation function G(r), (**Fig. 2A,B**) which quantifies the deviation from spatial randomness at a given length scale^21^. Values greater than or less than one indicate an enrichment or depletion, respectively, in the number of molecules at a given length compared to what would be expected from a spatially random distribution (G(r) = 1). The slope of the G(r) curve on a log-log plot provides information about the scaling behavior of spatial organization^22,23^. A linear slope is indicative of power law scaling which can be quantified concisely as the fractal dimension of the distribution 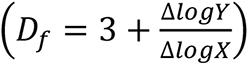 for points distributed within a 3D volume. The fractal dimension describes how optimally space-filling points are at a given length scale. *D*_*f*_ ≈ 3 indicates an optimally space filling random distribution whereas *D*_*f*_ ≈ 1 indicates a highly clustered distribution. Transitions between G(r) slope (fractal dimension) reveal changes in the organizational principles of a distribution at a given length scale. Importantly, unlike other methods to determine spatial clustering such as DBSCAN^24,25^ or tessellation-based approaches^26^ which require tunable parameters, G(r) does not make any assumptions about the target distribution such as cluster sizes or numbers. Furthermore, because G(r) is normalized by an equal number of randomly distributed points occupying the same space, it is independent of variations in label density due to antibody or probe efficiency. Further discussions about how we interpret unique aspects of the G(r) curve and its multi-species equivalent, the pairwise cross correlation function (PCCF), are provided in **Supplementary Note 1 and Fig. S21**.

**Figure 2.**
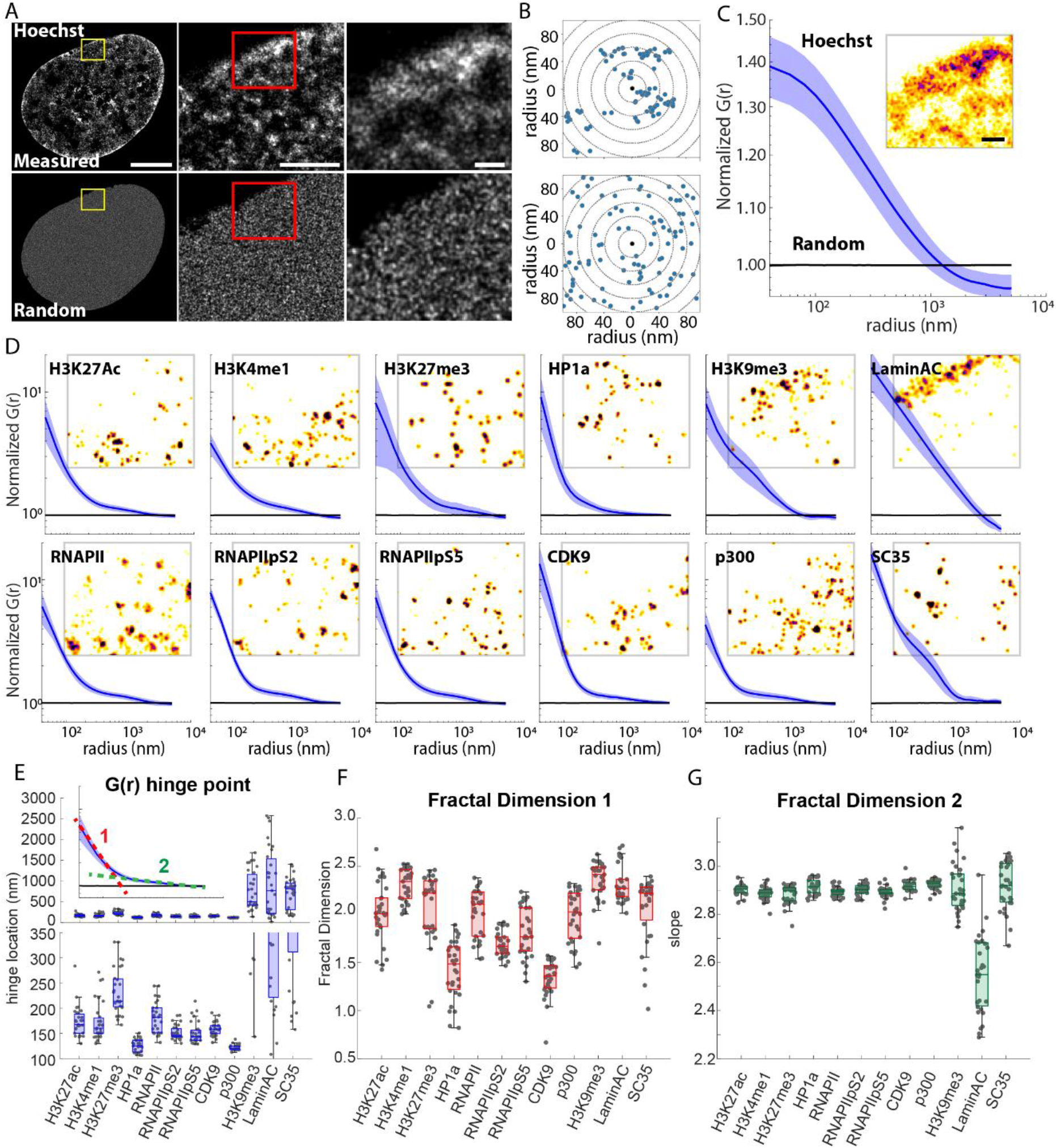
Quantification of Exchange-PAINT target localization using the pair correlation function and fractal dimension. (A) Representative rendering of a Hoechst labeled nucleus and a scrambled distribution for complete spatial randomness (CSR). **(B)** Demonstration of the pair correlation function G(r) for a simulated clustered distribution (upper) and a simulated CSR distribution (lower). **(C)** Pair correlation and representative inset for Hoechst. **(D)** Pair correlation for Exchange-PAINT targets. Representative insets are from the same region as in (A – right). **(E)** Box plots of the hinge points for bilinear fits corresponding to plots in (D). Inset shows a representative bilinear fit for H3K27Ac indicating the two linear regions. Lower panel shows a clipped y-axis to highlight changes between targets. **(F)** Box plots for the fractal dimension of the first linear region for plots in (D). **(G)** Box plots for the fractal dimension of the second linear region for plots in (D). Scale bars = 5 μm – (A – left), 1 μm – (A – center), 500 nm– (A – right and C,D - insets). Data from (C-G) are mean ± standard deviation. Data from (E-G) from n = 31 cells across 3 independent biological replicates.

In agreement with previous studies^23^, we found that the G(r) curve for DNA within the nucleus could be reasonably fit with a single linear model between 100 nm and one micron indicating a fractal dimension of 2.89 ± .016 (**Fig. 2C**). DNA distribution deviates from spatial randomness below approximately 1 micron and continues to become more spatially clustered down to approximately 100 nm. In contrast, most of the nuclear targets that we imaged displayed two clear scaling regimes (**Fig. 2D**). To account for this, we fit the log-log plotted G(r) curves to a bilinear model and reported the fractal dimensions and hinge-points between the two linear fits (**Table S2 – S4**). From this data, we found that most nuclear targets formed small clusters from the limit of our 3D localization precision of ∼40 nm up to a mean hinge point of 163 ± 41 nm with an average fractal dimension of 1.85 ± 0.39 (**Fig. 2 E,F**). At distances greater than the hinge point, the average fractal dimension increased to 2.9 ± 0.03 which was very similar to that of DNA (**Fig. 2G**). Three labels were a clear exception to this trend. The constitutive heterochromatin mark H3K9me3 and the splicing factor SC-35 still formed small clusters (with short-length fractal dimension of 2.36 ± 0.20 and 2.06 ± 0.35 respectively), but the transition point between linear regimes was substantially larger at 755 ± 456 nm and 716 ± 367 nm respectively (**Table S2 – S4**). As is visually apparent within the images and consistent with prior literature on constitutive heterochromatin and splicing speckles^10^, these two targets also formed larger, micron-scale, clusters within the nucleus. Finally, as expected, lamin A/C was highly enriched at the nuclear periphery, and transitioned from partitioned to clustered at approximately 5 microns and showed only a single linear scaling regime. These data reveal that at large length scales, most of our targets are nearly randomly dispersed throughout the nucleus and display similar scaling properties to DNA. At short length scales, both active and repressive targets form small self-associated clusters in a manner that is distinct from the underlying chromatin scaffold.

### Active and repressive nuclear environments become functionally specified below one-micron

To understand how groups of molecules coordinate to establish a functionally specialized nuclear environment, we first partitioned the localizations from our targets into those associated with active “A-Type” euchromatin and repressive “B-Type” heterochromatin^5^ (**Fig. 3A**). After combining the localizations from each category, we implemented the pair cross-correlation function (PCCF)^20^ to quantify spatial correlations between different targets. This is similar to the pair correlation function but is modified for two input localization lists. More specifically, values greater than or less than one indicate the enrichment or depletion of one molecule species relative to another at a given length compared to what would be expected from a spatially random distribution of the second molecule (PCCF = 1). In contrast to the G(r) curves, the slope of the PCCF curve cannot be summarized in terms of a fractal dimension. Nevertheless, the value and slope still describe the scaling behavior of intermolecular distributions. We found that specification into active and repressive nuclear environments occurs at length scales below approximately one micron which is substantially larger than the 100 - 200 nm clustering hinge point for the individual targets with themselves (**Fig. 3B**). Above one micron, both active and repressive labels display no self-association enrichment (PCCF ≈ 1), but below this length scale, we observe progressively enriched self-association of A-A and B-B labels. In addition to increased homotypic clustering, we also observe increased heterotypic partitioning between A and B classified targets below one micron (PCCF < 1). Above one micron, the PCCF curve between A-Type and B-Type targets is roughly equal to one. Together, these curves demonstrate that above one micron, chromatin does not appear to be specialized for either euchromatin or heterochromatin functions. However, below this length scale, it partitions into progressively specialized functional environments.

**Figure 3.**
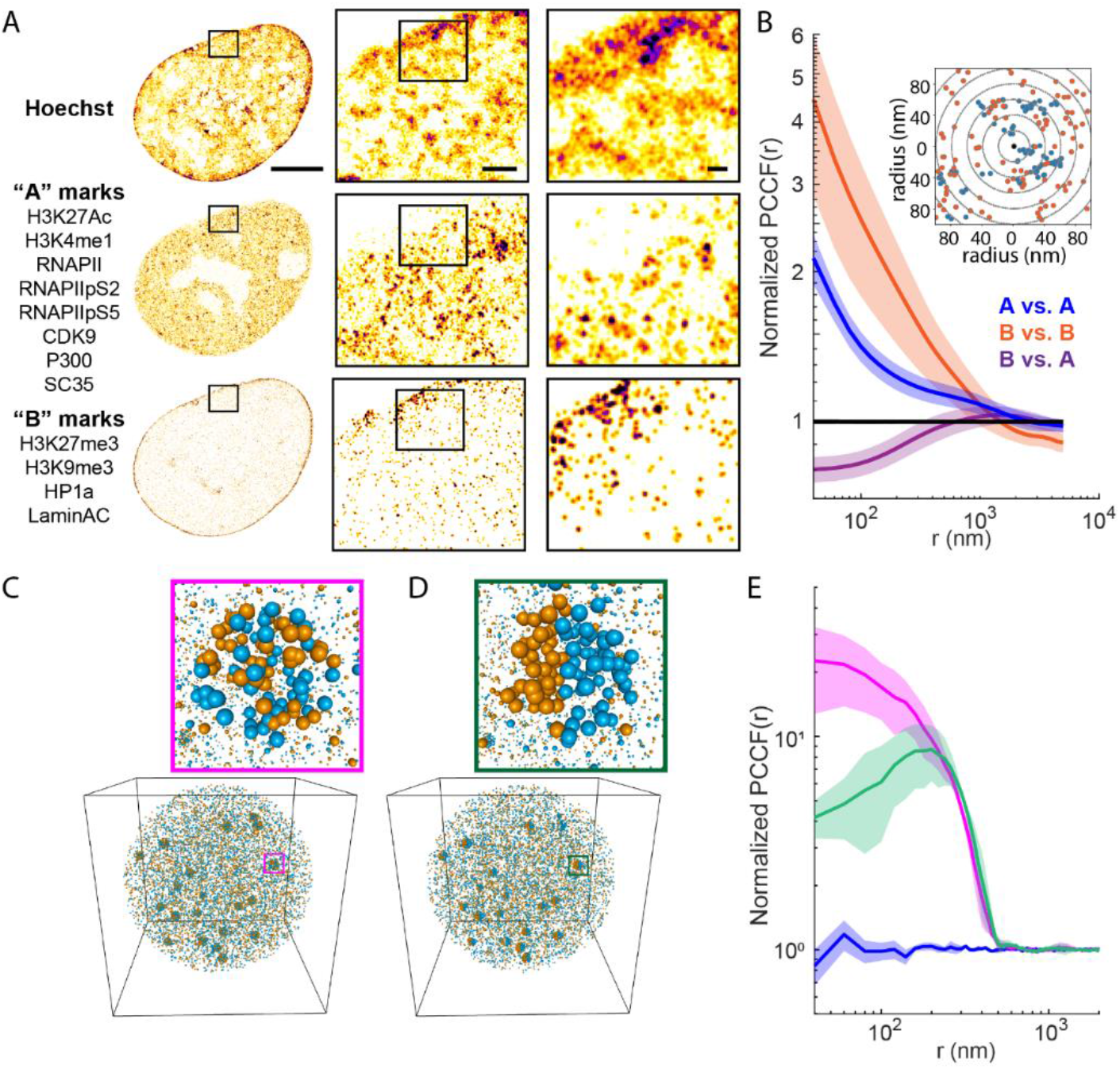
Quantification of A-Type and B-Type chromatin distributions using the pair cross correlation function. (A) Representative renderings of Hoechst, and aggregated A-type and B-type Exchange-PAINT targets. **(B)** Pair correlation and cross-correlation functions for aggregated labels. Inset - Demonstration of the pair cross-correlation function (PCCF(r)) for two different labels. **(C)** Simulated point distributions for two species wherein 20% of the molecules of each species are enriched together within 500 nm diameter clusters yet display no correlation within a cluster. **(D)** Simulated point distributions for two species wherein 20% of the molecules of each species are enriched together within 500 nm diameter clusters and partition to different halves within a cluster. **(E)** PCCF outputs corresponding to (C - magenta), (D - green), and CSR for each species (blue). Scale bars = 5 μm – (A – left), 500 nm – (A – center), 200 nm– (A – right). Data from (B, E) are mean ± standard deviation. Data in (B) are from n = 31 cells across 3 independent biological replicates. Data in (E) are from 3 independent simulated distributions.

### Exploring the PCCF behavior using simulated distributions

To better explore the relationship between PCCF curves and the distributions of nuclear targets, we turned to simulated datasets that were inspired by potential biophysical mechanisms including direct physical association and clustering or condensate formation (**Fig. 3C-D**). If we zoom in from the length scale of the whole nucleus to progressively smaller domains, we find that most of our measured targets and target-target pairs become progressively enriched or depleted at short length scales. The transition at which the curve deviates from a random distribution can be viewed as the length scale at which organization (e.g. clustering or depletion) begins to emerge. As we move from large to small radii, transitions in the slope of the curve suggest a switch between enrichment (negative slope), depletion (positive slope), or no correlation (zero slope) within a given length scale. For example, the PCCF curve for a simulated mixture of two species that are both enriched within 500 nm spherical domains, but randomly distributed within these domains (as might be expected from liquid-liquid phase separation) shows a clear increase at the cluster diameter, but then a plateau at shorter length scales. In contrast, the PCCF curve for a simulated mixture of two species that are both enriched within 500 nm spherical domains, but spatially partitioned within these domains (as might be expected from a diffusing transcription factor that predominantly exists in a non-chromatin-bound state) shows an increase at the cluster diameter and then a decrease at shorter length scales (**Fig. 3C-D, Fig. S21, Supplementary Note 1**). These simulations provide an intuitive relationship between the distributions of two molecular species and the shape of the PCCF plot, although we caution that the PCCF curve is not a one-to-one mapping and that alternative molecular distributions could also give rise to the same PCCF curves.

### Pair cross-correlation describes the length scale of associations between Exchange-PAINT targets and DNA

The full PCCF dataset for all thirteen labels consists of 169 pairwise interaction curves for each cell measured across two orders of magnitude in length ranging from 40 nm to 5 microns. To visualize this dataset, we averaged the PCCF curves to quantify the enrichment or depletion of inter-target interactions across two distinct length scales: 40 nm to 200 nm to assess nano-scale target-target interactions and 200 nm to 2000 nm to assess larger-scale changes in nuclear organization (**Fig. 4A, B**). These heatmaps showed that individual pairs of active (A-type) and repressive (B-type) targets showed a similar homotypic enrichment and a heterotypic partitioning as did the combined PCCF curves (**Fig. 3B**). These trends were strongest for short length scales (**Fig. 4A**) and decreased in strength at larger scales (**Fig. 4B**). To more fully explore these data, the PCCF curves for each cell were fit with a bi-linear model to report the short-length scale slope (slope 1), the long-length scale slope (slope 2), and the hinge point for each target-target pair (**Table S2-S4**). With these, we next investigated the PCCF curves between each Exchange-PAINT target and DNA (**Fig. 4C, D**). We observe a progressive transition for the PCCF curves where histone PTMs and proteins associated with constitutive and facultative heterochromatin formation are more strongly enriched with DNA and proteins associated with active transcription and RNA splicing are partitioned from DNA. As for the earlier results where we grouped proteins as corresponding to A-Type and B-Type chromatin, the enrichment/partitioning for each target with DNA began at approximately one micron and the trend became progressively stronger at shorter length scales. However, we also noted several exceptions to this trend. By nature of their incorporation into chromatin, all histone PTM’s were positively associated with DNA. Although activation-associated histone PTMs H3K4me1 and H3K27Ac were less enriched with DNA compared to repressive PTMs H3K9me3 and H3K27me3 in agreement with their localization to intermediate vs. dense chromatin regions respectively^18^. HP1α was enriched with DNA at length scales between 200 nm and two microns, consistent with its localization to heterochromatin^27^. However, below 200 nm, the slope of the HP1α - DNA PCCF curve was very close to zero (slope 1 = .0004 ± 0.0143, **Table S2 – S4**). This indicates that, at the scale of the nucleus, HP1α and DNA are co-enriched down to 200 nm, but below this scale, HP1α and DNA are randomly mixed. The lysine acetyltransferase p300 also showed a unique enrichment profile relative to DNA. From approximately 330 nm to two microns, p300 is enriched together with DNA. However, below 330 nm, there is a transition and the slope of the PCCF curves becomes positive (slope 1 = 0.0362 ± 0.0126, **Table S2 – S4**) which indicates a partitioning away from DNA at short length scales. Overall, these results demonstrate the ability of multiplexed single-molecule localization microscopy together with point statistics to reveal the spatial organization and scaling properties of different nuclear targets relative to chromatin. Importantly, the shape of these curves below the diffraction limit highlights unique organizational principles that could not be observed with conventional microscopes.

**Figure 4.**
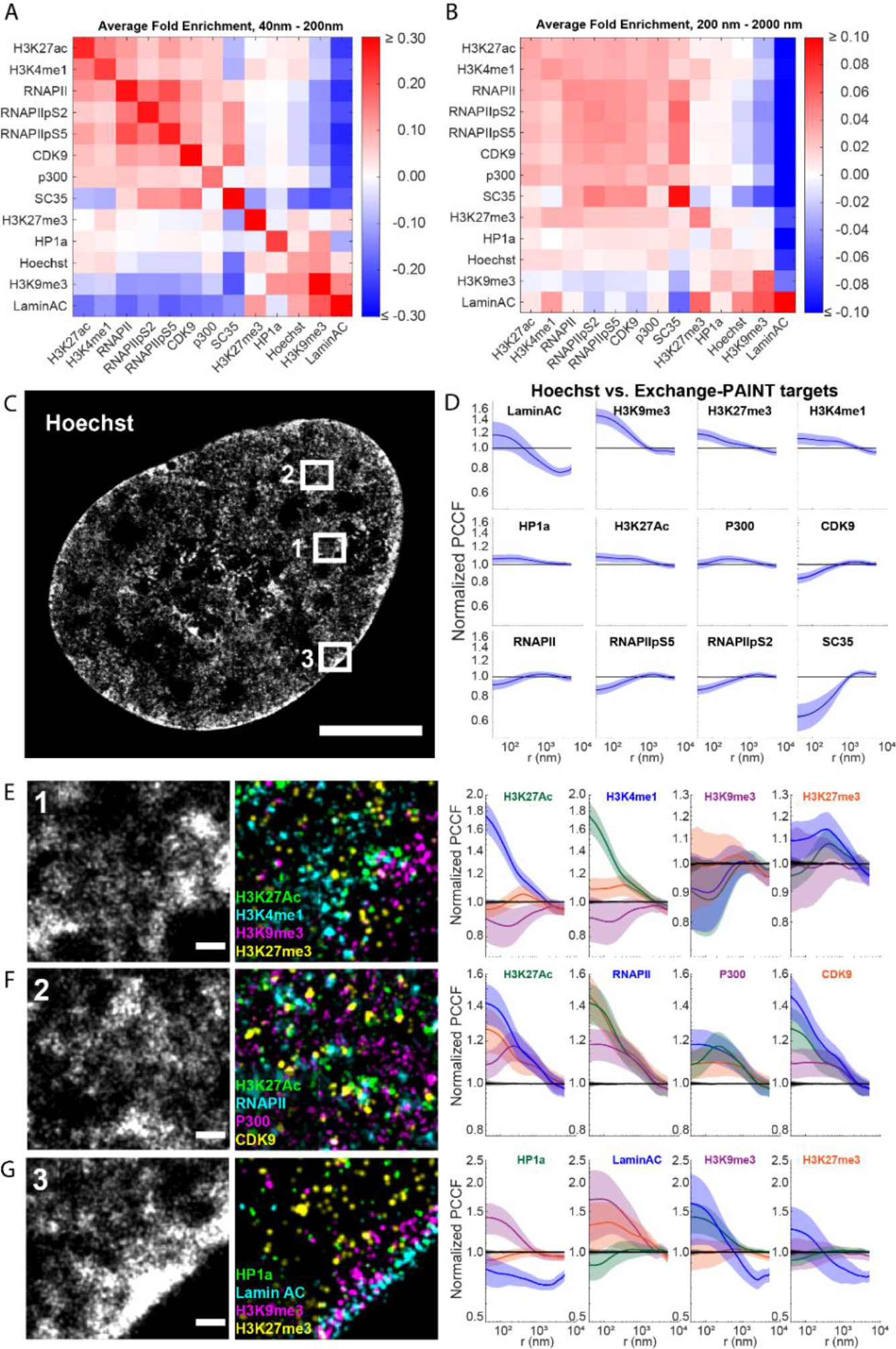
Quantification of Exchange-PAINT target distributions using the pair cross correlation function. (A, B) Heatmaps of the average fold enrichment computed from the normalized PCCF curves of each target-target pair over a radii range from 40 – 200 nm (A) and 200 – 2000 nm (B). Values are clipped at extrema to highlight variation. **(C)** Representative rendering of a Hoechst-labeled nucleus showing the region of interests for the renderings in E-G. **(D)** PCCF curves for Hoechst vs. Exchange-PAINT targets. (E-G) Representative renders and PCCF curves for histone PTM’s (E), transcription-associated targets (F), and heterochromatin-associated targets (G). Scale bars = 5 μm – (C), 200 nm – (E-G). Data from (D - G) are mean ± standard deviation. Data in (A-G) are from n = 31 cells across 3 independent biological replicates.

### Pairwise associations reveal the spatial organization of chromatin marks and transcriptional machinery

We next investigated the pairwise correlations between groups of proteins and histone PTMs with previously identified roles in gene regulation. Our antibody panel included four PTMs to the H3 histone tail that are known to be associated with: active gene enhancers and promoters (H3K27Ac)^28,29^ , active and primed enhancers (H3K4me1)^30^, poised enhancers and facultative heterochromatin (H3K27me3)^30–32^, and constitutive heterochromatin (H3K9me3)^33,34^ (**Fig. 4E**). Activation-associated H3K27Ac and H3K4me1 were positively correlated with each other and partitioned from H3K9me3 at all length scales. Interestingly, H3K27Ac and H3K4me1 are differently organized relative to H3K27me3. Both epigenetic marks were partitioned relative to H3K27me3 above two microns and enriched from two microns to approximately 500 nm. However, H3K27Ac and H3K27me3 become repartitioned below 400 nm (hinge = 413 ± 83 nm, slope 1 = 0.0518 ± 0.0201, **Table S2 – S4**), whereas H3K4me1 and H3K27me3 display no correlation (hinge = 433 ± 113 nm, slope 1 = 0.0139± 0.0190, **Table S2 – S4**). This reveals that while all three marks are enriched in the same approximate one-micron scale compartment, below 400 nm, poised enhancers are intermixed with facultative heterochromatin marks whereas active enhancers physically partition away from poised enhancers and facultative heterochromatin into discrete domains. This is reasonable considering that H3K27Ac and H3K27me3 are mutually exclusive PTMs on the same amino acid of histone H3. In contrast, H3K9me3 and H3K27me3 displayed no significant correlations (positive or negative) at any length scale, suggesting that facultative and constitutive heterochromatin domains are intermixed throughout the nucleus.

To study the organization of transcriptionally active nuclear regions, we next looked at the PCCF curves between H3K27Ac, RNAPII, p300, and CDK9 (**Fig. 4F**). H3K27Ac, RNAPII, and CDK9 were enriched at all length scales. In contrast, while p300 was enriched with the other three targets between ∼400 nm and one-micron, it lost its correlation with RNAPII (hinge = 474 ± 412 nm, slope 1 = -0.0138 ± 0.0145) and CDK9 (hinge = 575 ± 273 nm, slope 1 = 0.0067 ± 0.0176) below this length scale and became partitioned away from H3K27Ac (hinge = 242 ± 40 nm, slope 1 = 0.0555 ± 0.0243, **Table S2 – S4**). This is in general agreement with the p300-DNA PCCF curve and suggests that a portion of p300 in the nucleus is not directly interacting with either chromatin or the transcriptional machinery but is spatially partitioned below 300 nm.

Finally, we looked at the relationship between heterochromatin targets and the nuclear periphery (**Fig. 4G**). Like the trends for both targets relative to DNA, we found a strong correlation between HP1α and H3K9me3 that became enriched below one micron (hinge = 1177 ± 450 nm, slope 1 = - 0.1164 ± 0.0471, **Table S2 – S4**). H3K9me3 and H3K27me3 were enriched relative to lamin A/C but the correlation weakened at short length scales, suggesting that they do not directly interact with lamins (H3K9me3: hinge = 740 ± 878 nm, slope 1 = -0.0211 ± 0.1603, H3K27me3: hinge = 607 ± 863 nm, slope 1 = 0.0145 ± 0.0632, **Table S2 – S4**). In contrast, HP1α was neither enriched nor partitioned relative to lamin A/C at large length scales but became weakly partitioned below 300 nm suggesting that its concentration is slightly lower immediately adjacent to the nuclear membrane (hinge = 607 ± 669 nm, slope 1 = 0.0830 ± 0.0483, **Table S2 – S4**).

### Transcriptional machinery concentrates at the periphery of and within SC-35 splicing speckles

Finally, we sought to understand the spatial distribution of targets relative to splicing speckles marked by the protein SC-35. These micron-scale nuclear domains are thought to act as storage depots for splicing components^35^ and hubs to facilitate efficient mRNA splicing of nearby genes which are typically highly transcribed^11,36,37^ . Given this, we explored the spatial relationship between SC-35 and transcription-related targets (**Fig. 5A**). We found that SC-35 was positively enriched with RNAPII and its phosphorylated variates from approximately 400 nm up to two microns, but that below this level, there was no enrichment (**Fig. 5B**, RNAPII: hinge = 574 ± 167 nm, slope 1 = 0.0118 ± 0.0386, RNAPIIpS5: hinge = 383 ± 546 nm, slope 1 = -0.0134 ± 0.0263, RNAPIIpS2: hinge = 361 ± 256 nm, slope 1 = -0.0123 ± 0.0359, **Table S2 – S4**). This suggests that these targets are generally enriched within splicing speckles, but do not associate closely with SC- 35 in these regions. Interestingly, we found that phospho-specific RNAPII targets were more enriched with SC-35 than non-phospho-specific RNAPII, which is consistent with previous studies showing that the phosphorylation state of the RNAPII CTD can drive a switch between its localization to transcriptional vs. splicing condensates^38^. In contrast, SC-35 and CDK9 were positively correlated at all length scales suggesting that these two proteins closely associate (hinge = 1009 ± 811 nm, slope 1 = -0.1039 ± 0.0517, **Table S2 – S4**). This agrees with prior studies showing that SC-35 and CDK9 co-immunoprecipitate and that there is a secondary role for SC-35 of promoting RNAPII pause release and elongation^39^ . Both p300 and H3K27Ac showed a small enrichment with SC-35 at approximately one micron but became depleted below this length scale with H3K27Ac more strongly depleted than p300, consistent with splicing speckles being depleted of chromatin (p300: hinge = 979 ± 385 nm, slope 1 = 0.0238 ± 0.0296, H3K27Ac: hinge = 686 ± 211 nm, slope 1 = 0.1404 ± 0.0287, **Table S2 – S4**).

**Figure 5.**
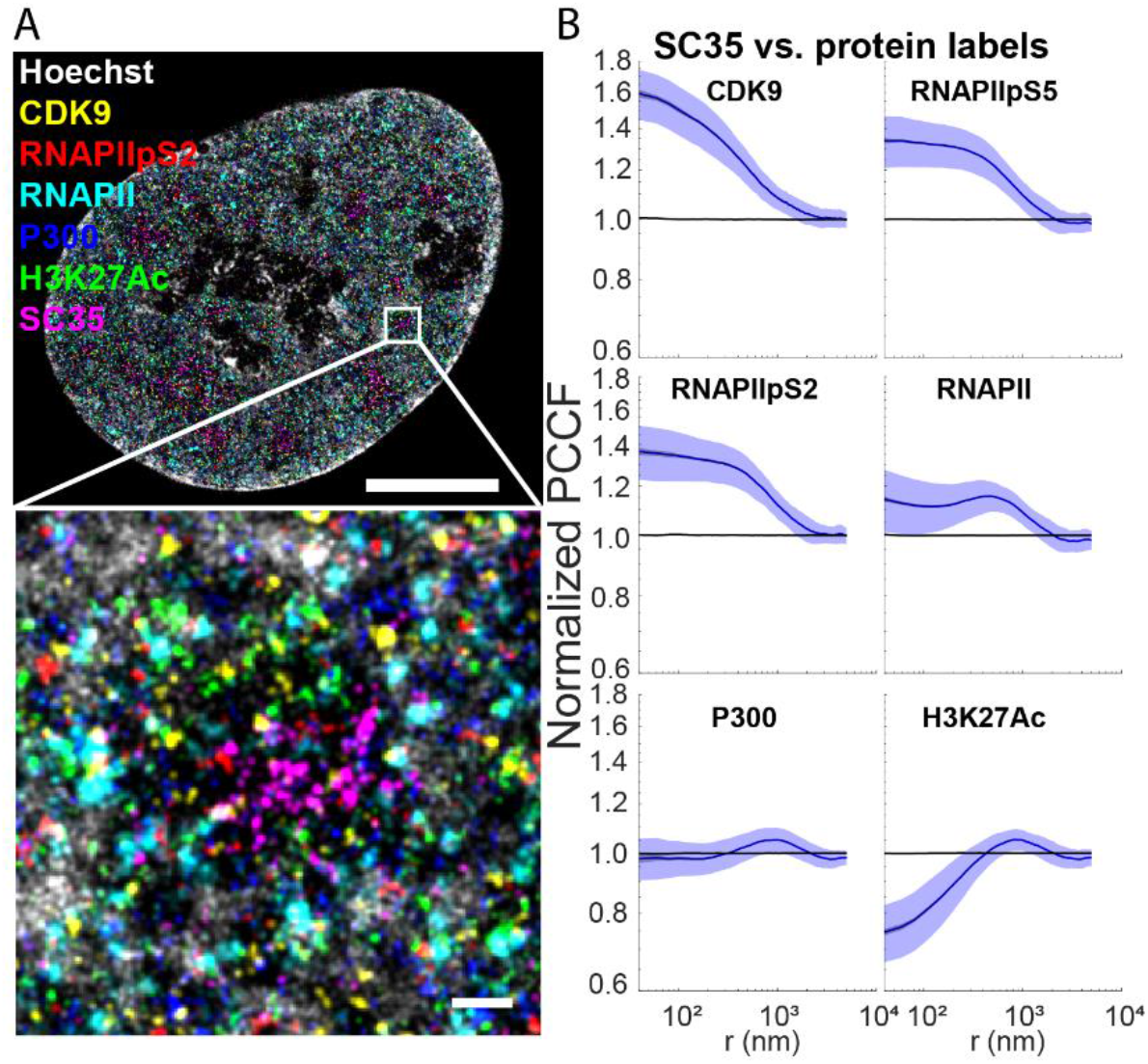
Multiplexed imaging and quantification of SC-35 speckles. (A) Representative rendering and zoomed inset of speckle-associated Exchange-PAINT targets. **(B)** PCCF curves for SC-35 vs. select Exchange-PAINT targets. Scale bars = 5 μm – (A), 200 nm – (zoom). Data from (B) are mean ± standard deviation and are from n = 31 cells across 3 independent biological replicates.

Overall, these results reveal the spatial organization of DNA, proteins and epigenetic marks in the nucleus of single cells from the nano to the microscale. Using Exchange-PAINT and high sensitivity imaging, our precision is comparable to the ∼11 nm diameter of a single nucleosome^40^, which allows for the precise localization of specific epigenetic marks and transcriptional machinery. We found that most of the targets we imaged display increased self-association below ∼200 nm which could be explained by protein oligomerization or homotypic clustering of epigenetic chromatin marks. By comparing pairwise localizations between different groups of targets, we find that the local microenvironment of the nucleus becomes functionally specified below approximately one micron. Above this scale, a given region of the nucleus is equally likely to contain marks pertaining to both active euchromatin and repressed heterochromatin. Understanding the mechanisms that shape the 3D genome organization is an active and robust area of research^2,3^ and several organizational principles have been proposed including polymer models^41,42^, histone tail charge-charge interactions^43–45^ and phase separation and condensate formation^46–49^. Active processes such as loop extrusion^50,51^ and transcription^52–55^ also locally perturb chromatin, leading to long-term structural changes. Given this, we next sought to understand how perturbations to epigenetic marks and nuclear processes might impact the above results.

### Histone hyperactylation reduces nuclear compartmentalization and decreases the molecular differences between active and repressive nuclear environments

We first induced chromatin hyperacetylation using the broad-spectrum histone deacetylase inhibitor trichostatin A (TSA). TSA treatment increased and dispersed histone acetylation throughout the nucleus, resulting in a nearly ten-fold increase in H3K27Ac levels (**Fig. 6A, B - inset**) while minimally perturbing the localization density of other Exchange-PAINT targets **(Fig. S22**). G(r) curves for H3K27Ac under control and TSA treated conditions revealed that TSA reduced H3K27Ac clustering at short length scales in agreement with smaller and more uniformly dispersed H3K27Ac foci in the images (**Fig. 6B**). This suggests that, in addition to reinforcing existing acetylated domains, TSA treatment leads to hyperacetylation throughout the nucleus^56–58^; however, it may also lead to counterintuitive effects such as hypoacetylation of some gene regulatory elements, such as promoters^58,59^. Concordant with the increased histone acetylation, and in agreement with prior literature^43,60^, TSA treatment also decondensed DNA (**Fig. 6C**). Given these results, we next sought to understand how chromatin hyperacetylation changed the pairwise interactions between other Exchange-PAINT targets (**Figures S23-S38**).

**Figure 6.**
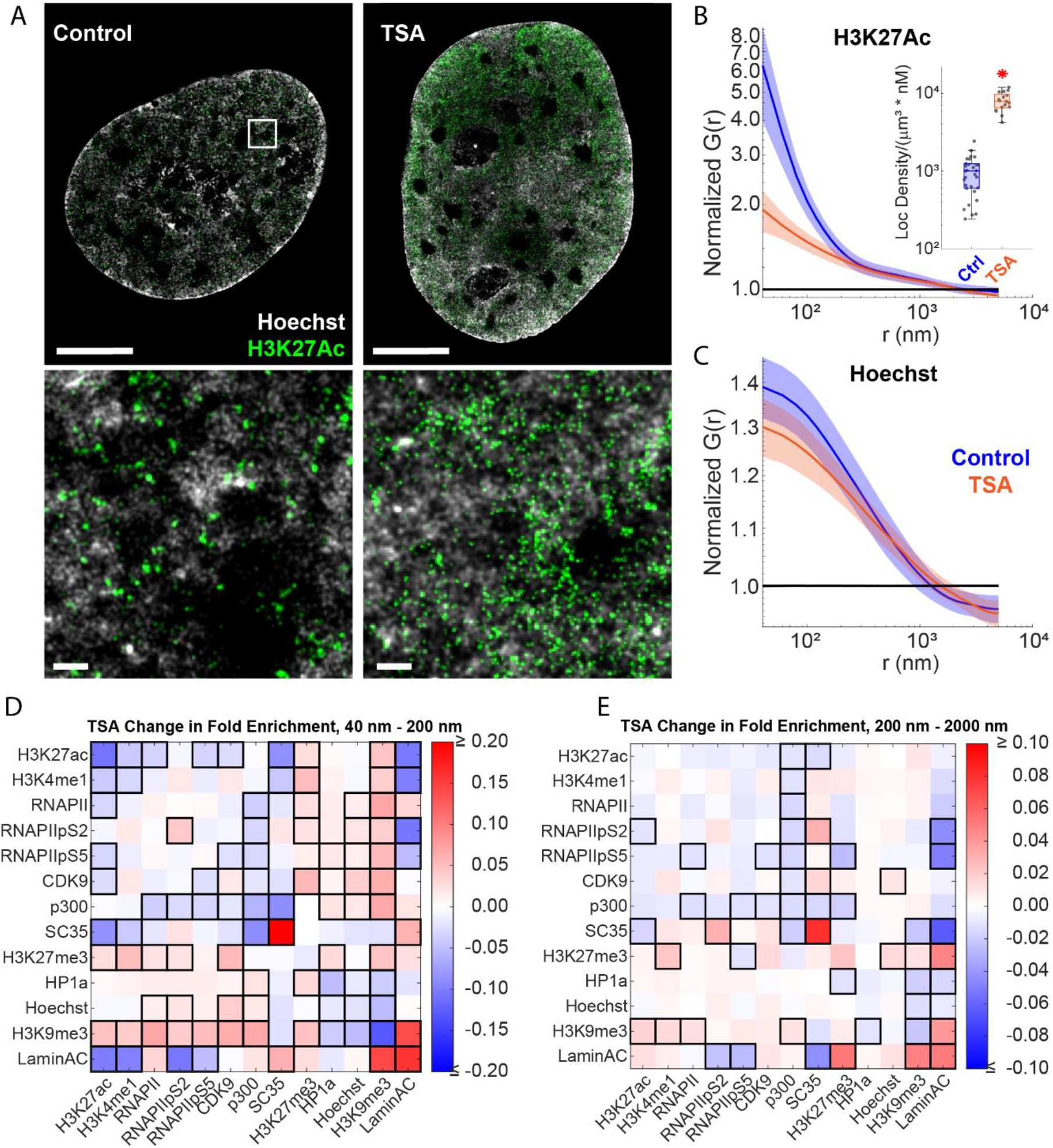
Quantifying the effects of histone deacetylase inhibition on nuclear organization. **(A)** Representative renderings and zoomed insets of control and TSA treated nuclei showing Hoechst and H3K27Ac. **(B)** Pair correlation function quantification of H3K27Ac self-association under control (blue) and TSA (orange) treated conditions. Inset shows a boxplot of the localization density normalized by the imager strand concentration for each condition. **(C)** Pair correlation function quantification of Hoechst self-association under control (blue) and TSA (orange) treated conditions. **(D, E)** Heatmap of the difference between control and TSA conditions on the average fold enrichment for each target-target pair over a radii range from 40–200 nm (D) and 200 – 2000 nm (E). Values are clipped at extrema to highlight variation. Black outlines represent comparisons where p <= 0.05 by Student’s T-test. Scale bars = 5 μm – (A), 200 nm – (zoom). Data from (B, C) are mean ± standard deviation. Data from (B-E) are from n = 31 cells across 3 independent biological replicates for control and n = 17 cells across 2 independent biological replicates for TSA.

To quantify perturbations to target pair associations in response to drug perturbation, we computed the difference in fold enrichment relative to control over two length scale regimes. Compared to control cells, TSA did not change the overall grouping of homotypic clustering between euchromatin and heterochromatin-associated targets, but it did reduce the strength of association or depletion for most target pairs, resulting in a more intermixed and randomly distributed nucleus (**Fig. 6D, E, Figs. S23-38**). TSA-induced hyperacetylation altered the transcriptionally active nuclear compartment by reducing the tight spatial enrichment of active enhancers marked by H3K27Ac and transcription associated machinery RNAPII, RNAPIIpS5 and CDK9, and the co-occurrence of H3K27Ac and H3K4me1, especially at short length scales (**Fig. 6D**). It also disrupted the co-association between p300 and CDK9, RNAPII, and SC-35. Furthermore, TSA reduced the normal partitioning of H3K27Ac and heterochromatin marks H3K27me3 and H3K9me3 while increasing the partitioning of H3K27Ac from splicing speckles (SC-35) and the nuclear periphery (lamin A/C). TSA also altered heterochromatin in the nucleus. It transitioned facultative heterochromatin marked by H3K27me3 into a signature more similar to active euchromatin by increasing its enrichment for H3K4me1, p300, and RNAPII (**Fig. 6D, E, S28**). Finally, TSA altered constitutive heterochromatin by decreasing the enrichment between H3K9me3 and HP1α and enriching H3K9me3 with transcription machinery (p300, RNAPII, and CDK9) (**Fig. 6D,E, Fig. S24**). Interestingly, after TSA treatment, constitutive heterochromatin marked by H3K9me3 became enriched with H3K27me3 and both H3K9me3 and H3K27me3 became more tightly correlated with the nuclear lamina (**Fig. 6D, E, Figs. S26, S28**). This data agrees with previous biochemical assays showing that H3K27me3 is a promiscuous chromatin mark that can spread to constitutive heterochromatin sites upon reduction of H3K9me3^61,62^ and also suggests that heterochromatin at lamin-associated domains may be more resistant to the changes induced by chromatin hyperacetylation.

### Inhibition of the histone acetyltransferase p300 disrupts the localization of transcriptional machinery relative to epigenetic marks

We next explored the effects of reducing histone acetylation in active genomic regions using targeted catalytic inhibition of p300 with A-485. A-485 treatment caused an approximate ten-fold drop in H3K27ac levels, with smaller reductions in the densities of other marks including HP1α, RNAPII, CDK9, and H3K9me3 (**Fig. 7A, B – inset, Fig. S22**). Residual H3K27ac marks appeared as dispersed high-intensity puncta corresponding to increased self-association at short length scales in the G(r) curve and lower enrichment at larger length scales (∼150 nm, **Fig. 7A, B**). Reduced H3K27ac also led to a small increase in chromatin condensation, especially at smaller length scales below ∼200 nm (**Fig. 7C**). In contrast to TSA, which had significant effects on both heterochromatin and euchromatin target interactions across both short and large length scales, the effects of A485 were much more limited (**Fig. 7D, E, Fig. S21, Figs. S39-S53**). Residual H3K27Ac marks lost their enrichment for H3K4me1, as well as with transcription-associated targets RNAPII, its phosphorylated variants and CDK9 (**Fig 7D, S42**). H3K27Ac also showed reduced enrichment for both p300 and SC-35 at larger length scales (**Fig. 7E, Fig. S42**), but remained co-associated and even more enriched relative to control cells below 100 nm (**Fig. 7D, Fig. S42**) suggesting that there is lower turnover of histone H3K27 acetylation in proximity to nuclear speckles. Reducing H3K27ac levels also reduced transcription-associated targets (RNAPII, RNAPIIpS5, CDK9) in the proximity of H3K4me1 at short length scales ≲200 nm (**Fig. 7D**). This effect was especially apparent in terms of transcriptional pause-release, as indicated by the drop in pairwise correlations between H3K4me1 with RNAPII pS5 and CDK9 (**Fig. S40**). However, these changes were not maintained at length scales greater than 200 nm (**Fig. 7E**), suggesting that A-485 disrupts the close associations within transcriptionally-active regions without altering the overall partitioning between euchromatin and heterochromatin in the nucleus. Finally, A485 treatment also caused an enrichment between lamin A/C and transcriptionally active targets relative to control cells (**Fig. 7D, E, Fig. S46**), as well as between H3K27me3 and SC-35 (**Fig. 7D, E, Fig. S43**).

**Figure 7.**
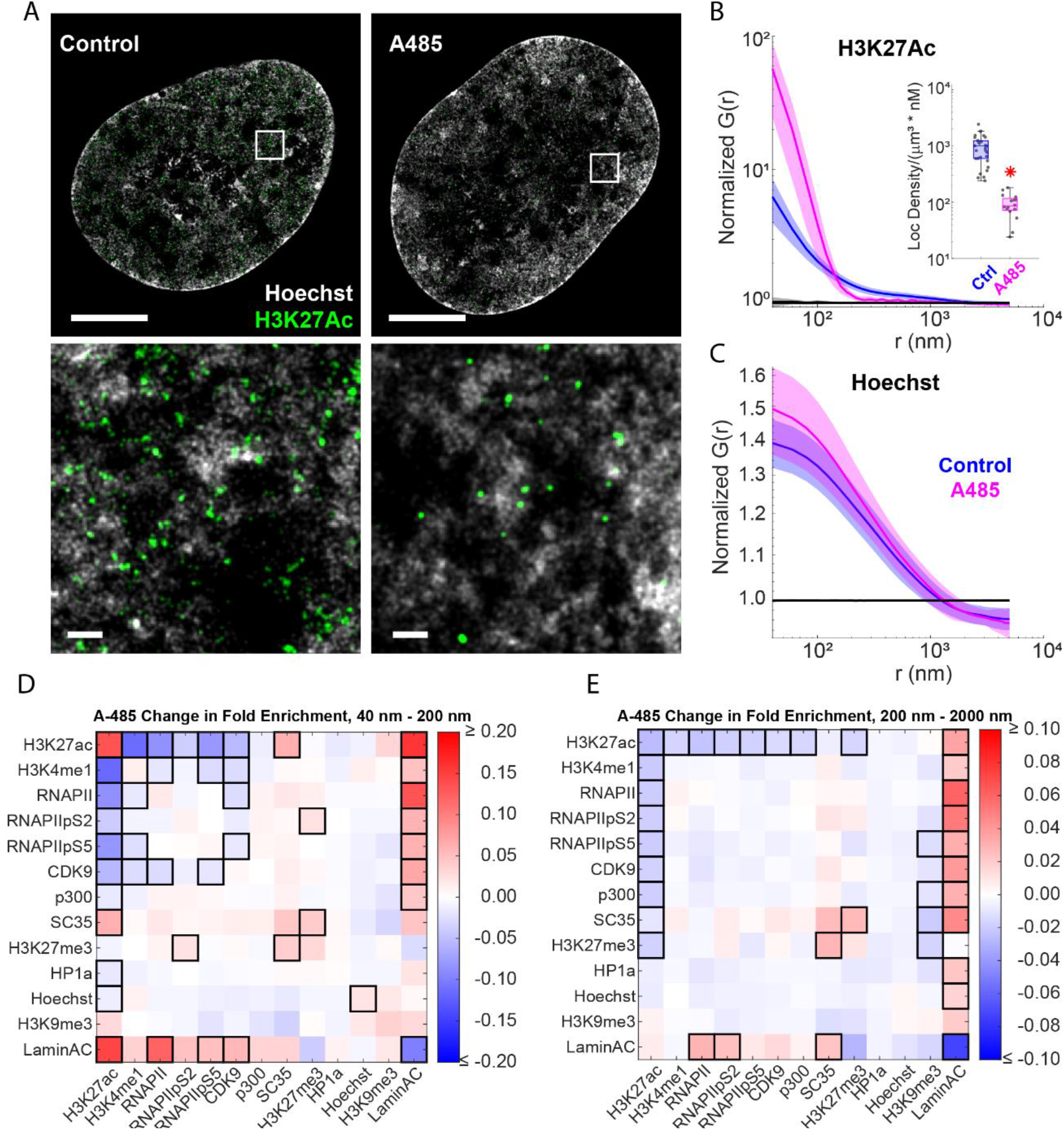
Quantifying the effects of p300 catalytic inhibition on nuclear organization. **(A)** Representative renderings and zoomed insets of control and A485 treated nuclei showing Hoechst and H3K27Ac. **(B)** Pair correlation function quantification of H3K27Ac self-association under control (blue) and A485 (magenta) treated conditions. Inset shows a boxplot of the localization density normalized by the imager strand concentration for each condition. **(C)** Pair correlation function quantification of Hoechst self-association under control (blue) and A485 (magenta) treated conditions. **(D, E)** Heatmap of the difference between control and A485 conditions on the average fold enrichment for each target-target pair over a radii range from 40 – 200 nm (D) and 200 – 2000 nm (E). Values are clipped at extrema to highlight variation. Black outlines represent comparisons where p <= 0.05 by Student’s T-test. Scale bars = 5 μm – (A), 200 nm – (zoom). Data from (B, C) are mean ± standard deviation. Data from (B-E) are from n = 31 cells across 3 independent biological replicates for control and n = 16 cells across 2 independent biological replicates for A485.

### Transcription inhibition disrupts protein-protein interactions within euchromatin and strengthens heterochromatin compartmentalization

As a final perturbation, we investigated the effects of transcriptional inhibition. Treatment of cells with α-amanitin (αAM) leads to ubiquitination and degradation of RPB1, the major subunit of RNAPII^63^. αAM treatment reduced the levels of RNAPII and its phosphorylated variants to approximately 40% of unperturbed levels (**Fig. 8A, B**). This treatment also caused a dramatic reduction in the levels of several other targets including H3K27ac, CDK9, p300, and lamin A/C (**Fig. S22**). Similar to the effects of A-485 on H3K27Ac, residual RNAPII localizations showed small clusters with increased self-association at short length scales (**Fig. 8A**). α AM treatment also led to a dramatic increase in the DNA condensation below one micron (**Fig. 8C**). Heatmaps of the integrated fold enrichment and PCCF plots for target interactions revealed that the organization of the euchromatic nuclear environment was completely disrupted with reduced enrichment between all transcription-associated targets and histone PTMs (**Fig. 8 D, E, Figs. S54 - S68**). Residual localizations for active targets including H3K27Ac, RNAPII, CDK9, and p300 were organized in small clusters that all had reduced inter-target enrichment compared to control cells. These targets also had broadly reduced proximity to both active and repressive chromatin marks and DNA and instead became enriched within SC-35 positive speckles and near the nuclear periphery (**Figs. S61, S68**). H3K27Ac showed increased self-association at short length scales and decreased self-association at large length scales whereas H3K4me1 was unaffected. Finally, both facultative (H3K27me3) and constitutive (H3K9me3) heterochromatin marks showed increased enrichment with each other and with DNA at all length scales (**Fig. 8 D, E, Figs. S56, S58**).

**Figure 8.**
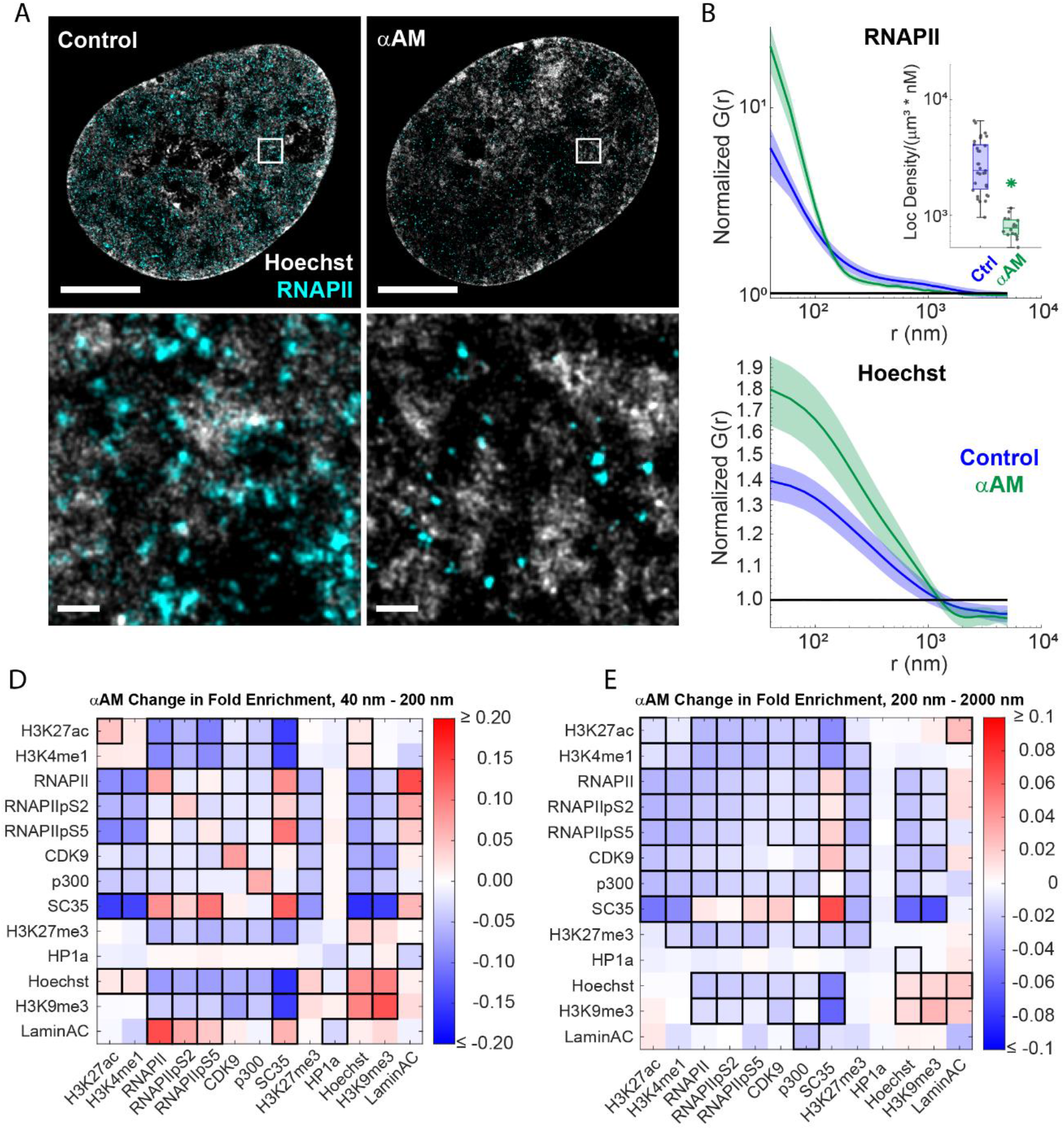
Quantifying the effects of transcription inhibition on nuclear organization. **(A)** Representative renderings and zoomed insets of control and αAM treated nuclei showing Hoechst and RNAPII. **(B)** Pair correlation function quantification of RNAPII self-association under control (blue) and αAM (green) treated conditions. Inset shows a boxplot of the localization density normalized by the imager strand concentration for each condition. **(C)** Pair correlation function quantification of Hoechst self-association under control (blue) and αAM (green) treated conditions. **(D, E)** Heatmap of the difference between control and αAM conditions on the average fold enrichment for each target-target pair over a radii range from 40 – 200 nm (D) and 200 – 2000 nm (E). Values are clipped at extrema to highlight variation. Black outlines represent comparisons where p <= 0.05 by Student’s T-test. Scale bars = 5 μm – (A), 200 nm – (zoom). Data from (B, C) are mean ± standard deviation. Data from (B-E) are from n = 31 cells across 3 independent biological replicates for control and n = 16 cells across 2 independent biological replicates for αAM.

## Discussion

We describe the application of multiplexed single molecule Exchange-PAINT together with HIST illumination microscopy to study the spatial distributions of DNA, histone PTMs and nuclear proteins, in 3D, from the nano to the microscale. By revealing the organization of these labels at the nanoscale, this work complements prior studies that used chromosome tracing and immunofluorescence imaging to study how specific genetic sequences are organized relative to larger subnuclear structures and chromatin marks via diffraction limited imaging^10,11^. It builds upon prior super-resolution imaging studies of the cell nucleus^19,43^ by improving the spatial resolution, increasing the field of view, and extending multiplexing to image many different targets together. Overall, we find that the G(r) self-association curves for most nuclear targets display two power-law scaling regimes suggestive of a fractal-like organization. The lower regime spans from our localization precision up to approximately 200 nm. Above this length scale, the distributions became closer to random. This contrasts with a view of a compartmentalized nucleus with clear delineating boundaries between different regions. Given that our targets are either directly associated with or bind to chromatin, we envision that most of this large-scale organization arises from the fractal-like distribution of chromatin itself^23,42^ which forms a viscoelastic gel^49^ that templates histone PTMs and shapes the pore space for diffusing nuclear proteins. The results from drug perturbation experiments suggest that changes to histone PTMs and transcriptional inhibition feedback to change the strength and fractal dimension of chromatin, but do not fundamentally change its physical structure.

Within this context, our pairwise correlation measurements between different proteins complement prior super-resolution or multiplexed studies of the nucleus, particularly the concepts of homotypic clustering^64,65^ or partial demixing of type A/B nuclear compartments^11,50^, zonation of gene activation- or repression-associated targets^10^, and the association of splicing speckle edges with high transcriptional activity^10,36,66^. Using a multiplexed approach, we build upon these prior studies to better define the length scale of homotypic clustering and the pairwise distances of gene activation- and repression-associated targets down to the nanoscale. Importantly, we describe how the shape of the PCCF curve relates to and informs biological mechanisms. For example, the PCCF curves for DNA vs. HP1α and DNA vs. p300 both show preferential enrichment of DNA with each protein from 2 microns down to approximately 200 nm. However below 200 nm, the PCCF curves plateau (for HP1α) or decrease (for p300) indicating these proteins lose their spatial correlation with DNA or become partitioned respectively. These results reinforce prior observations that HP1α is a chromatin crosslinking protein that is enriched within, but can also freely diffuse though heterochromatin regions ^67–69^ and that p300 can coalesce into kinetically less active condensate domains that become compartmentalized from chromatin^70^. Further, our results also describe the organization of active transcriptional environments, revealing that active enhancers (H3K27Ac + / H3K4me1 +) and poised enhancers (H3K4me1 + / H3K27me3 +) are both enriched at ∼500 nm, but that active enhancers partition away from poised enhancers below 200 nm.

For all drug perturbations tested, we found that, at the micron length scale, the nucleus is surprisingly robust to changes in histone acetylation, but that these perturbations have more pronounced effects on the spatial organization below 200 nm. This may represent the roles of larger nuclear features such as the nucleolus and the nuclear lamina which are less impacted by these perturbations. Histone hyperacetylation by TSA reduced the partitioning between active and repressive targets. It decreased the co-enrichment of activation-associated targets and redistributed them to traditionally heterochromatic regions which is in agreement with prior studies demonstrating that TSA leads to both up- and downregulated gene expression ^71,72^.

In contrast, catalytic inhibition of p300 by A-485 reduced the co-enrichment of activation-associated targets in euchromatin and instead shifted them closer to the nuclear lamina. A-485 treatment has been shown to reduce transcription by preventing the binding of pre-initiation complex components TFIID and TATA-box-binding protein (TBP) and thus decrease the recruitment of RNAPII and its phosphorylated variants to chromatin, particularly at enhancers and super-enhancer regions^73,74^. Together with our data, this supports the role of p300 acetyltransferase activity in localizing the transcriptional machinery to actively transcribed regions; although, we cannot currently distinguish whether this is due to the reduction in chromatin acetylation or its effects on other substrates.

Transcription inhibition with αAM increased the partitioning between euchromatin and heterochromatin and led to DNA condensation. Activation-associated markers became further partitioned from DNA and concentrated in splicing speckles while repression-associated targets became more closely associated with DNA. Interestingly, CDK9 and p300 showed reduced association with DNA, suggesting a decrease in binding due to RNAPII degradation. This is in agreement with the role of both in transcriptional elongation^74,75^. Furthermore, increased DNA and heterochromatin self-association supports a role for active transcription in resisting DNA compaction^54^. Surprisingly though, although we observe changes in the self-association curves for individual histone PTM’s, we do not see changes in the relative organization between activation-associated or repression-associated histone PTM’s after treatment with αAM. Transcription can influence chromatin structure by relocalizing cohesin^76,77^ and stabilizing nucleosomes in transcriptionally active regions^60,78,79^, potentially through phase separation^80^. Our results reveal that although transcription inhibition with αAM alters the local compaction state of both heterochromatin and euchromatin, it does not alter the spatial separation between these two environments.

Like all antibody-based methods, our approach is dependent on highly efficient and specific antibodies toward each target. Whenever possible, we selected knockout-validated antibodies that had been confirmed to have worked for immunofluorescence. When not possible, we tried to select antibodies that had previously been documented and validated via consortiums such as the Encyclopedia of DNA Elements (ENCODE)^81^. Nevertheless, only roughly 46% of the antibodies we tested proved suitable for imaging (**Table S5**). In addition to antibody specificity, errors in our measurements could arise from non-specific binding of the imager strands to DNA or insufficient washing during probe exchanges. Although we found this to be relatively low, typically between 1 and 10% for non-specific imager strand binding, and less than 10% residual localizations after washing (**Figs. S1-S3**) these measurements should be confirmed for any new probe designs. Fortunately, the pair correlation function is relatively insensitive to varying the number of target localizations for a given species (**Fig. S21**), allowing for quantitative comparison of localization distances utilizing antibodies of varying efficiency. Nonspecific binding events will serve to drive the correlation curves closer to a random distribution. This would reduce our sensitivity to detect organizational changes but would be unlikely to introduce artifactual correlations between the targets. When aggregating labels for analysis of A- or B-type-associated targets, average values will be disproportionately weighted by more abundant targets. Finally, pair correlation functions represent the average spatial enrichment or depletion between two species at a given length scale. This makes them especially useful to capture trends under incomplete or unknown labeling efficiency but precludes the use of these curves for describing the characteristics of a specific region within the nucleus of a given cell.

Nevertheless, we believe that the methods and computational approach demonstrated here will be a useful addition to understand nuclear organization in addition to other cellular processes. In contrast to co-immunoprecipitation, each PCCF curve reports the spatial co-localization between pairs of targets from 40 nm to 2 microns, thus revealing organizational principals that could not be detected with diffraction limited imaging or by studying direct binding via biochemical crosslinking methods alone. Because the number of pairwise interactions scales as the square of the number of labels, our 13-plex dataset produced 169 pairwise curves from a single experiment, greatly exceeding what could be done with individual pull downs. Combining our approach together with automated fluid exchange, recent improvements to DNA PAINT that utilize transient adaptor strands^82,83^, and instruments that afford even higher precision localization^84,85^ will increase throughput and resolution, decrease cost, and could potentially pave the way for diagnostic or translational applications^86–88^.

## Materials and Methods

### 1. Cell culture

The human telomerase-immortalized hTERT RPE-1 cell line (ATCC CRL-4000) was cultured with complete growth medium consisting of DMEM/F-12 (Gibco 11320033) supplemented with fetal bovine serum to 10% and penicillin/streptomycin to 1%. Cells were cultured in 5% CO2 at 37°C. For non-perturbed, A-485, and α-amanitin conditions, cells were plated at an approximate density of 6 x 10^4^ cells/cm^2^, and for the TSA condition, cells were plated at an approximate density of 3 x 10^4^ cells/cm^2^. Cells were grown overnight in 5% CO2 at 37°C on #1.5 glass – either on 24-well glass bottom plates (Cellvis P24-1.5H-N) for DNA- and Exchange-PAINT experiments or 12 mm round coverslips (Warner Instruments CS-12R15, cat. 64-0712) for diffraction-limited imaging. Drug dosage was optimized using widefield quantification of upregulated H3K27ac for TSA-induced hyperacetylation, downregulated H3K27ac for A-485-inhibited p300 activity, and decreased RNAPII signal with αAM-induced RNAPII inhibition and degradation (**Fig. S69**). For the A-485 Exchange-PAINT condition, cells were treated for two hours in 10 μM A-485 (Selleck S8740) immediately prior to fixation. For the TSA Exchange-PAINT condition, cells were treated for 20 hours in 500 nM TSA (Sigma T8552-1MG) immediately prior to fixation. For the α-amanitin Exchange-PAINT condition, cells were treated for 2 hours in 100 μg/mL α-amanitin (Sigma A2263) prior to fixation.

### 2. Fixation, permeabilization, and blocking

Cells were fixed with 4% paraformaldehyde (in 1x PBS, diluted from 16% paraformaldehyde, Electron Microscopy Sciences 15710) for 20 minutes at room temperature and quenched with 100 mM ammonium chloride (Sigma A9434-500G) in 1x PBS (Corning 46-013-CM) for 10 minutes at room temperature. This was followed by three five-minute washes in 1x PBS on a rocker. Cells were permeabilized in 0.2% Triton X-100 (VWR 97062-208) for 20 minutes and then washed again, three times for five minutes in 1x PBS. Cells were then blocked for a minimum of overnight and up to three days at 4°C in a blocking buffer of filtered 3% bovine serum albumin (Sigma A9418- 50G), 0.02% Tween-20 (Thermo 28320), 1 mM EDTA (Sigma E6758-100G or Thermo J60893.K2), and 1x PBS.

### 3. Immunofluorescence staining

#### 3.1. Widefield imaging sample preparation

Stains with the FluoTag series (FluoTag®-X2 anti-Mouse IgG Kappa Light Chain, NanoTag N1202, and FluoTag®-X2 anti-Rabbit IgG, NanoTag N2402) were prepared sequentially for primary antibody concentration optimization – samples were incubated for one hour with a primary dilution in the blocking buffer, three five-minute washes with PBS, and then one hour incubation with a 200x secondary antibody dilution in blocking buffer. For representative widefield images at selected primary concentrations, FluoTag secondaries and selected primary dilutions were diluted and pre-associated in blocking buffer at a 2.5 molar ratio for one hour at room temperature. The mixed primary and secondary solutions were then incubated with the sample for one hour at room temperature and the sample washed three times for five minutes with 1x PBS. For DAPI (Invitrogen D1306) staining of DNA when applicable, samples were additionally incubated in 1.25 μg/mL DAPI dilutions in 1x PBS for 10 minutes at room temperature on a rocker, then washed again in PBS. Samples plated on coverslips for antibody concentration optimization were then mounted in ProLong Gold Antifade Mountant (Invitrogen P10144) on glass slides (Fisher 125444) and sealed with clear nail polish. Samples were stored for up to a week at 4°C in PBS.

Primary antibodies and concentrations used in subsequent Exchange-PAINT experiments are listed in **Table S1**. All tested antibodies and screening results are listed in **Table S5**.

#### 3.2. Exchange-PAINT sample preparation

Exchange-PAINT secondary antibodies were custom prepared by Massive Photonics for orthogonality and low background in a nuclear environment with oligonucleotide sequences listed in Table S1.

Primary antibodies and Exchange-PAINT secondary antibodies (P1, P3-5, P8, P9, P13, P16, P17, P19, P20, and P24) were diluted and pre-associated in blocking buffer at a 2.5 molar ratio, based on pre-selected antibody dilutions from FluoTag-based optimization and a stock concentration of 5 μM for the Exchange-PAINT probes, and brought to 5x volume by blocking buffer, for one hour at room temperature. For the last 10 minutes of preincubation of the primary and secondary antibodies, NanoTag multiplexing blocker (for mouse host primaries, catalog number K0102-50, and for rabbit host primaries, catalog number K0202-50) was added at a volume to weight ratio of 10 μL/μg primary to bind remaining unbound primary antibodies and minimize antibody hopping during multiplexed labeling. The mixed primary, secondary, and multiplexing blocker solutions were then combined to a total volume of 250 μL with blocking buffer per sample well of a 24-well glass bottom plate and incubated with the sample for one hour at room temperature. The sample was washed twice to remove unbound antibody for five minutes with 1x PBS. A 750x v/v dilution of wheat germ agglutinin-conjugated fluorescent nanodiamonds from Adámas Nanotechnologies (NDNV100nm-WGA) in 1x PBS was sonicated for 20 minutes and then added to the sample for 10 minutes on a rocker to adhere point- and broad spectrum-emitting, non-bleaching fiducial markers to cell membranes. The sample was then washed twice for five minutes in 1x PBS. The sample was post-fixed in 4% PFA for 20 minutes, to covalently bind multiplexed antibody and wheat germ agglutinin labels to the sample and each other, and fixation quenched by 100 mM ammonium chloride for 10 minutes. Samples were stored for up to a week at 4°C until imaging. Empty wells on glass-bottom plates containing each Exchange-PAINT sample were prepared for calibration of imaging settings at room temperature. They were coated in 0.2% weight/volume poly-L-lysine (Sigma P2636-500MG) diluted in deionized water (diH2O) for 20 minutes, washed three times for five minutes in diH2O, coated in 100nm TetraSpeck beads (Sigma T7279) of varying dilutions (500x-5000x) in diH2O, and washed again in diH2O three times for five minutes each. Prior to coating, TetraSpeck beads were sonicated for 20 minutes.

Targets were imaged in pairs such that one target each was paired with one imager fluorophore (ATTO655 or Cy3B) and bound fluorophores were in a density range appropriate for single molecule localization. Before imaging, the first pair of Exchange-PAINT imager probes were diluted in imaging buffer (1x PBS, 637 mM NaCl, 1 mM EDTA, pH 7.5) and added to the sample. The sample plate was covered with a custom lid modified to have metallic inlet and outlet ports through blunt end 20g needle tips (SAI B20-150). The tips were adhered with Krazy Glue through the plastic plate lid and sealed with parafilm, connected with plastic tubing external to the lid to interface with syringes. The plate was then hot glued to a multi-well plate holder for the Nikon Eclipse Ti2 inverted microscope and allowed to cure for 20 minutes at room temperature. Working imager strand concentrations are listed in **Table S1**.

### 4. Widefield and Highly Inclined Swept Illumination (HIST) microscopy

All imaging was performed on a Nikon Eclipse Ti2 inverted microscope with separate optical paths for widefield and HIST microscopy. The widefield path was illuminated with a Lumencor Spectra III Light Engine with 390, 440, 475, 510, 555, 575, 635, and 740 nm excitation bands and imaged onto an Orca Flash 4.0 sCMOS camera (Hamamatsu C13440-20CU). The HIST path was illuminated via 560 nm (2RU-VFL-P-2000-560-B1R) and 642 nm (2RU-VFL-P-2000-642-B1R) emission lasers from MPB Communications Inc through a custom light path as described below and imaged onto a pair of Orca Fusion BT sCMOS cameras (Hamamatsu C15440-20UP). All images were acquired using a 60x 1.27 NA water immersion objective (Nikon MRY10060).

#### 4.1. Widefield microscopy and analysis

FluoTag 488 or 647 fluorescent nanobodies were imaged with 475 and 635 nm excitation, respectively. DNA stains (DAPI) were imaged with 390 nm excitation. Acquired images were analyzed in MATLAB and FIJI^89^ after import of ND2 files through BioFormats^90^. For quantification of relative FluoTag signal across conditions (e.g., in drug concentration optimization), images were taken for both FluoTag and DAPI channels, then nuclei were masked with a manually tuned threshold on the DAPI channel and the average value of masked nuclei signal on the FluoTag channel was used to estimate relative abundance of a target across conditions using MATLAB.

#### 4.2 HIST microscopy

HIST imaging was performed using the general design described in Tang et al. ^17^. To accomplish this, we designed a custom laser light path (**Fig S4**). Briefly, lasers were aligned and made colinear using a custom laser launch (described in^91^) before transmitting through an acousto-optic tunable filter (Quanta Tech AOTFnC-400.650-CPCh-TN) to regulate power and laser blanking. After the AOTF, the lasers were fiber coupled into an optical fiber (kineFLEX-HPV) and transmitted into the HIST illumination module. Fiber output was first passed through a Powell lens, before passing through two pairs of beam expanding lenses and focused onto a scanning galvo. This scanning galvo was then relayed onto the back pupil of the objective using relay lenses that are internal to the Nikon microscope. All part numbers, lens distances, and well as a CAD illustration of the mechanical frame are listed in **Fig. S4**. After imaging through the objective, this resulted in an inclined light sheet with a 1.6 µm beam waist (FWHM) at an inclined angle of 39.6 degrees relative to the coverslip plane. Galvo scanning and AOTF timing was synchronized to the lightsheet readout mode of the Orca cameras, with a readout width of 20 pixels, using Labview software similar to that described in^91^. To split the emission light onto two separate cameras and introduce astigmatism for 3D localization microscopy, it was first passed through a 5-band dichroic filter (SPCTRIII 390/475/555/635/747) and tube lens that were internal to the microscope body and then through a 1000 mm focal length cylindrical lens (Thorlabs LJ1516RM-A) that was located approximately 10 mm from the location of the virtual image of the sample plane. This virtual image was then relayed through a pair of achromatic doublet relay lenses (Thorlabs ACT508-300-A). The emission light was split into two separate paths and imaged onto separate cameras using a dichroic filter (Semrock Di03-R561-t3-25x36) that was located halfway between the two relay lenses (**Fig S5**). Each camera also incorporated a spectrally selective emission filter to pass the light from either Cy3B (Semrock FF01-607/70-25) or ATTO655 (Semrock BLP01-635R).

After tuning the objective correction collar for variations in glass thickness, the imaging light sheet angle was fine-tuned prior to imaging to achieve sufficient optical sectioning for differentiation of bound DNA-PAINT fluorophores from unbound imager background and a minimum signal to noise ratio of 2 (200 counts per localization above a dark current of 100). For 3D localization, calibration z-stacks (20 nm spacing, 501 images) were acquired using the adjacent “calibration” well with 100nm TetraSpeck beads and input into SMAP as described below. Chromatic aberrations and slight differences in lateral magnification between cameras were corrected using the calculated positions of TetraSpeck beads in the calibration datasets, which should emit in the same positions across cameras when ideally aligned. Differences in distribution of beads in summed projections were calculated using the createDistanceMatrix function from u-track^92^ and used to fit a second-degree polynomial magnification correction function for correcting lateral positions of localizations.

For each dataset, an 1800 x 512-pixel field of view was selected with a minimum of three in-focus fiducial nanodiamonds and seven nuclei at a focal plane that was roughly 3-4 microns above the coverslip surface. For all DNA-PAINT (antibody) labels, 100,000 images at 50 ms exposures were acquired on both cameras simultaneously using 13.5 mW 560 nm excitation and 43.3 mW 642 nm excitation to excite both the nanodiamond fiducials, Cy3B and ATTO655 probes. For Hoechst 630b labels^93^, a minimum of 600,000 images at 25 ms exposures and 33.4 mW power for 642 nm excitation light were acquired. After imaging each pair of targets, samples were washed at least 6 times for approximately 5 minutes on average in imaging buffer. 2000 images were acquired after the washes to verify that the old imaging strands had been removed. Then, a new pair of imager strands diluted in imaging buffer (for Exchange-PAINT) or, in the final round, the Hoechst 630b blinking dye diluted in PBS to a final concentration of 10 nM (for single molecule localization of DNA^93^) were exchanged with the imaging buffer used for the final wash. 5 mL syringes (Henke 5050-X00V0) and blunt end needles were used to exchange imaging buffer between washes and imaging runs.

To characterize the replicability of Exchange-PAINT imaging over multiple rounds of imaging and labeling (panel F in **Figs. S6-S17**), single Exchange-PAINT targets were imaged in three rounds of 100,000 images at 50 ms exposures (later described as triple repeat labeling experiments), with washes in between as described above. For multiplexed Exchange-PAINT experiments, targets were paired and imaged in seven rounds – the first six being unique pairs of labels and the last a repeat of the lamin A/C and RNA Polymerase II targets – followed by Hoechst 630b labeled imaging of DNA^93^.

#### 4.3 Single molecule localization and registration

For each label, emitters were localized using the MATLAB-based Super-Resolution Microscopy Analysis Platform (SMAP^94^) using a camera conversion factor of 0.24 for Cy3B and 0.23 for ATTO655 fluorophores; a localization ROI size of 7 pixels, a minimum distance of 3 pixels between localizations, and a spline-fit function (the calibrate3DsplinePSF plugin) to the 3D calibration stacks to calculate positions in z. To group Exchange-PAINT binding events across frames and increase localization precision, individual localizations were linked across frames when they were identified within 100 nm radius in sequential frames. To select for stable Exchange-PAINT binding events which typically have a hybridization time of several imaging frames, the detected localizations were filtered to discard any localizations that were >50 nm in lateral precision uncertainty, >200 nm in axial precision uncertainty, <100 emitted photons per localization and to discard any localizations there were not found in at least two sequential frames. The Hoechst-630B datasets utilized similar filters; however, because this probe utilizes a different blinking mechanism than Exchange-PAINT and has a much shorter blinking time, we did not use the final minimum sequential frame filter. Finally, the localization coordinates for each Exchange-PAINT target were all registered in 3D using the common fiducial locations within each dataset.

#### 4.4 SMLM data analysis and rendering

Ripley’s g(r), bi-linear fitting, and fractal dimension calculation was applied to analyze the spatial distribution single molecule localization microscopy datasets as described previously^60^. To ensure that these curves were not impacted by potential repeat localizations from multiple imager strand docking events to the same antibody complex, we only analyzed the G(r) curves from 40 nm up to 5 microns, which is greater than our localization precision and up to roughly half the diameter of a single nucleus. To compare the distributions between different Exchange-PAINT targets, this code was modified to compute the pair cross correlation by using one species as the seed to center each spherical shell and counting the number of molecules for the second species within each shell. Both the Ripley’s G(r) and the PCCF curves were normalized by a simulated random distribution consisting of the same number of localizations within each nucleus. 20 simulated random distributions were created for each comparison to estimate expected error for both the G(r) and PCCF calculations. G(r) and PCCF computations scale as the square of the number of points in each dataset. Therefore, to increase efficiency, these computations were only calculated on a subset of 300,000 localizations from each individual target. To parallelize the computations (13 x 13 targets x 20 simulated random cases x ∼10 cells per condition = 33800 PCCF calls per dataset), these calculations were performed on the computational cluster (UNC Longleaf).

To characterize the imaging repeatability for each target and also to test whether multiplexed labeling with many antibodies in parallel might alter the spatial characteristics of each target (e.g. due to steric hindrance of antibody binding), we conducted triple-repeat labeling experiments wherein samples were labeled for a single target and imaged three sequential times with the same imaging strand (with washes in between each run as described above). From these datasets, we compared the self-association of the label via Ripley’s G(r) within an imaging run (intra-run) with the cross correlation PCCF of the same label computed between different imaging runs (inter-run). Finally, we also plotted the curves as compared to the average self-association g(r) curves computed from the multi-label Exchange-PAINT datasets used in this manuscript. Differences between the intra-run and inter-run curves could arise, for example if there is drift or swelling between imaging runs due to washing steps or if we are incompletely localizing the bound antibody docking strands within the 100,000 imaging frames of a single run. In these cases, we might expect a lower inter-run correlation than intra-run correlation. Likewise, steric hindrance within the multiplex Exchange-PAINT imaging run may lead to lower correlations at short length scales. In general, we found all three curves to be similar for all labels (panel F in **Figs. S6-S17**) with slightly weaker correlations for the intra-run and multiplex-run curves with the most pronounced differences occurring below approximately 100 nm.

To compute the heat maps of the average fold enrichment the PCCF curves, we took the base-10 logarithm of the normalized PCCF curves and computed the average value between the indicated length scales (40 – 200 nm and 200 – 2000 nm) for each target-target comparison in each cell. To compare the difference in enrichment between control and drug treated conditions, we subtracted the average fold enrichment, averaged over all cells, for the drug treated condition from the control condition for each target-target comparison. Due to a few of the comparisons having a much larger change than the majority of the others, we clipped the heatmaps to the specified minimum and maximum values to highlight the patterns of changes and report the significance values on the raw data. We also provide the full (non-clipped) tabulated data for the heatmaps as source data.

Super-resolution images were reconstructed as 3D histograms of molecule counts, then blurred by a Gaussian kernel to match the average localization precision in x, y, and z for each label. Images were rendered with a voxel size appropriate to Nyquist sampling the signal, i.e., near half the average precision, at 6.5 x 6.5 x 27.5 nm.

#### 4.5 Statistical analysis

For all box and whisker plots in this manuscript, the box indicates the inter- quartile range. The horizontal line in the middle indicates median, and the bars indicate 1.5 x the upper and lower limits of inter-quartile range. Dots indicate fitted values from a single cell. To compare changes in the localization densities for each target and to compare the significance of changes to the average fold enrichment between drug-treated vs. control conditions, we utilized a two-sided student’s t-test with each cell considered as an independent sample. All PCCF graphs are presented as mean ± standard deviation. To compare the significance of changes to the hinge points and slopes of each PCCF curve between drug-treated vs. control conditions, we utilized a Wilcoxon rank sum test with each cell considered as an independent sample and report the p-values in **Figs. S24 – S68**. Sample sizes and biological replicates are reported within each figure legend.

## Acknowledgments

We thank Luke Lavis for providing the Hoechst JF630b reagent, Eric Betzig for sharing an early design of the HIST microscopy module, Dan Milkie for assistance with the HIST microscopy software, Michaela Clynes for assistance in antibody concentration optimization, and Yu Shi for his contributions to the single molecule drift correction and linking code. We also thank Regan Moore, Ralf Jungmann, Brian Strahl, Dorothy Erie, Brian Diekman, and Amy Gladfelter for helpful discussions and feedback.

## Author contributions

Conceptualization: WRL, FR Methodology: FR, WRL, VA Investigation: FR, VA, ET, WRL Visualization: FR, CA, WRL Supervision: WRL Writing: FR, WRL with feedback from all authors.

## Data and materials availability

Due to the inordinate size of the raw image data (∼100TB), it is not currently feasible to deposit this into a central repository; however, all datasets underlying the results in this manuscript are available from the corresponding author upon request. To the extent possible, the authors will try to meet all requests for data sharing within 2 weeks from the original request. Source data are provided with this paper. All data needed to evaluate the conclusions in the paper are present in the paper and/or the Supplementary Materials.

