## Supplementary Materials for "Mapping the nuclear landscape with multiplexed super-resolution fluorescence microscopy"

Figure S35. RNAPII repeat vs. all, TSA vs. Control PCCF. 37
Figure S36. RNAPII pS2 vs. all, TSA vs. Control PCCF. 38
Figure S37. RNAPII pS5 vs. all, TSA vs. Control PCCF. 39
Figure S38. SC-35 vs. all, TSA vs. Control PCCF. 40
Figure S39. CDK9 vs. all, A-485 vs. Control PCCF. 41
Figure S40. H3K4me1 vs. all, A-485 vs. Control PCCF. 42
Figure S41. H3K9me3 vs. all, A-485 vs. Control PCCF. 43
Figure S42. H3K27ac vs. all, A-485 vs. Control PCCF. 44
Figure S43. H3K27me3 vs. all, A-485 vs. Control PCCF. 45
Figure S44. Hoechst vs. all, A-485 vs. Control PCCF. 46
Figure S45. HP1α vs. all, A-485 vs. Control PCCF. 47
Figure S46. Lamin A/C vs. all, A-485 vs. Control PCCF. 48
Figure S47. Lamin A/C repeat vs. all, A-485 vs. Control PCCF. 49
Figure S48. p300 vs. all, A-485 vs. Control PCCF. 50
Figure S49. RNAPII vs. all, A-485 vs. Control PCCF. 51
Figure S50. RNAPII repeat vs. all, A-485 vs. Control PCCF. 52
Figure S51. RNAPII pS2 vs. all, A-485 vs. Control PCCF. 53
Figure S52. RNAPII pS5 vs. all, A-485 vs. Control PCCF. 54
Figure S53. SC-35 vs. all, A-485 vs. Control PCCF. 55
Figure S54. CDK9 vs. all, αAM vs. Control PCCF. 56
Figure S55. H3K4me1 vs. all, αAM vs. Control PCCF. 57
Figure S56. H3K9me3 vs. all, αAM vs. Control PCCF. 58
Figure S57. H3K27ac vs. all, αAM vs. Control PCCF. 59
Figure S58. H3K27me3 vs. all, αAM vs. Control PCCF. 60
Figure S59. Hoechst vs. all, αAM vs. Control PCCF. 61
Figure S60. HP1α vs. all, αAM vs. Control PCCF. 62
Figure S61. Lamin A/C vs. all, αAM vs. Control PCCF. 63
Figure S62. Lamin A/C repeat vs. all, αAM vs. Control PCCF. 64
Figure S63. p300 vs. all, αAM vs. Control PCCF. 65
Figure S64. RNAPII vs. all, αAM vs. Control PCCF. 66
Figure S65. RNAPII repeat vs. all, αAM vs. Control PCCF. 67
Figure S66. RNAPII pS2 vs. all, αAM vs. Control PCCF. 68
Figure S67. RNAPII pS5 vs. all, αAM vs. Control PCCF. 69
Figure S68. SC-35 vs. all, αAM vs. Control PCCF. 70
Figure S69. Widefield imaging and quantification of perturbation effects 71
Table S1. Primary antibody and DNA-PAINT probe characteristics 72
Table S2. Hinge points of the PCCF curves for control cells 73
Table S3. Short length-scale slopes of the PCCF curves for control cells 74
Table S4. Long length-scale slopes of the PCCF curves for control cells 75
Table S5. Primary antibody screening results 76
Movie S1. Representative multiplex rendering of a single HIST field of view 78
Supplementary Note 1 79

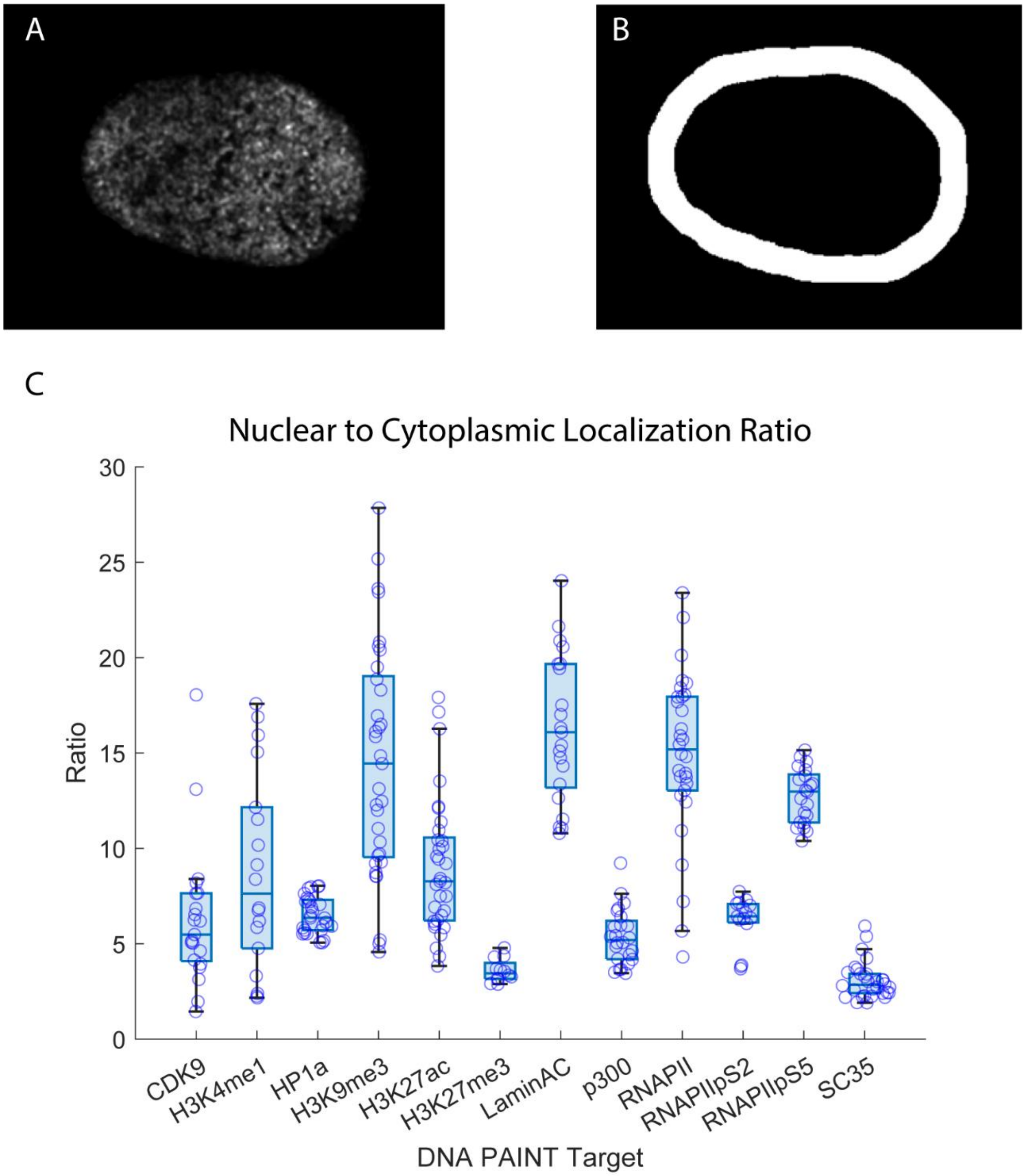

**Figure S1. Nuclear specificity estimates for DNA-PAINT targets.**

(A) Representative nuclear-masked DNA-PAINT reconstruction (RNAPII S2p). (B) Representative dilated and subtracted mask used to estimate cytoplasmic localization density. (C) Ratio of nuclear localization density to cytoplasmic localization density (estimated immediately outside nuclear mask edge). Points represent individual masks across 3 imaging runs with a minimum of n=4 cells (mean 8 cells).

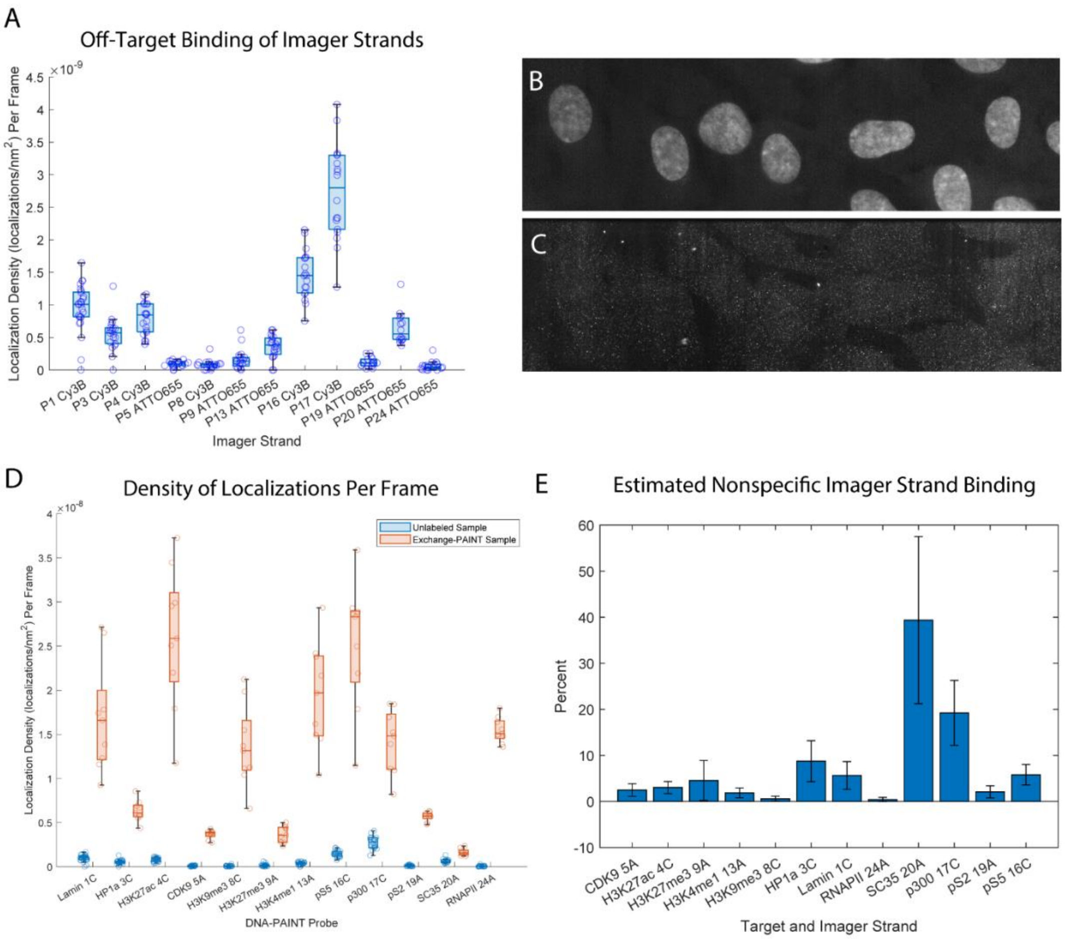

**Figure S2. Non-specific binding estimates for DNA-PAINT imager strands.**

(A) Nonspecific sample binding of imager strands (localization density per frame) within nuclear regions on a sample labeled with DAPI and with imager strands at concentrations matched to control Exchange-PAINT datasets – 0.25 nM for P8 Cy3B (8C); 1 nM for P9 ATTO655 (9A), P13 ATTO655 (13A), P19 ATTO655 (19A), and P24 ATTO655 (24A); 2.5 nM for P1 Cy3B (1C), P3 Cy3B (3C), P4 Cy3B (4C), P16 Cy3B (16C), P17 Cy3B (17C), and P20 ATTO655 (20A); and 5 nM for P5 ATTO655 (5A). Nuclear regions were masked semi-automatically using DAPI staining and manually edited to exclude fiducial markers or extranuclear signal as needed. Points represent a minimum of n=15 individual nuclear masks across two fields of view per probe (with a mean of 20 masks per probe). (B) Representative maximum intensity projection (MIP) of DNA stained with DAPI on a selected field of view. Nuclear masks were generated with manual thresholding of DAPI stain. (C) Representative MIP of P1 Cy3B binding on the same field of view as in (B) over 2000 frames, 50 ms exposure. (D) Localization density per frame of nonspecifically bound imager strands (blue) vs. localization density per frame of DNA-PAINT targets in the first control Exchange-PAINT dataset within nuclear regions (n=9, orange). (E) Percent of binding events estimated as nonspecific based on the fraction of off-target localization density per frame over total off-target and on-target localization density per frame. Error bars represent standard deviation after multiplicative error propagation.

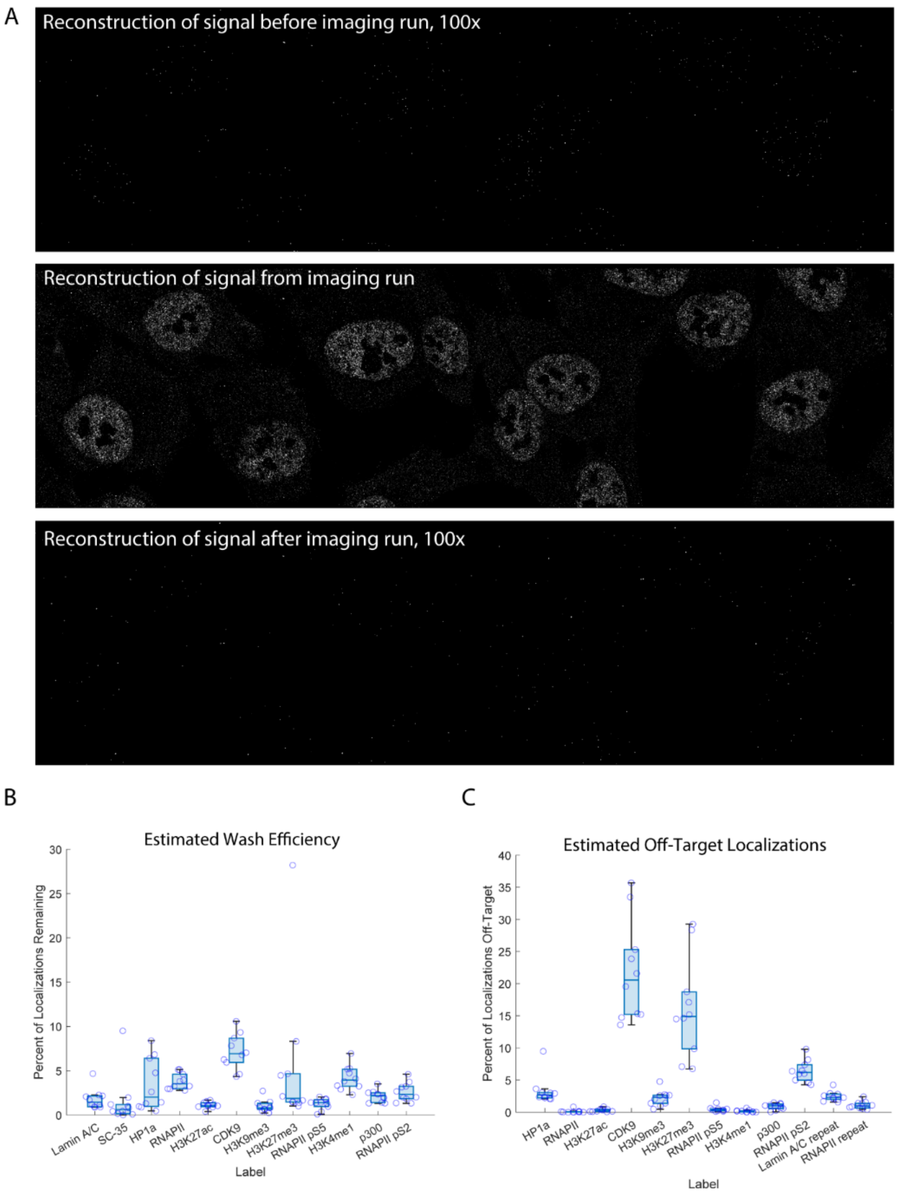

**Figure S3. Wash efficiency and off-target localization estimates for DNA-PAINT imager strands.**

(A) Top: representative reconstruction from 2000 frames of imaging after washing and prior to imaging RNAPII pS2 on corresponding camera, 100x intensity-scaled relative to middle panel. Middle: reconstruction from 2000 frames of imaging RNAPII pS2. Bottom: reconstruction of 2000 frames of imaging after washing, following RNAPII pS2 imaging run on corresponding camera, 100x intensity-scaled relative to middle panel. (B) Wash efficiency percentage calculated using localization density from last 2000 frames for each DNA-PAINT probe within masks vs. localization density on corresponding camera after washing. Points represent n=10 cells. (C) Estimated off-target localization percentage due to prior imager strand crosstalk calculated using localization density after washing vs. localization density on corresponding camera from first 2000 frames within masks for each probe. Points represent n=10 cells.

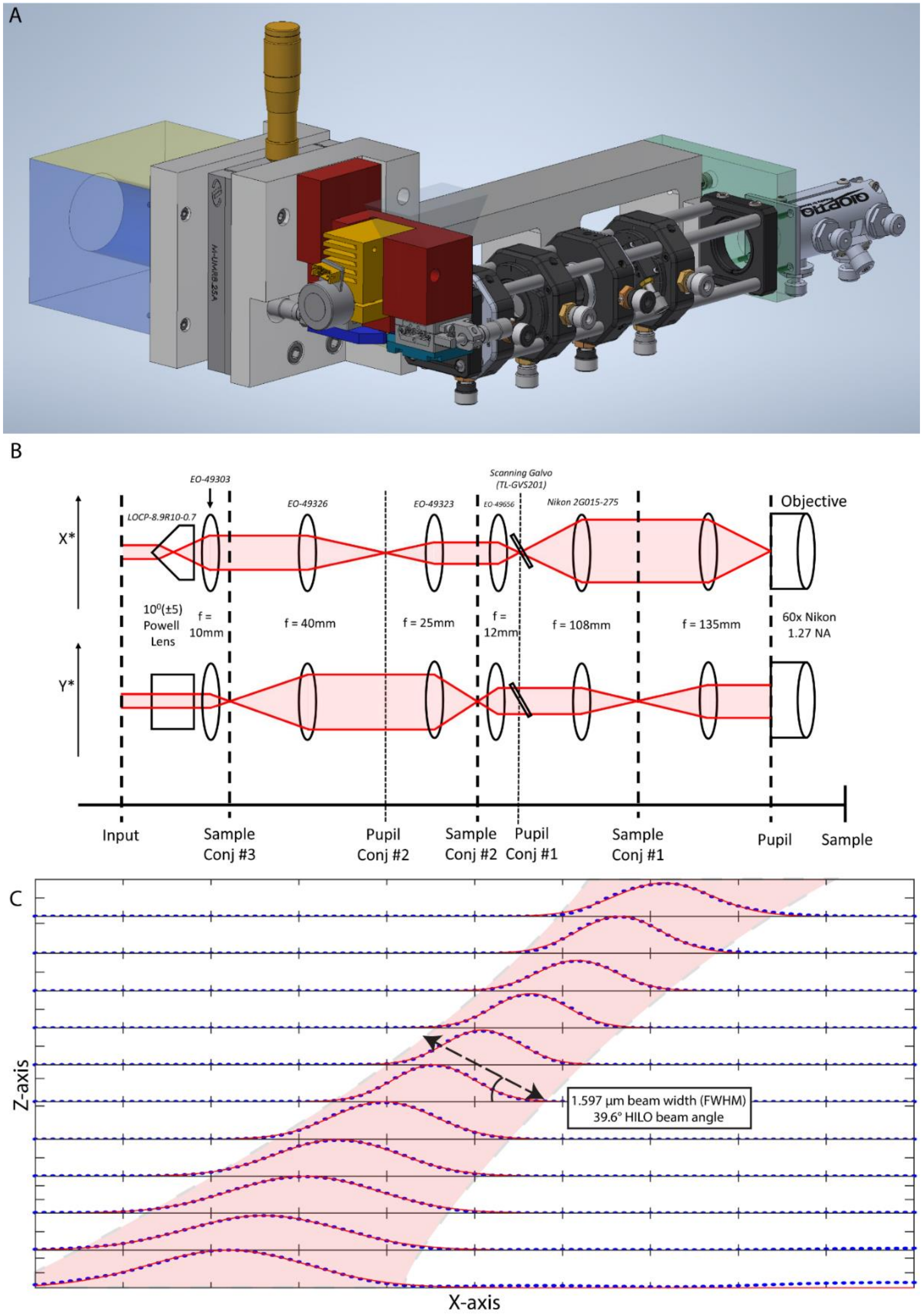

**Figure S4. Custom HIST beam shaping attachment.**

(A) CAD Model of the HIST beam shaping attachment. (B) Simplified optical path for the HIST beam shaping attachment. (C) Characterization of the produced HIST beam for a typical imaging session.

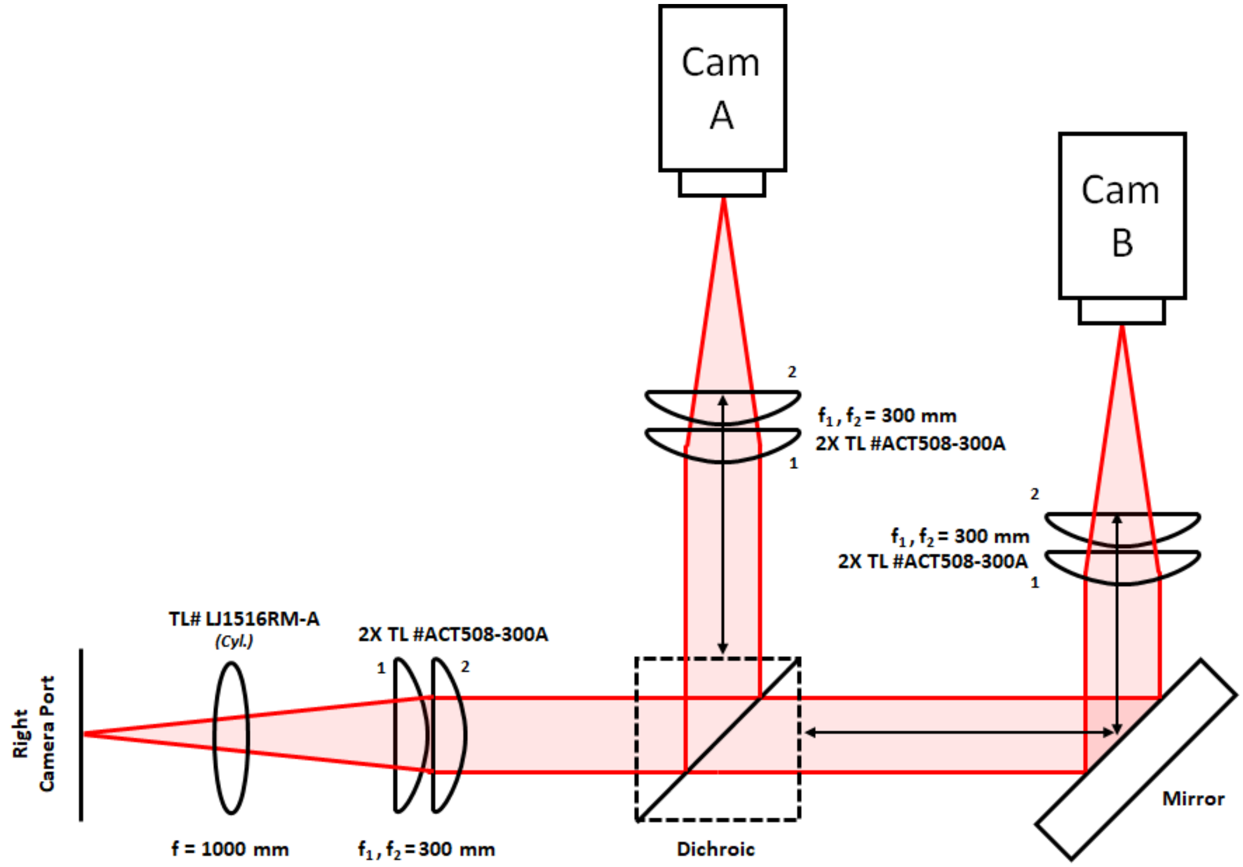

**Figure S5. HIST emission path.**

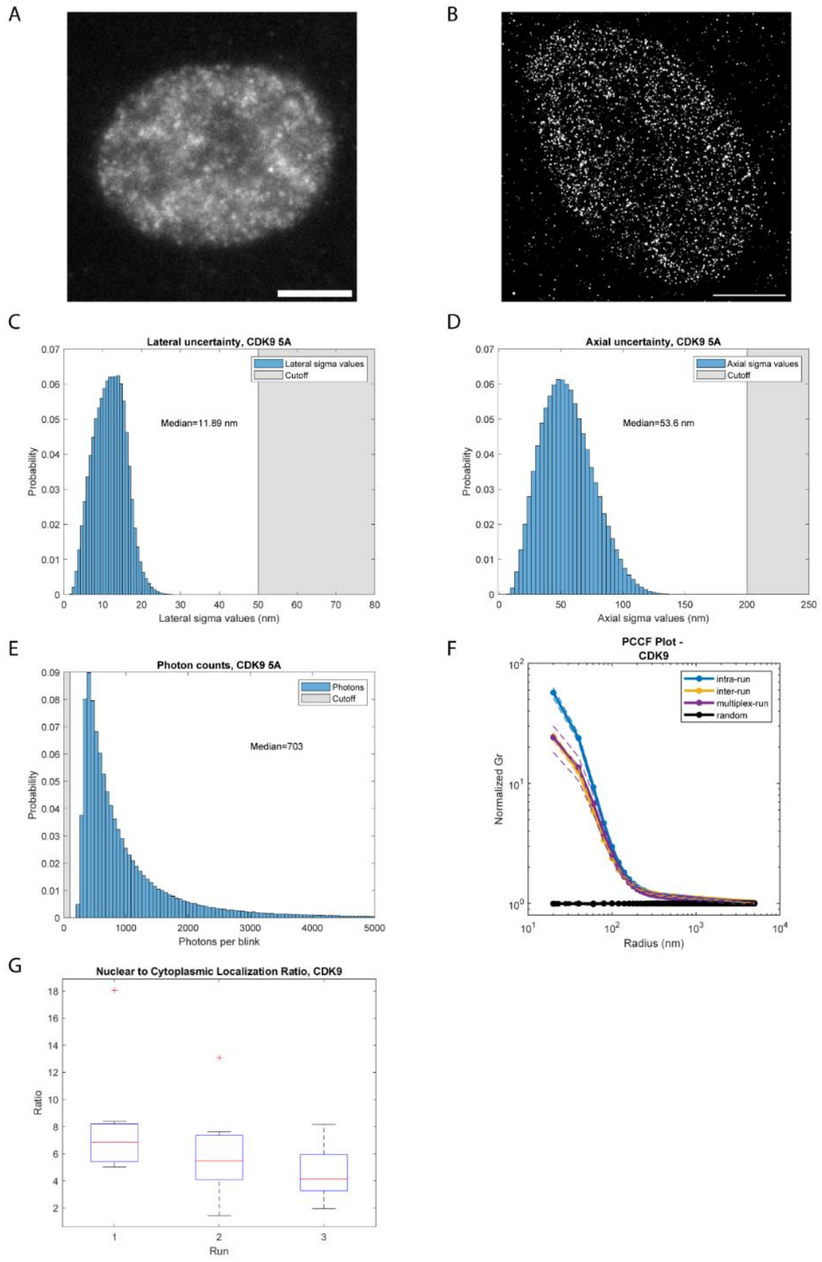

**Figure S6. Characterization of single molecule localization imaging of the target CDK9.**

(A) Widefield (60x) imaging of CDK9 within a single nucleus, labeled with anti-rabbit FluoTag 488 (FT488). Scale bar represents 5 µm. (B) DNA-PAINT single molecule localization (SML) reconstruction of CDK9, labeled with Probe 5 (P5) ATTO655. Scale bar represents 10 µm. (C-E) Post-filtering single molecule localization histograms from representative control Exchange-PAINT dataset. (C) Photon counts per localization. (D) Lateral uncertainty per localization. (E) Axial uncertainty per localization. (F) Characterization of replicable DNA-PAINT imaging with the pair cross-correlation function (PCCF). Inter- and intra-run error represents standard deviation of n=6 cells across three imaging runs. Multiplex-run error represents standard deviation of n=31 cells across three biological replicates. (G) Box plot of nuclear to cytoplasmic localization ratio for three imaging runs across n=6 cells.

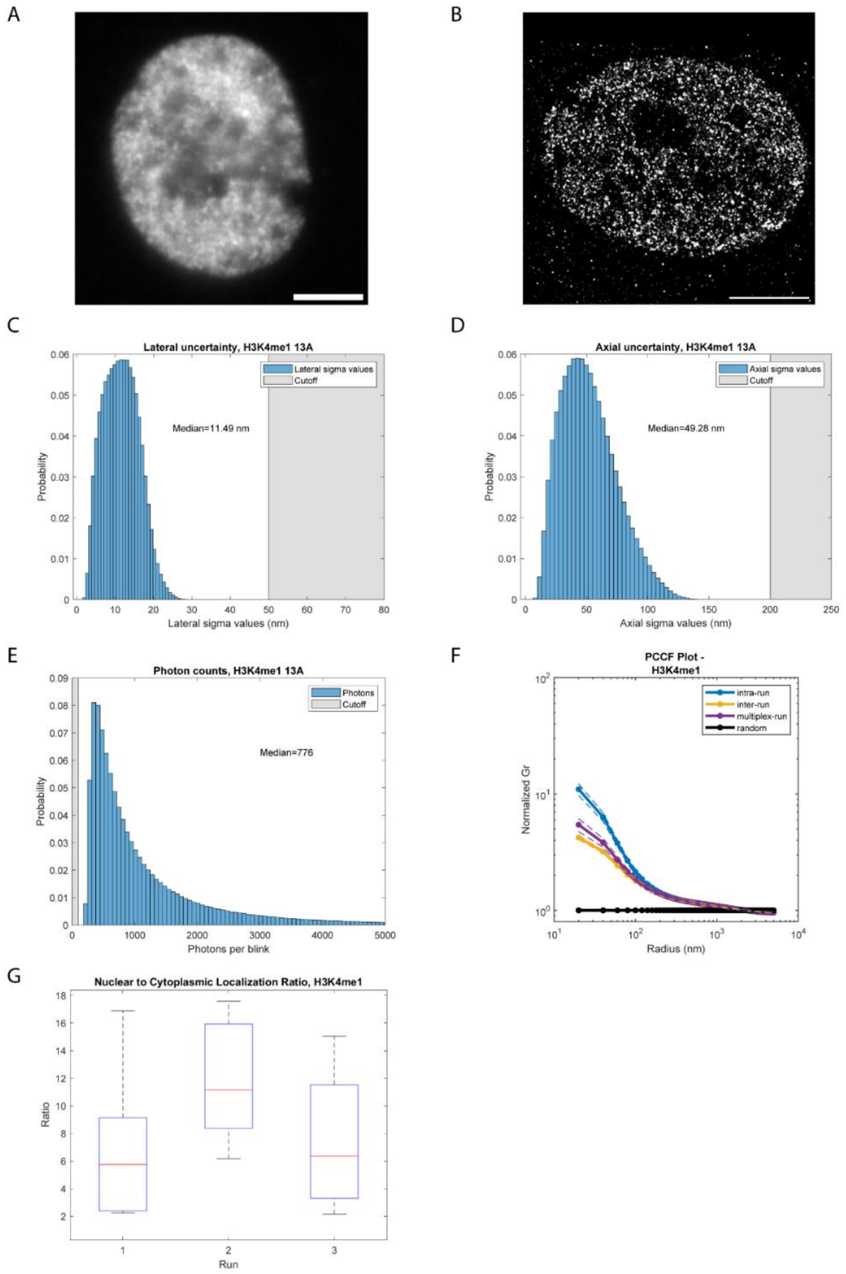

**Figure S7. Characterization of single molecule localization imaging of the target H3K4me1.**

(A) Widefield (60x) imaging of H3K4me1 within a single nucleus, labeled with anti-rabbit FluoTag 488 (FT488). (B) DNA-PAINT single molecule localization (SML) reconstruction of H3K4me1, labeled with P13 ATTO655. (C-E) Post-filtering single molecule localization histograms from representative control Exchange-PAINT dataset. (C) Photon counts per localization. (D) Lateral uncertainty per localization. (E) Axial uncertainty per localization. (F) Characterization of replicable DNA-PAINT imaging with the pair cross-correlation function (PCCF). Inter- and intra-run error represents standard deviation of n=6 cells across three imaging runs. Multiplex-run error represents standard deviation of n=31 cells across three biological replicates. (G) Box plot of nuclear to cytoplasmic localization ratio for three imaging runs across n=6 cells.

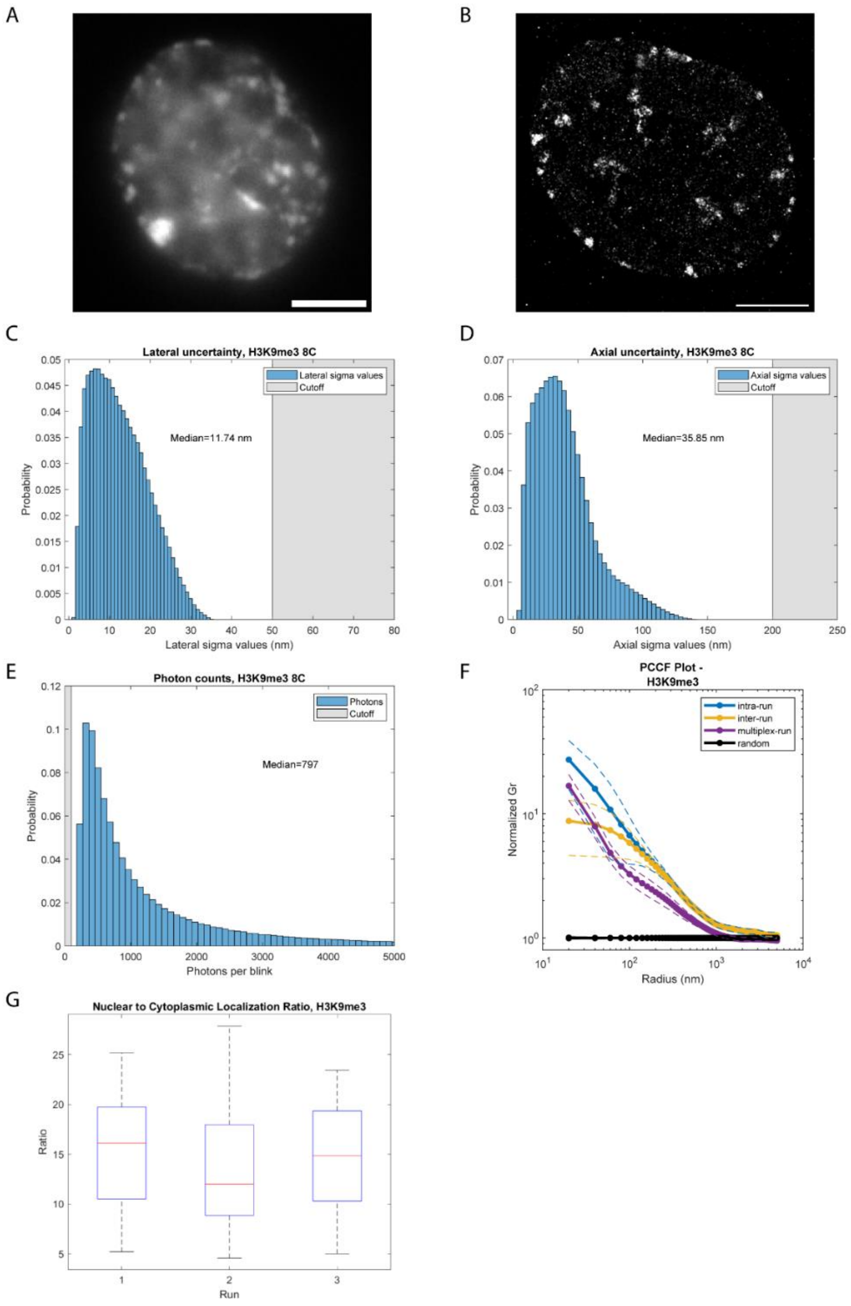

**Figure S8. Characterization of single molecule localization imaging of the target H3K9me3**.

(A) Widefield (60x) imaging of H3K9me3 within a single nucleus, labeled with anti-rabbit FluoTag 488 (FT488). (B) DNA-PAINT single molecule localization (SML) reconstruction of H3K4me1, labeled with P8 Cy3B. (C-E) Post-filtering single molecule localization histograms from representative control Exchange-PAINT dataset. (C) Photon counts per localization. (D) Lateral uncertainty per localization. (E) Axial uncertainty per localization. (F) Characterization of replicable DNA-PAINT imaging with the pair cross-correlation function (PCCF). Inter- and intra-run error represents standard deviation of n=11 cells across three imaging runs. Multiplex-run error represents standard deviation of n=31 cells across three biological replicates. (G) Box plot of nuclear to cytoplasmic localization ratio for three imaging runs across n=11 cells.

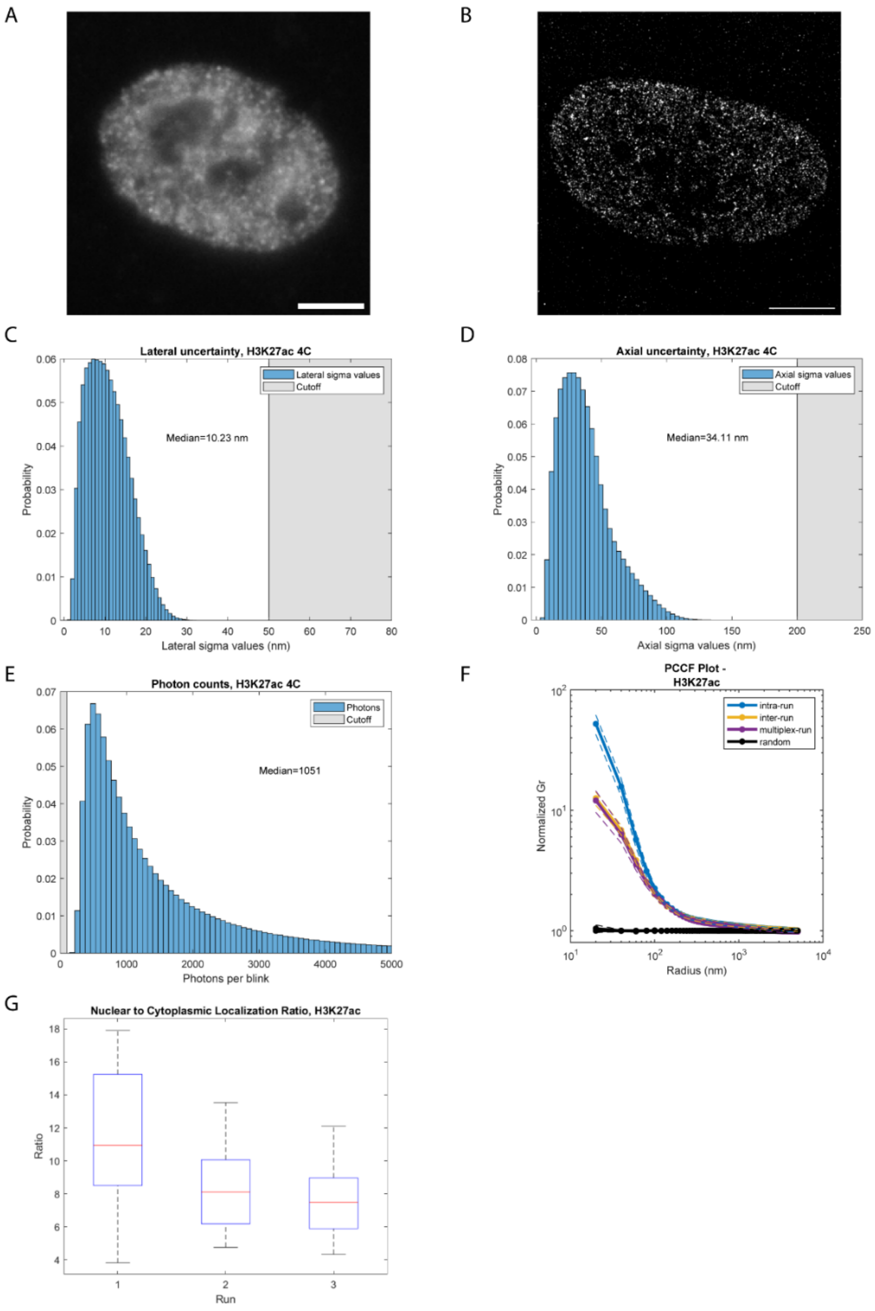

**Figure S9. Characterization of single molecule localization imaging of the target H3K27ac.**

(A) Widefield (60x) imaging of H3K27ac within a single nucleus, labeled with anti-rabbit FluoTag 488 (FT488). (B) DNA-PAINT single molecule localization (SML) reconstruction of H3K27ac, labeled with P4 Cy3B. (C-E) Post-filtering single molecule localization histograms from representative control Exchange-PAINT dataset. (C) Photon counts per localization. (D) Lateral uncertainty per localization. (E) Axial uncertainty per localization. (F) Characterization of replicable DNA-PAINT imaging with the pair cross-correlation function (PCCF). Inter- and intra-run error represents standard deviation of n=10 cells across three imaging runs. Multiplex-run error represents standard deviation of n=31 cells across three biological replicates. (G) Box plot of nuclear to cytoplasmic localization ratio for three imaging runs across n=10 cells.

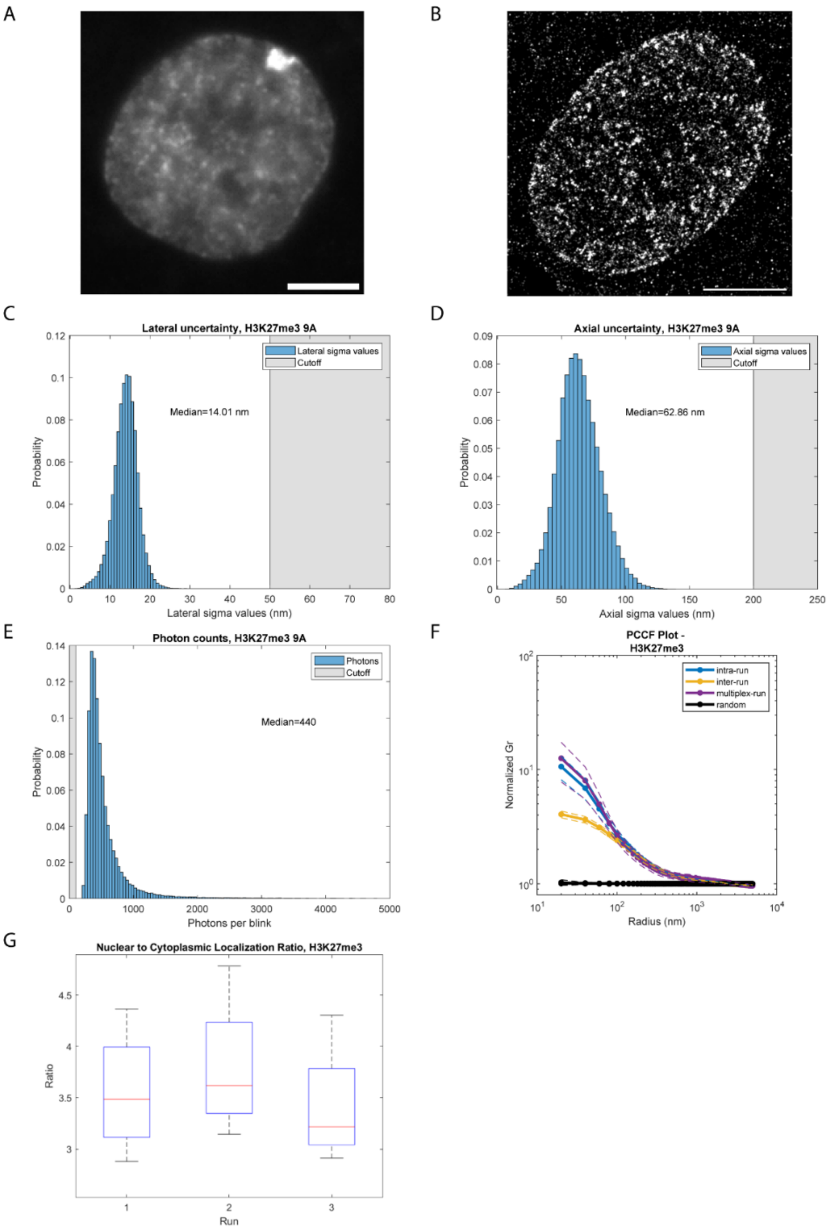

**Figure S10. Characterization of single molecule localization imaging of the target H3K27me3.**

(A) Widefield (60x) imaging of H3K27me3 within a single nucleus, labeled with anti-rabbit FluoTag 488 (FT488). (B) DNA-PAINT single molecule localization (SML) reconstruction of H3K27me3, labeled with P9 ATTO655. (C-E) Post-filtering single molecule localization histograms from representative control Exchange-PAINT dataset. (C) Photon counts per localization. (D) Lateral uncertainty per localization. (E) Axial uncertainty per localization. (F) Characterization of replicable DNA-PAINT imaging with the pair cross-correlation function (PCCF). Inter- and intra-run error represents standard deviation of n=6 cells across three imaging runs. Multiplex-run error represents standard deviation of n=31 cells across three biological replicates. (G) Box plot of nuclear to cytoplasmic localization ratio for three imaging runs across n=6 cells.

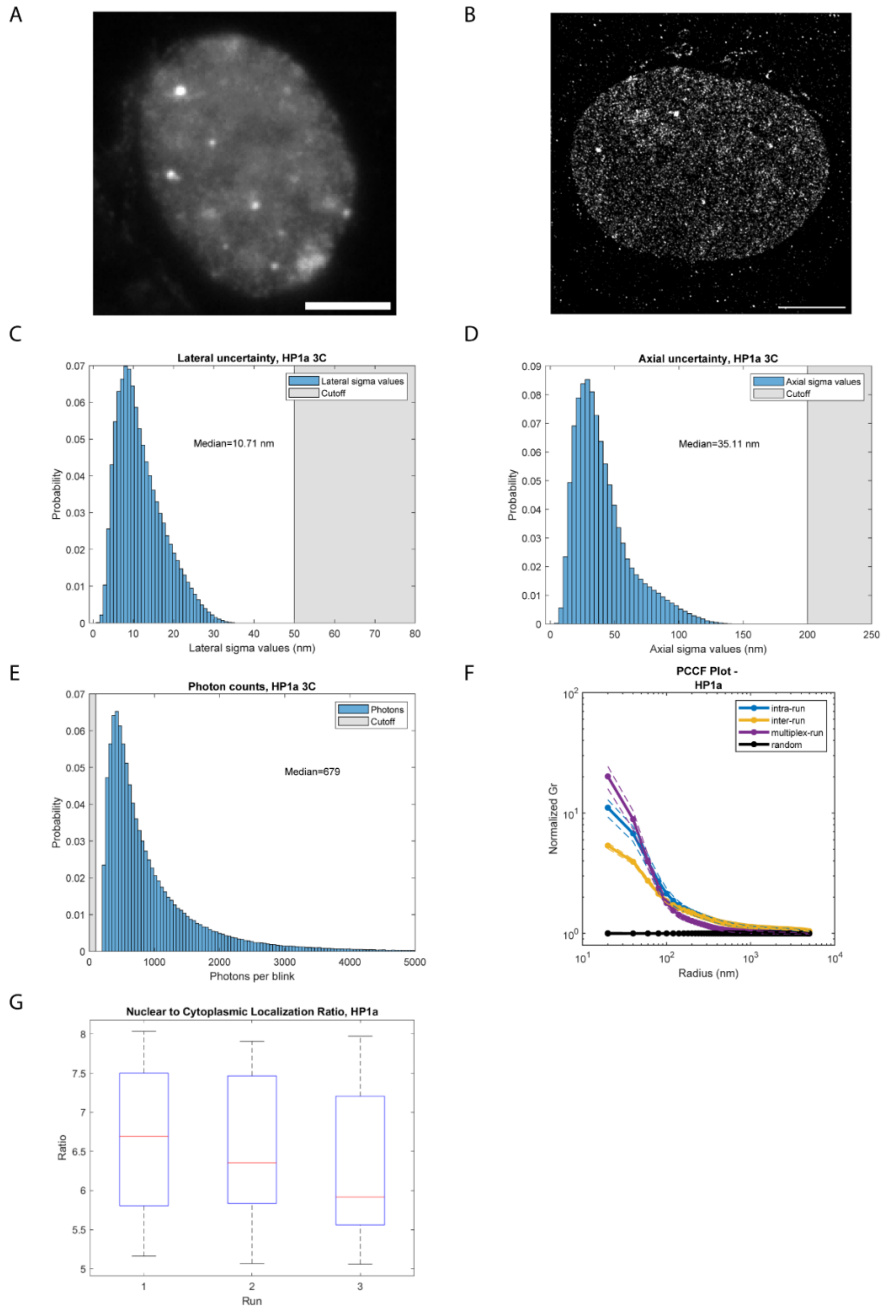

**Figure S11. Characterization of single molecule localization imaging of the target HP1α.**

(A) Widefield (60x) imaging of HP1α within a single nucleus, labeled with anti-rabbit FluoTag 488 (FT488). (B) DNA-PAINT single molecule localization (SML) reconstruction of HP1α, labeled with P3 Cy3B. (C-E) Post-filtering single molecule localization histograms from representative control Exchange-PAINT dataset. (C) Photon counts per localization. (D) Lateral uncertainty per localization. (E) Axial uncertainty per localization. (F) Characterization of replicable DNA-PAINT imaging with the pair cross-correlation function (PCCF). Inter- and intra-run error represents standard deviation of n=11 cells across three imaging runs. Multiplex-run error represents standard deviation of n=31 cells across three biological replicates. (G) Box plot of nuclear to cytoplasmic localization ratio for three imaging runs across n=11 cells.

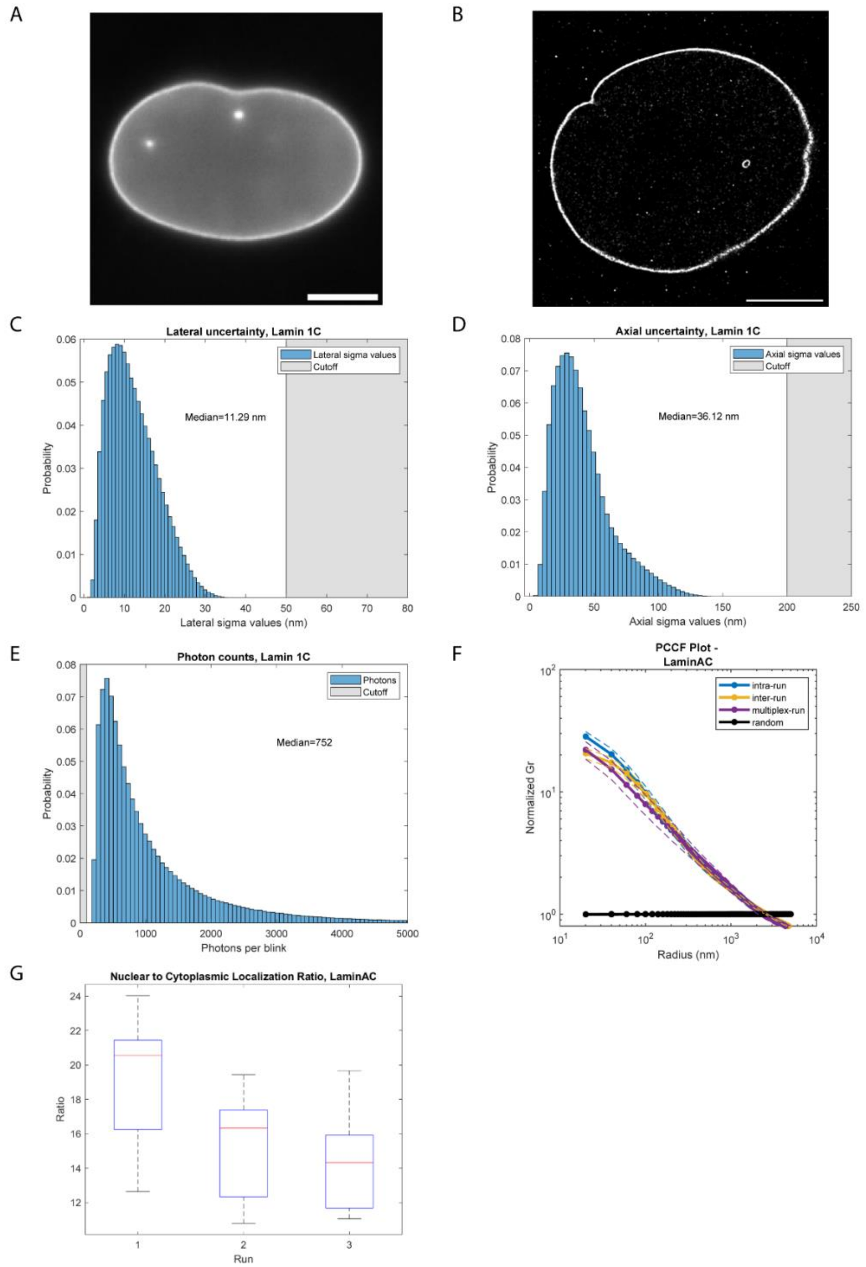

**Figure S12. Characterization of single molecule localization imaging of the target Lamin A/C.**

(A) Widefield (60x) imaging of Lamin A/C within a single nucleus, labeled with anti-mouse FluoTag 488 (FT488). (B) DNA-PAINT single molecule localization (SML) reconstruction of H3K27ac, labeled with P1 Cy3B. (C-E) Post-filtering single molecule localization histograms from representative control Exchange-PAINT dataset. (C) Photon counts per localization. (D) Lateral uncertainty per localization. (E) Axial uncertainty per localization. (F) Characterization of replicable DNA-PAINT imaging with the pair cross-correlation function (PCCF). Inter- and intra-run error represents standard deviation of n=7 cells across three imaging runs. Multiplex-run error represents standard deviation of n=31 cells across three biological replicates. (G) Box plot of nuclear to cytoplasmic localization ratio for three imaging runs across n=7 cells.

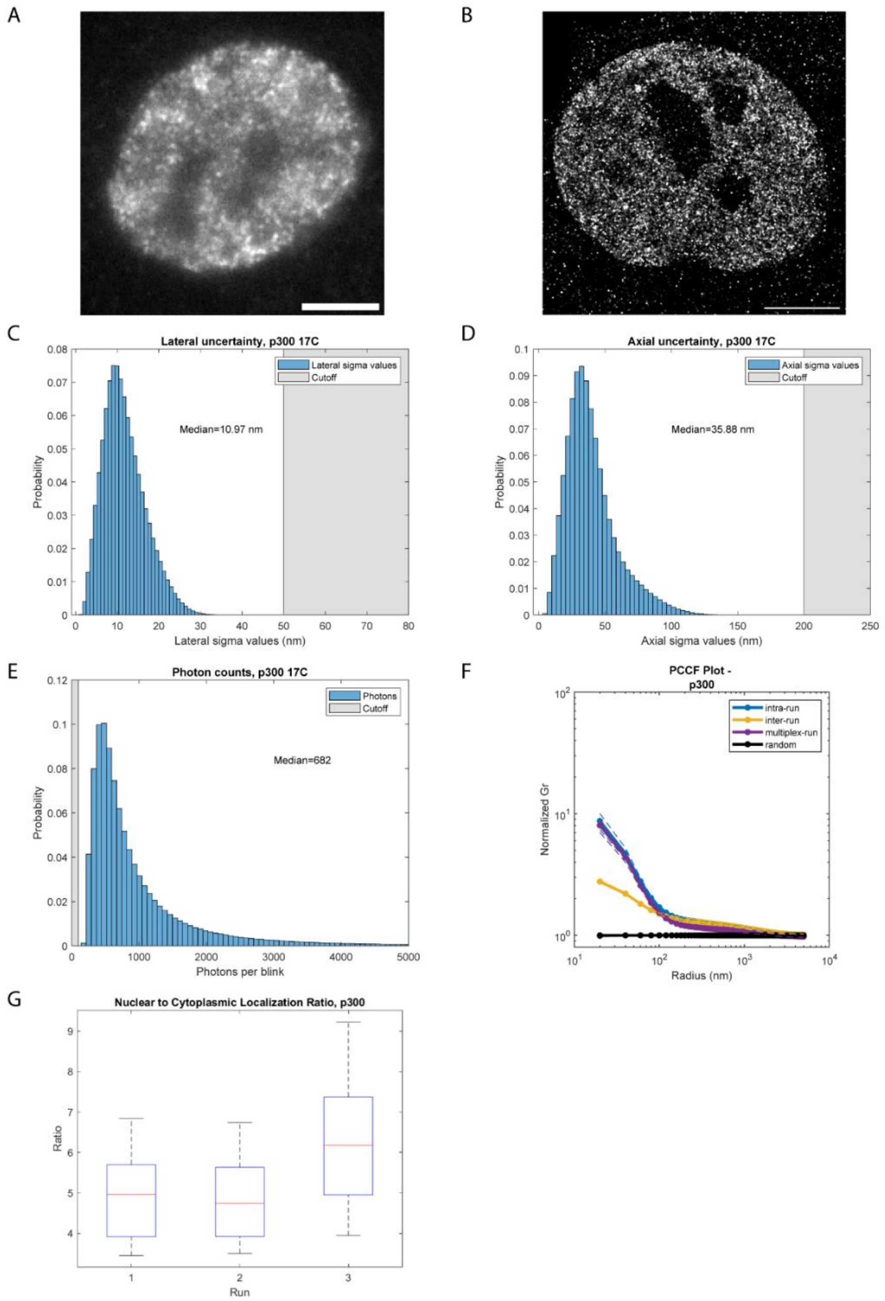

**Figure S13. Characterization of single molecule localization imaging of the target p300.**

(A) Widefield (60x) imaging of p300 within a single nucleus, labeled with anti-rabbit FluoTag 488 (FT488). (B) DNA-PAINT single molecule localization (SML) reconstruction of p300, labeled with P17 Cy3B. (C-E) Post-filtering single molecule localization histograms from representative control Exchange-PAINT dataset. (C) Photon counts per localization. (D) Lateral uncertainty per localization. (E) Axial uncertainty per localization. (F) Characterization of replicable DNA-PAINT imaging with the pair cross-correlation function (PCCF). Inter- and intra-run error represents standard deviation of n=8 cells across three imaging runs. Multiplex-run error represents standard deviation of n=31 cells across three biological replicates. (G) Box plot of nuclear to cytoplasmic localization ratio for three imaging runs across n=8 cells.

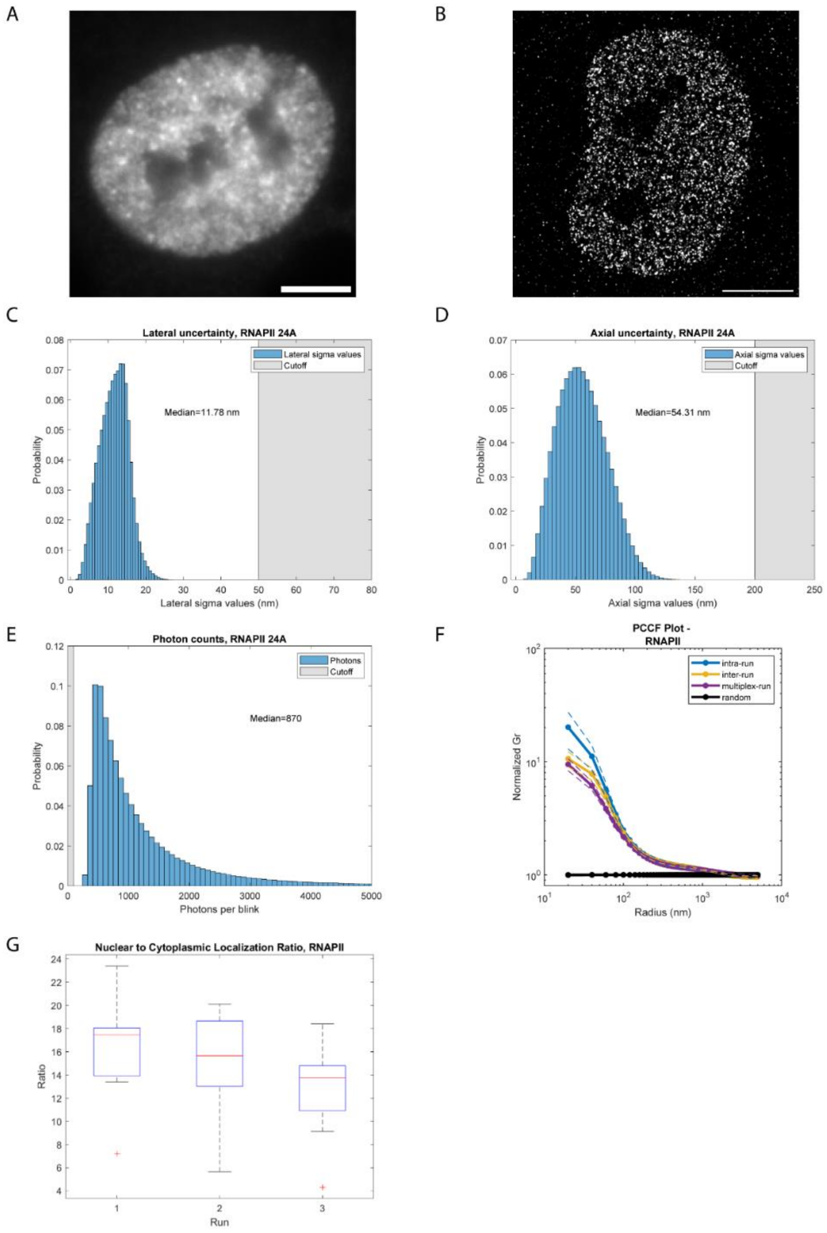

**Figure S14. Characterization of single molecule localization imaging of the target RNA Polymerase II.**

(A) Widefield (60x) imaging of RNA Polymerase II (RNAPII) within a single nucleus, labeled with anti-mouse FluoTag 488 (FT488). (B) DNA-PAINT single molecule localization (SML) reconstruction of RNAPII, labeled with P24 ATTO655. (C-E) Post-filtering single molecule localization histograms from representative control Exchange-PAINT dataset. (C) Photon counts per localization. (D) Lateral uncertainty per localization. (E) Axial uncertainty per localization. (F) Characterization of replicable DNA-PAINT imaging with the pair cross-correlation function (PCCF). Inter- and intra-run error represents standard deviation of n=9 cells across three imaging runs. Multiplex-run error represents standard deviation of n=31 cells across three biological replicates. (G) Box plot of nuclear to cytoplasmic localization ratio for three imaging runs across n=9 cells.

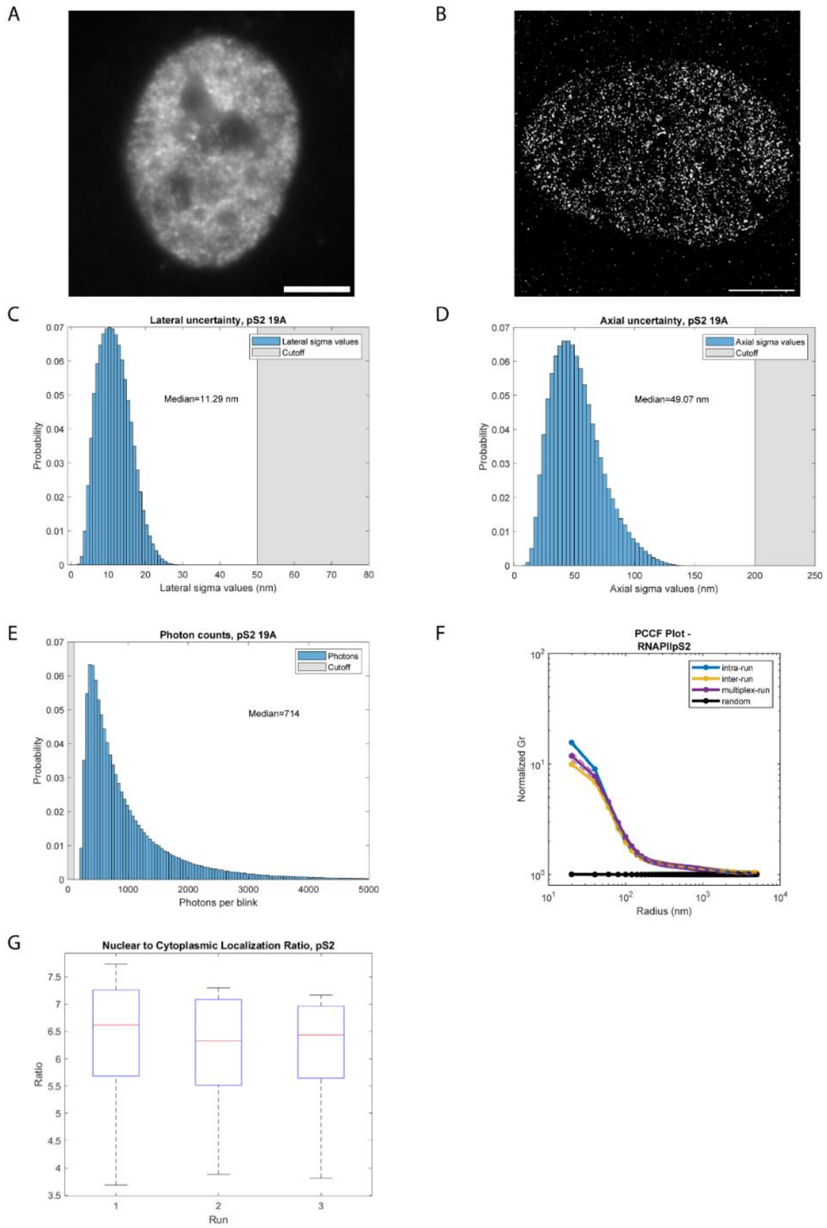

**Figure S15. Characterization of single molecule localization imaging of the target RNA Polymerase Ser2p.**

(A) Widefield (60x) imaging of RNA Polymerase Ser2p (RNAPII pS2) within a single nucleus, labeled with anti-rabbit FluoTag 488 (FT488). (B) DNA-PAINT single molecule localization (SML) reconstruction of RNAPII pS2, labeled with P19 ATTO655. (C-E) Post-filtering single molecule localization histograms from representative control Exchange-PAINT dataset. (C) Photon counts per localization. (D) Lateral uncertainty per localization. (E) Axial uncertainty per localization. (F) Characterization of replicable DNA-PAINT imaging with the pair cross-correlation function (PCCF). Inter- and intra-run error represents standard deviation of n=4 cells across three imaging runs. Multiplex-run error represents standard deviation of n=31 cells across three biological replicates. (G) Box plot of nuclear to cytoplasmic localization ratio for three imaging runs across n=4 cells.

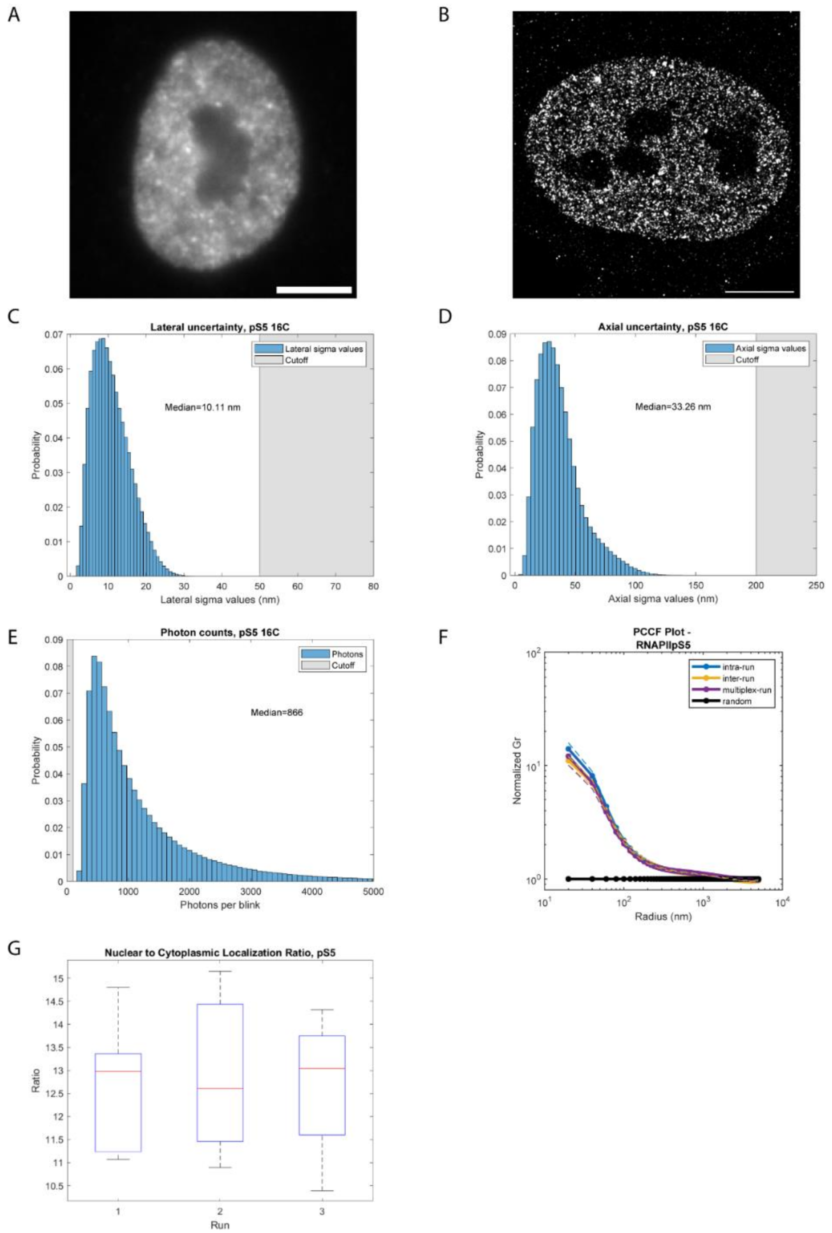

**Figure S16. Characterization of single molecule localization imaging of the target RNA Polymerase Ser5p.**

(A) Widefield (60x) imaging of RNA Polymerase Ser5p (RNAPII pS5) within a single nucleus, labeled with anti-rabbit FluoTag 488 (FT488). (B) DNA-PAINT single molecule localization (SML) reconstruction of RNAPII pS5, labeled with P16 Cy3B. (C-E) Post-filtering single molecule localization histograms from representative control Exchange-PAINT dataset. (C) Photon counts per localization. (D) Lateral uncertainty per localization. (E) Axial uncertainty per localization. (F) Characterization of replicable DNA-PAINT imaging with the pair cross-correlation function (PCCF). Inter- and intra-run error represents standard deviation of n=7 cells across three imaging runs. Multiplex-run error represents standard deviation of n=31 cells across three biological replicates. (G) Box plot of nuclear to cytoplasmic localization ratio for three imaging runs across n=7 cells.

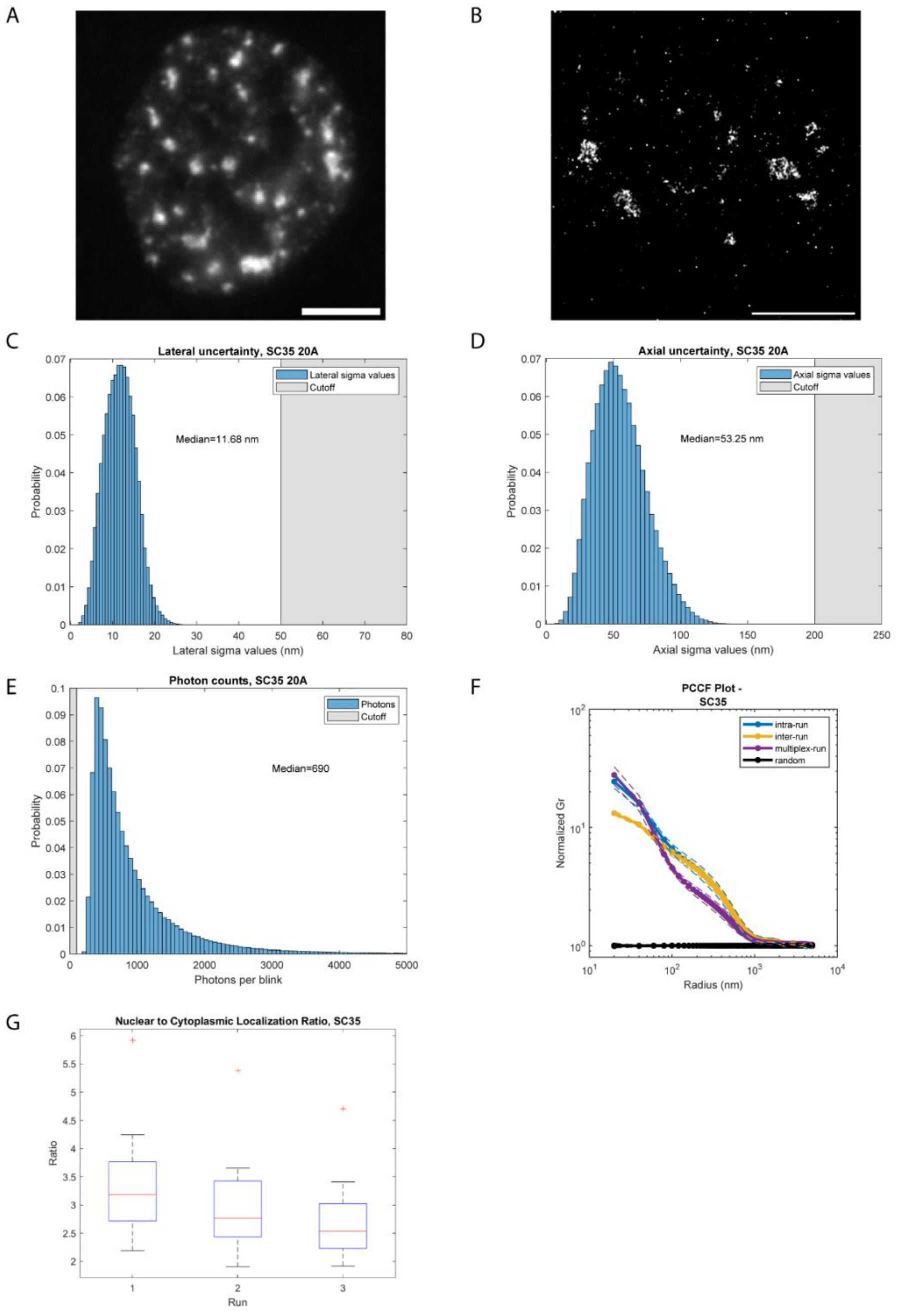

**Figure S17. Characterization of single molecule localization imaging of the target SC-35.**

(A) Widefield (60x) imaging of SC-35 within a single nucleus, labeled with anti-mouse FluoTag 488 (FT488). (B) DNA-PAINT single molecule localization (SML) reconstruction of SC-35, labeled with P20 ATTO655. (C-E) Post-filtering single molecule localization histograms from representative control Exchange-PAINT dataset. (C) Photon counts per localization. (D) Lateral uncertainty per localization. (E) Axial uncertainty per localization. (F) Characterization of replicable DNA-PAINT imaging with the pair cross-correlation function (PCCF). Inter- and intra-run error represents standard deviation of n=9 cells across three imaging runs. Multiplex-run error represents standard deviation of n=31 cells across three biological replicates. (G) Box plot of nuclear to cytoplasmic localization ratio for three imaging runs across n=9 cells.

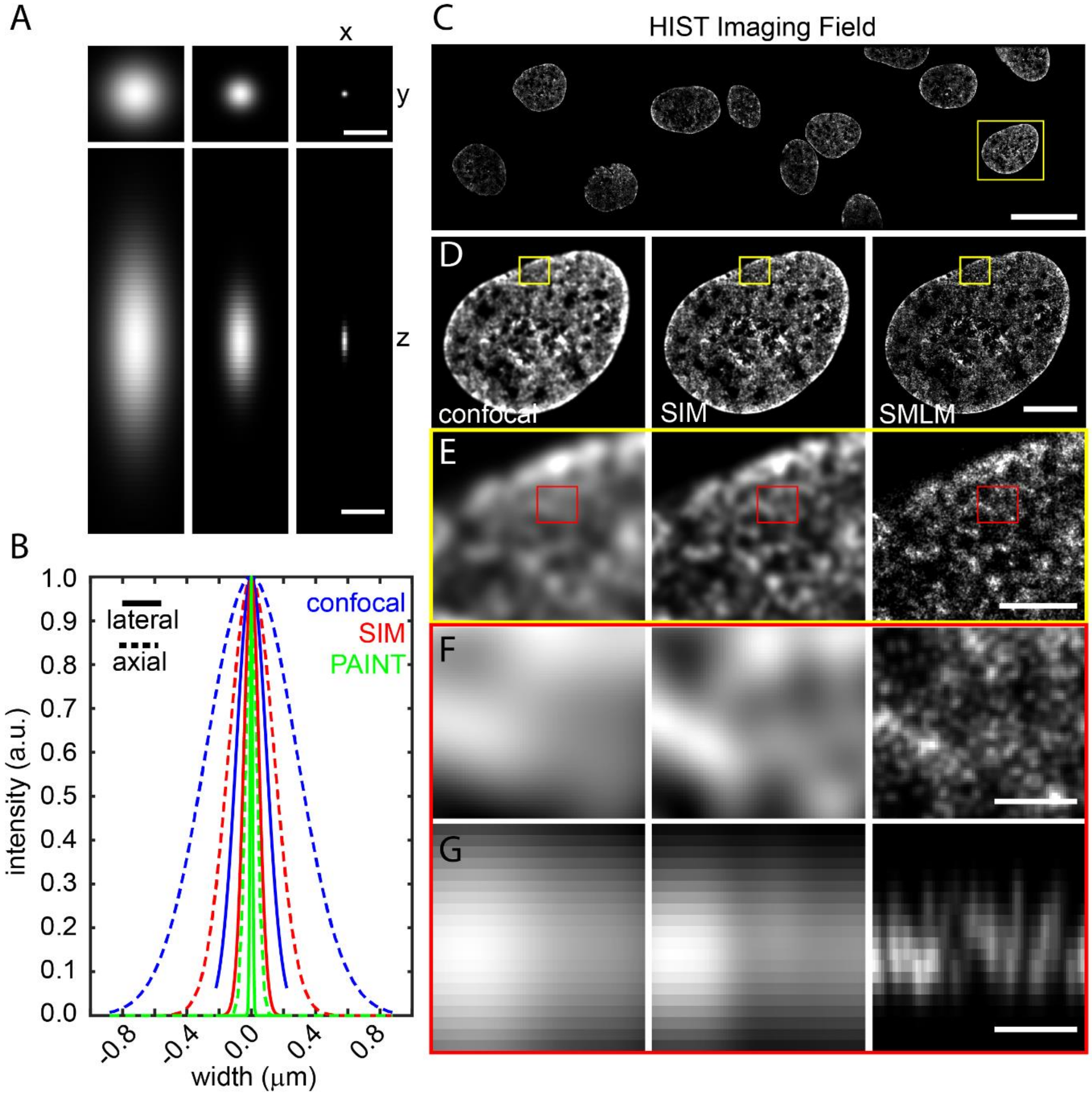

**Figure S18. Resolution comparison of nuclear Exchange-PAINT with simulated confocal and structured illumination microscopy (SIM) datasets.**

(A, B) Intensity plots of simulated point spread functions for confocal (blue), SIM (red), and SMLM (green) imaging in xy (lateral, solid line) and z (axial, dotted). (C) Representative HIST field of view with SMLM rendering of DNA label (Hoechst). (D) Computationally simulated confocal, structured illumination microscopy, and Exchange-PAINT renderings of inset in (C). (E-F) Zoomed-in views of the regions indicated in (D). (G) Axial XZ projection through a single slice midway through (F). Scale bars = 200 nm – A, 20 µm – C, 5 µm – D, 1 µm – E, 200 nm – F, G.

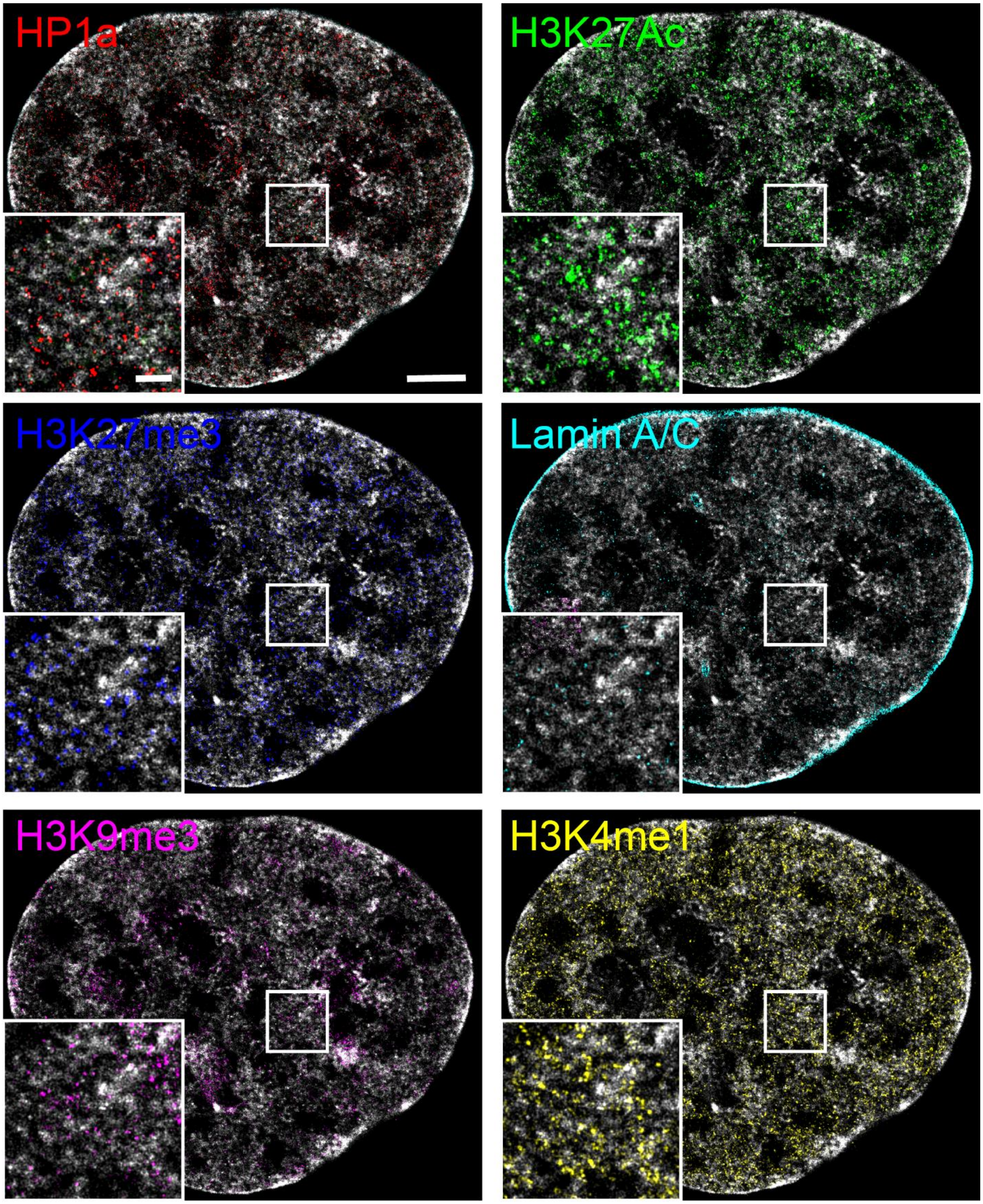

**Figure S19. Super-resolution multiplexed renderings of nuclear targets (1-6).**

Super resolution renderings of Exchange-PAINT targets and DNA. Scale bars = 5 µm , 1 µm zoom.

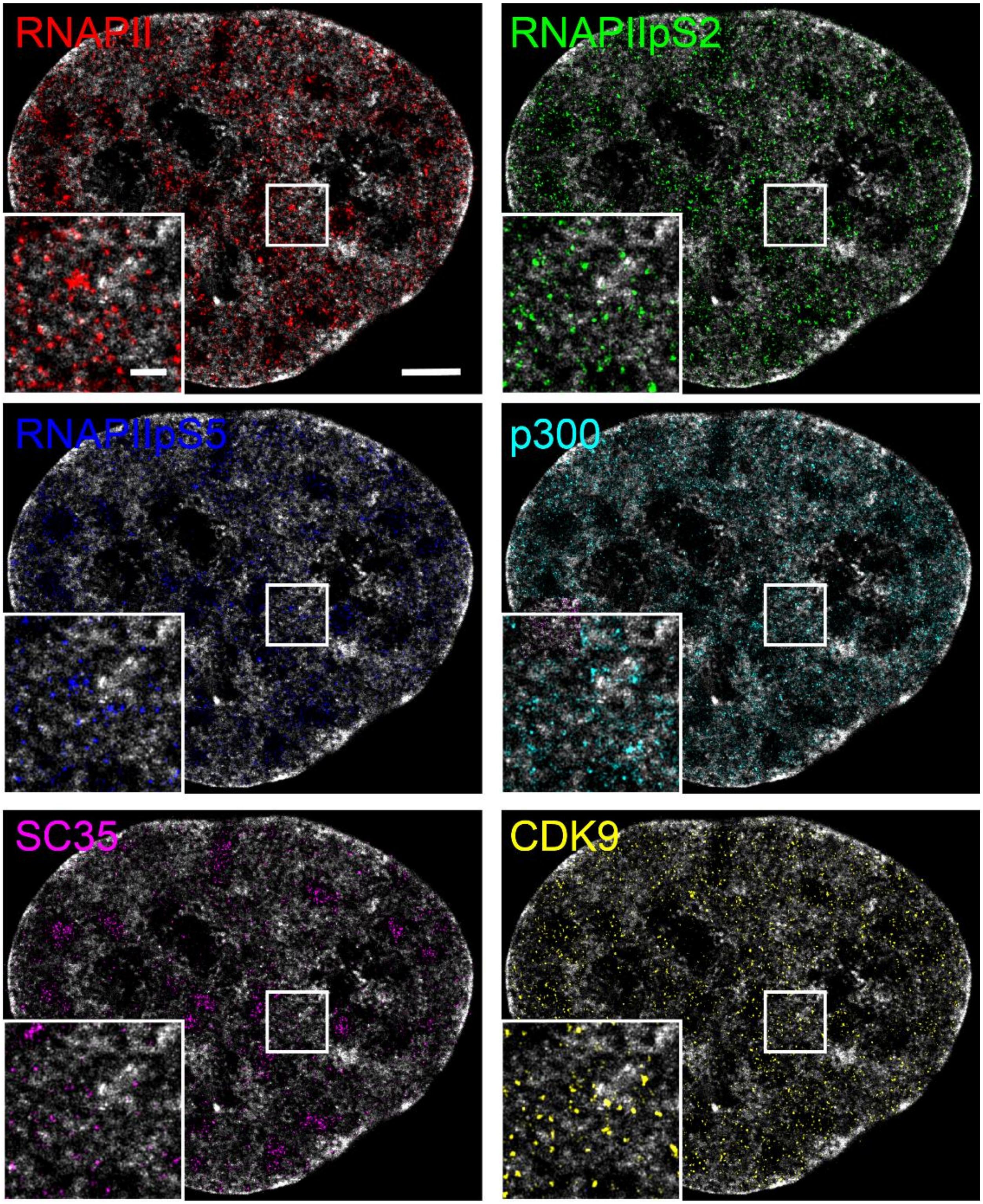

**Figure S20. Super-resolution multiplexed renderings of nuclear targets (7-12).**

Super resolution renderings of Exchange-PAINT targets and DNA. Scale bars = 5 µm , 1 µm zoom.

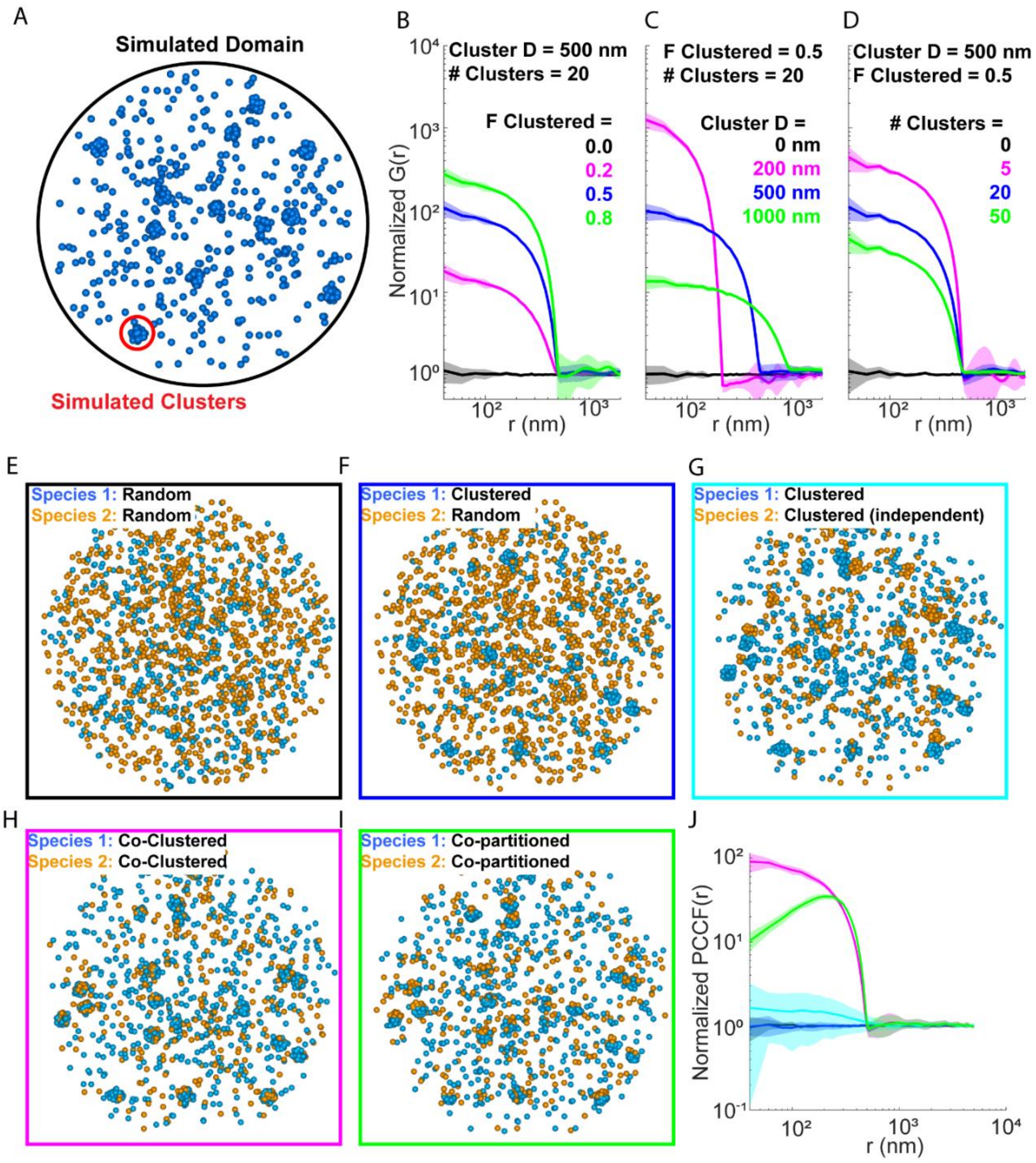

**Figure S21. Pair cross-correlation function (PCCF) simulations.**

A) Representative simulation domain for the single species G(r) calculations with following parameters: Domain Diameter = 10 micron, Fraction Clustered = 0.5, Cluster Diameter = 500 nm, Number of clusters = 20. B-D) Normalized G(r) curves demonstrating the effects of varying simulation parameters for fraction clustered (B), cluster diameter (C), and number of clusters (D). E-I) Representative simulation domains for two species PCCF curves showing two random distributions (E), clustered species 1 and a random species 2 (F), two independently clustered species with randomly localized cluster centroids (G), two co-clustered species with overlapping centroids (H), and two clustered species that share cluster centroids, but partition within different regions of the cluster domain (I). All cases used the same parameters outlined in (A). (J) Normalized PCCF curves for the conditions outlined in E-I.

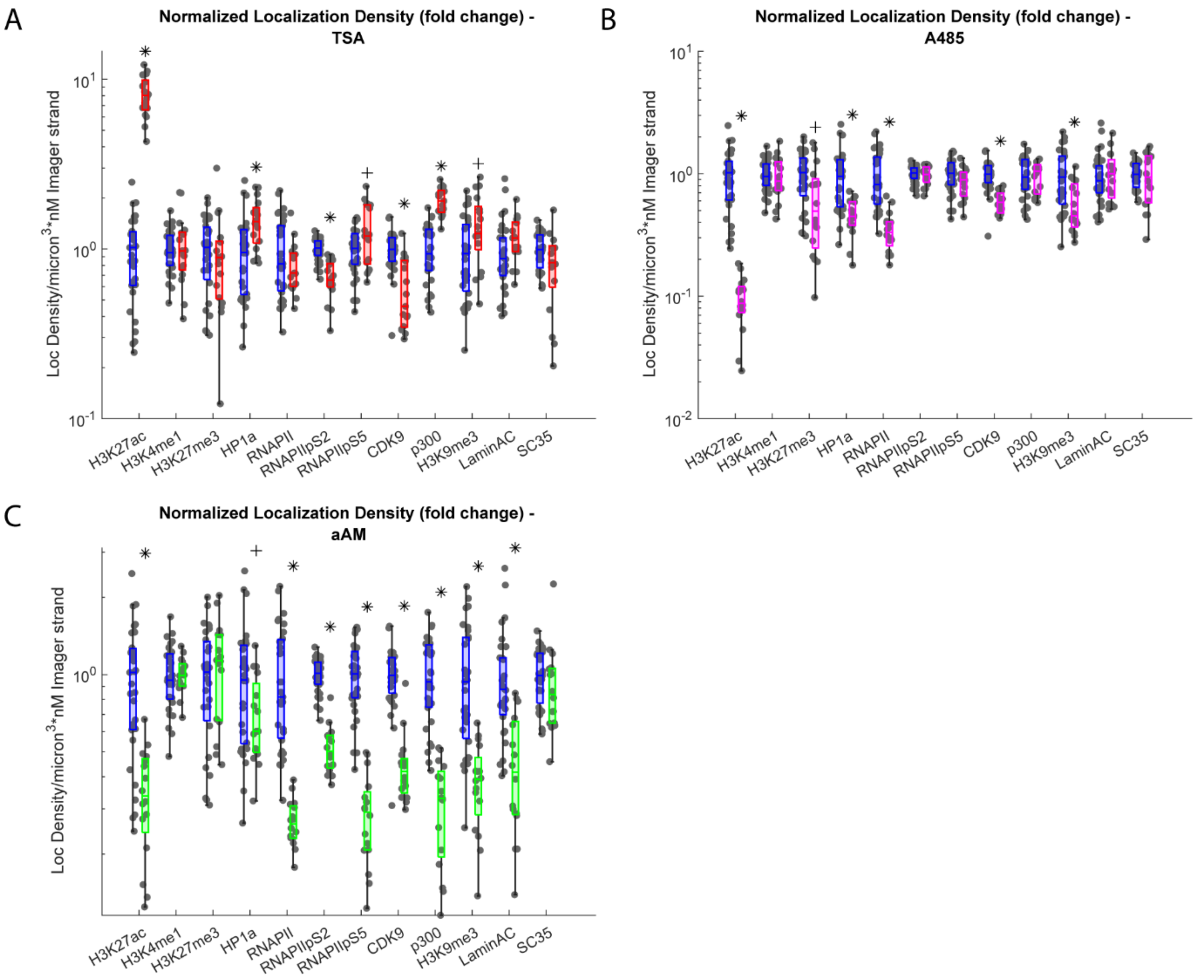

**Figure S22. Localization density changes with perturbations.**

(A) Box plot of localization densities for control vs. TSA datasets inversely scaled by imager strand concentration plotted on log scale (differences represent fold changes, blue indicates control, and red indicates TSA). Points represent n=28 cells across three biological replicates for control and n=17 cells across two biological replicates for TSA. (B) Box plot of localization densities for control vs. A-485 datasets inversely scaled by imager strand concentration plotted on log scale (blue represents control and magenta A-485). Points represent n=28 cells across three biological replicates for control and n=15 cells across two biological replicates for A-485. (C) Box plot of localization densities for control vs. αAM datasets inversely scaled by imager strand concentration plotted on log scale (blue represents control and green αAM). Points represent n=28 cells across three biological replicates for control and n=16 cells across two biological replicates for αAM. * $p \leq 0.01$, + $p \leq 0.05$ via two-sided student's t-test.

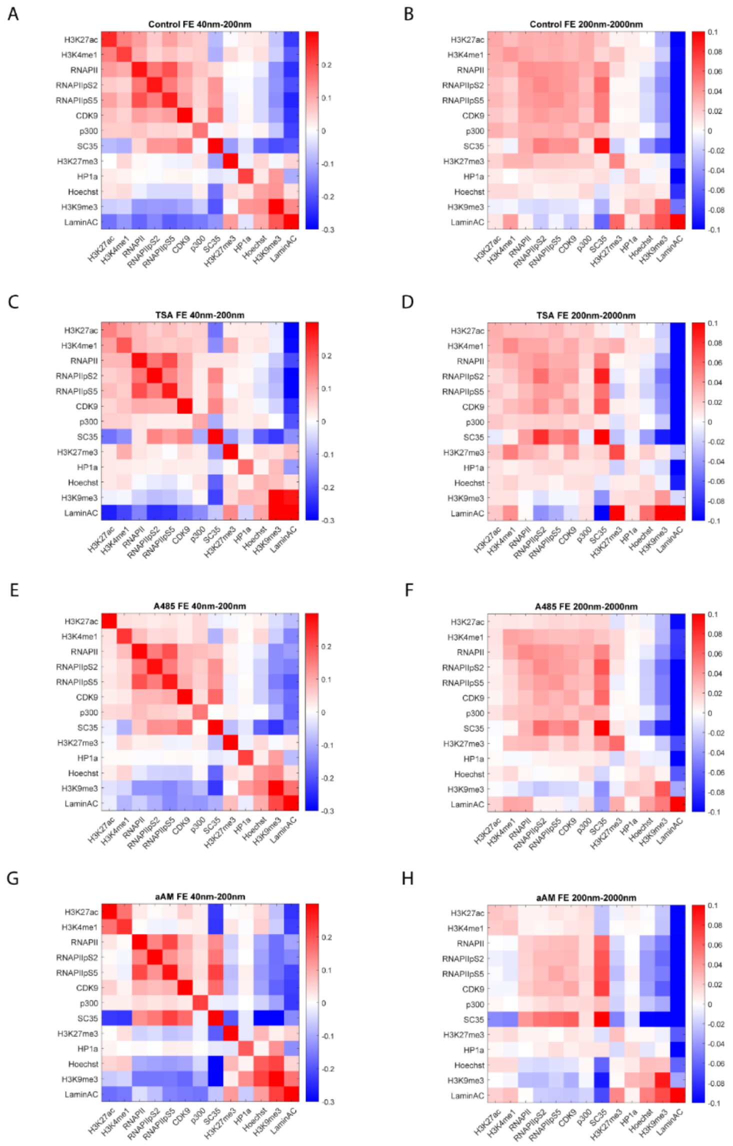

**Figure S23. Heat maps of perturbation effects.**

(A) Heat map of fold enrichment (FE) for control datasets based on normalized PCCF curves versus random within the 40-200 nm length scales. (B) Heat map of fold enrichment (FE) for control datasets based on normalized PCCF curves versus random within the 200-2000 nm length scales. (C) Heat map of fold enrichment (FE) for TSA datasets based on normalized PCCF curves versus random within the 40-200 nm length scales. (D) Heat map of fold enrichment (FE) for TSA datasets based on normalized PCCF curves versus random within the 200-2000 nm length scales. (E) Heat map of fold enrichment (FE) for A-485 datasets based on normalized PCCF curves versus random within the 40-200 nm length scales. (F) Heat map of fold enrichment (FE) for A-485 datasets based on normalized PCCF curves versus random within the 200-2000 nm length scales. (G) Heat map of fold enrichment (FE) for αAM datasets based on normalized PCCF curves versus random within the 40-200 nm length scales. (H) Heat map of fold enrichment (FE) for αAM datasets based on normalized PCCF curves versus random within the 200-2000 nm length scales.

A

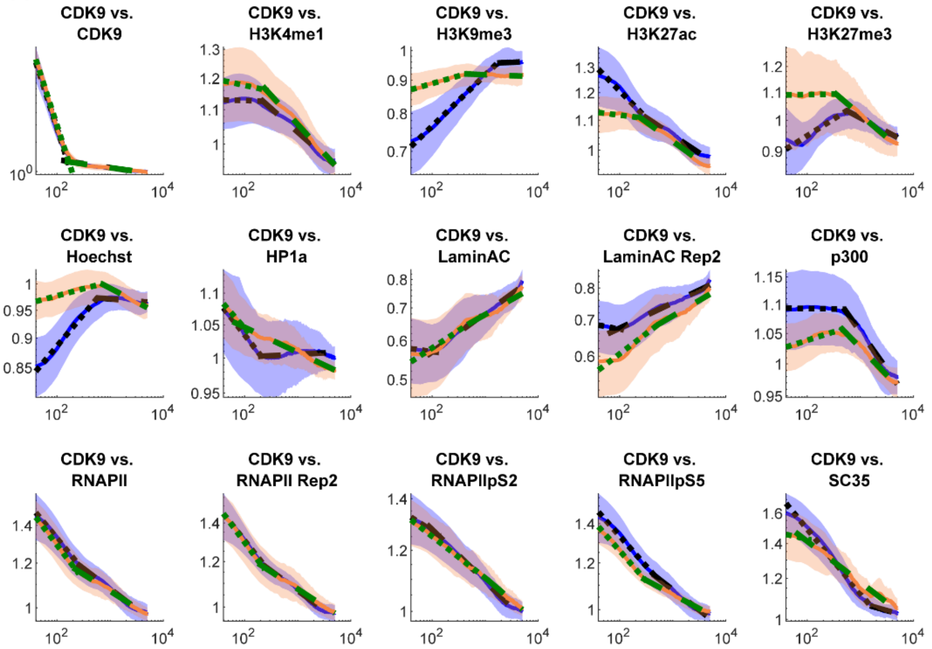

B

**Wilcoxon rank sum control vs. TSA: CDK9 vs.**

| | Hinge | Slope 1 | Slope 2 |
|---|---|---|---|
| **CDK9** | 0.033872 | 0.919528 | 0.028601 |
| **H3K4me1** | 0.081872 | 0.694403 | 0.479445 |
| **H3K9me3** | 0.028601 | 2.10E-05 | 0.018971 |
| **H3K27Ac** | 0.677895 | 5.58E-05 | 0.044502 |
| **H3K27me3** | 0.024052 | 0.000155 | 0.04948 |
| **Hoechst** | 0.094411 | 3.13E-08 | 0.001489 |
| **HP1a** | 0.991044 | 0.21276 | 3.12E-05 |
| **LaminAC** | 0.613461 | 0.247596 | 0.919528 |
| **LaminAC Rep2** | 0.991044 | 0.006377 | 0.597784 |
| **p300** | 0.451993 | 0.001274 | 0.064 |
| **RNAPII** | 0.866284 | 0.18906 | 0.221118 |
| **RNAPII Rep2** | 0.363217 | 0.66154 | 0.955241 |
| **RNAPIIpS2** | 1 | 0.813641 | 0.493505 |
| **RNAPIIpS5** | 0.074289 | 0.727851 | 0.118684 |
| **SC35** | 0.848658 | 0.000129 | 0.451993 |

**Figure S24. CDK9 vs. all, TSA vs. Control PCCF.**

(A) Normalized PCCF curves versus random (blue indicates control, orange indicates TSA). (B) Wilcoxon rank sum p-values for each pair of comparisons.

A

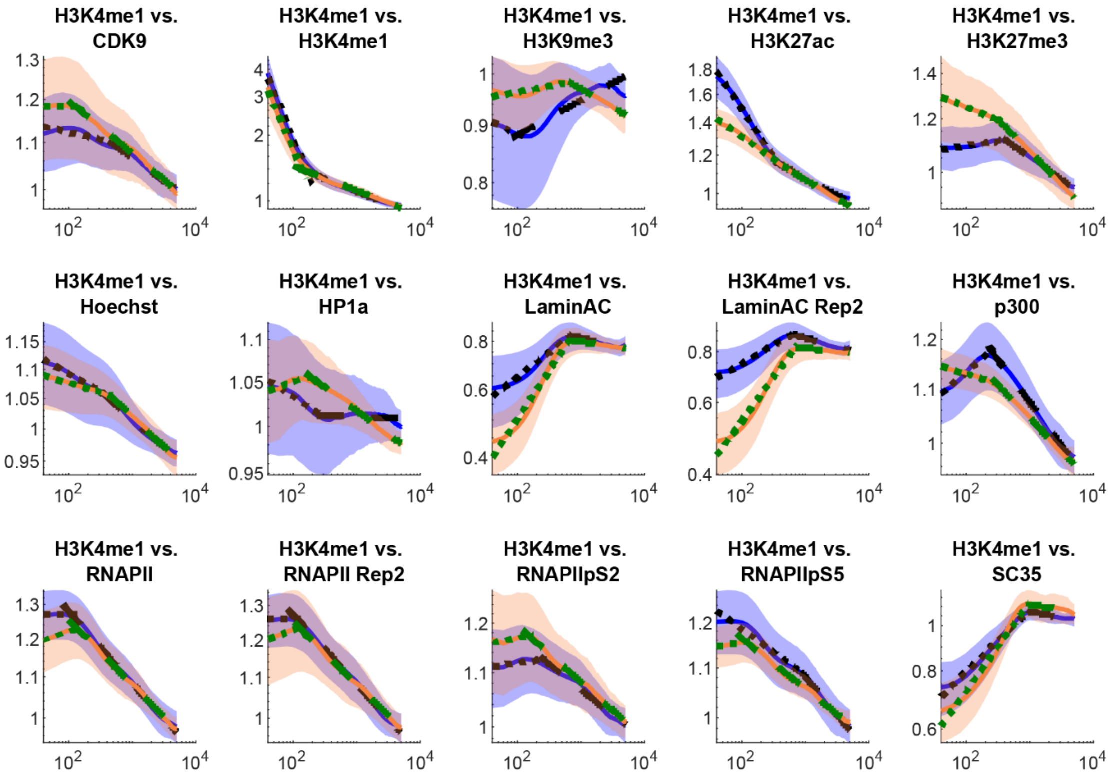

B

**Wilcoxon rank sum control vs. TSA: H3K4me1 vs.**

| | Hinge | Slope 1 | Slope 2 |
|---|---|---|---|
| **CDK9** | 0.0073 | 0.727851 | 0.883977 |
| **H3K4me1** | 0.003392 | 0.465606 | 0.399844 |
| **H3K9me3** | 0.147598 | 0.18161 | 0.042175 |
| **H3K27Ac** | 0.006824 | 3.13E-08 | 0.694403 |
| **H3K27me3** | 0.042175 | 1.40E-07 | 0.005188 |
| **Hoechst** | 0.711058 | 0.005956 | 0.307015 |
| **HP1a** | 0.020145 | 2.83E-05 | 0.002933 |
| **LaminAC** | 0.955241 | 3.55E-06 | 0.479445 |
| **LaminAC Rep2** | 0.21276 | 5.90E-08 | 0.042175 |
| **p300** | 0.081872 | 2.75E-08 | 0.017858 |
| **RNAPII** | 0.328765 | 1.14E-05 | 0.328765 |
| **RNAPII Rep2** | 0.955241 | 1.40E-05 | 0.081872 |
| **RNAPIIpS2** | 0.078006 | 0.727851 | 0.919528 |
| **RNAPIIpS5** | 0.060844 | 0.001274 | 0.129666 |
| **SC35** | 0.991044 | 9.81E-05 | 0.973136 |

**Figure S25. H3K4me1 vs. all, TSA vs. Control PCCF.**

(A) Normalized PCCF curves versus random (blue indicates control, orange indicates TSA). (B) Wilcoxon rank sum p-values for each pair of comparisons.

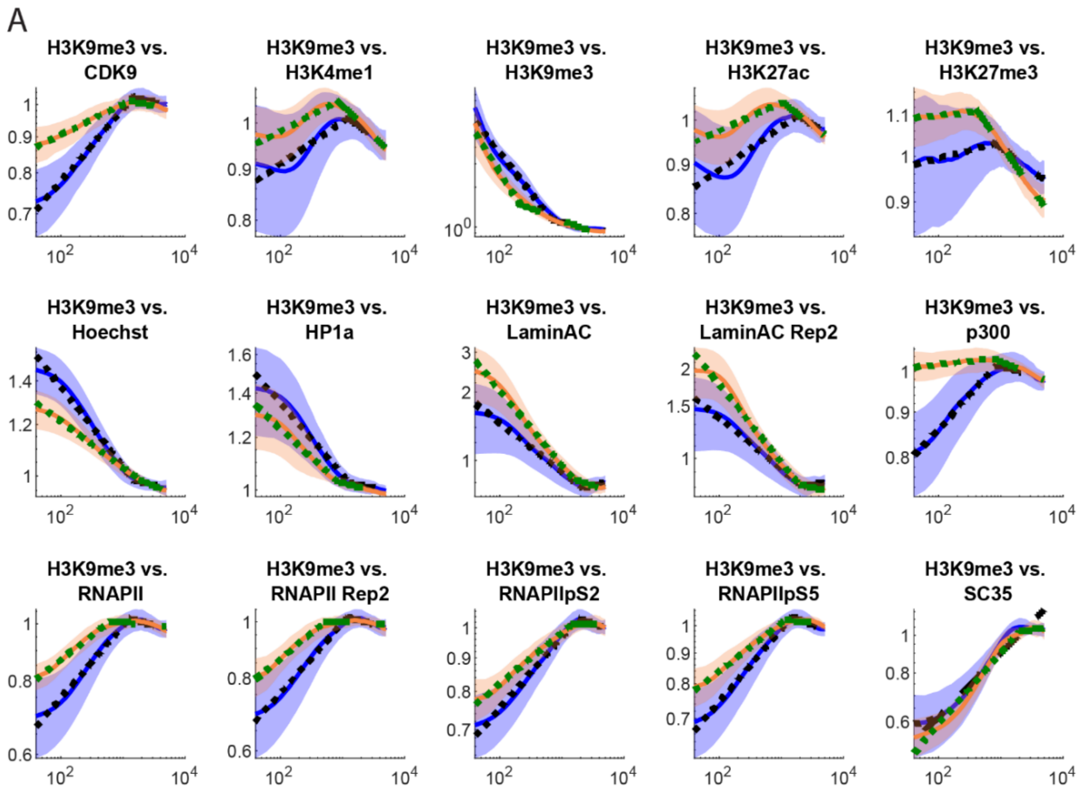

B

**Wilcoxon rank sum control vs. TSA: H3K9me3 vs.**

| | Hinge | Slope 1 | Slope 2 |
|---|---|---|---|
| **CDK9** | 0.085892 | 1.31E-06 | 0.451993 |
| **H3K4me1** | 0.991044 | 0.866284 | 0.042175 |
| **H3K9me3** | 1.71E-05 | 0.167366 | 0.035805 |
| **H3K27Ac** | 0.597784 | 0.266436 | 0.022682 |
| **H3K27me3** | 0.001737 | 0.266436 | 0.001274 |
| **Hoechst** | 0.727851 | 0.000129 | 0.060844 |
| **HP1a** | 0.141421 | 0.296506 | 0.328765 |
| **LaminAC** | 0.307015 | 0.002933 | 0.17438 |
| **LaminAC Rep2** | 0.883977 | 0.098918 | 0.060844 |
| **p300** | 0.919528 | 3.56E-08 | 0.451993 |
| **RNAPII** | 1.84E-06 | 0.000288 | 0.078006 |
| **RNAPII Rep2** | 1.14E-05 | 0.002725 | 0.020145 |
| **RNAPIIpS2** | 0.141421 | 0.033872 | 0.153979 |
| **RNAPIIpS5** | 0.016803 | 0.000288 | 0.008341 |
| **SC35** | 0.296506 | 0.12408 | 0.727851 |

**Figure S26. H3K9me3 vs. all, TSA vs. Control PCCF.**

(A) Normalized PCCF curves versus random (blue indicates control, orange indicates TSA). (B) Wilcoxon rank sum p-values for each pair of comparisons.

A

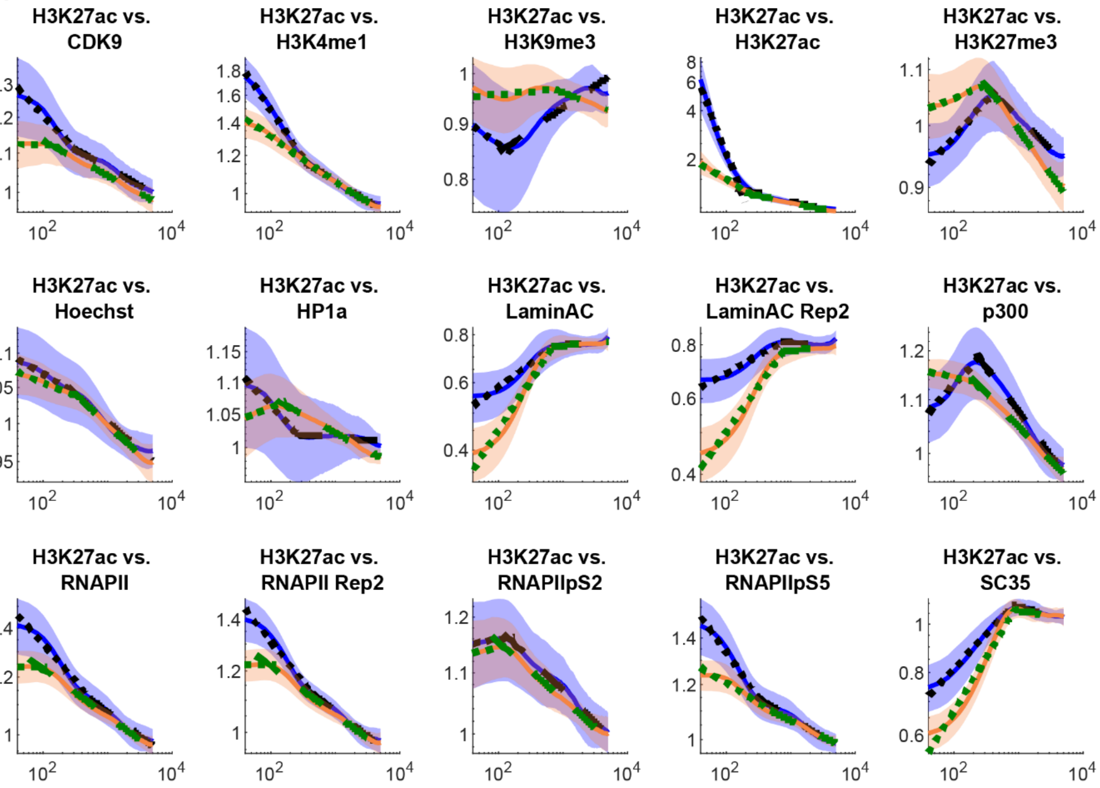

B

**Wilcoxon rank sum control vs. TSA: H3K27ac vs.**

| | Hinge | Slope 1 | Slope 2 |
|---|---|---|---|
| **CDK9** | 0.078006 | 1.26E-05 | 0.438607 |
| **H3K4me1** | 0.425452 | 2.75E-08 | 0.66154 |
| **H3K9me3** | 0.06729 | 0.015802 | 0.000242 |
| **H3K27Ac** | 1.26E-05 | 4.04E-08 | 0.00891 |
| **H3K27me3** | 3.79E-05 | 2.29E-06 | 0.003915 |
| **Hoechst** | 0.551882 | 0.013958 | 0.307015 |
| **HP1a** | 0.03203 | 2.28E-07 | 0.000222 |
| **LaminAC** | 0.052139 | 9.32E-07 | 0.973136 |
| **LaminAC Rep2** | 0.566989 | 5.20E-08 | 0.340006 |
| **p300** | 0.41253 | 2.75E-08 | 0.01083 |
| **RNAPII** | 0.522274 | 1.84E-06 | 0.919528 |
| **RNAPII Rep2** | 0.054917 | 2.28E-07 | 0.711058 |
| **RNAPIIpS2** | 0.566989 | 0.399844 | 0.153979 |
| **RNAPIIpS5** | 0.052139 | 5.88E-07 | 0.039953 |
| **SC35** | 0.582291 | 3.26E-07 | 0.479445 |

**Figure S27. H3K27ac vs. all, TSA vs. Control PCCF.**

(A) Normalized PCCF curves versus random (blue indicates control, orange indicates TSA). (B) Wilcoxon rank sum p-values for each pair of comparisons.

A

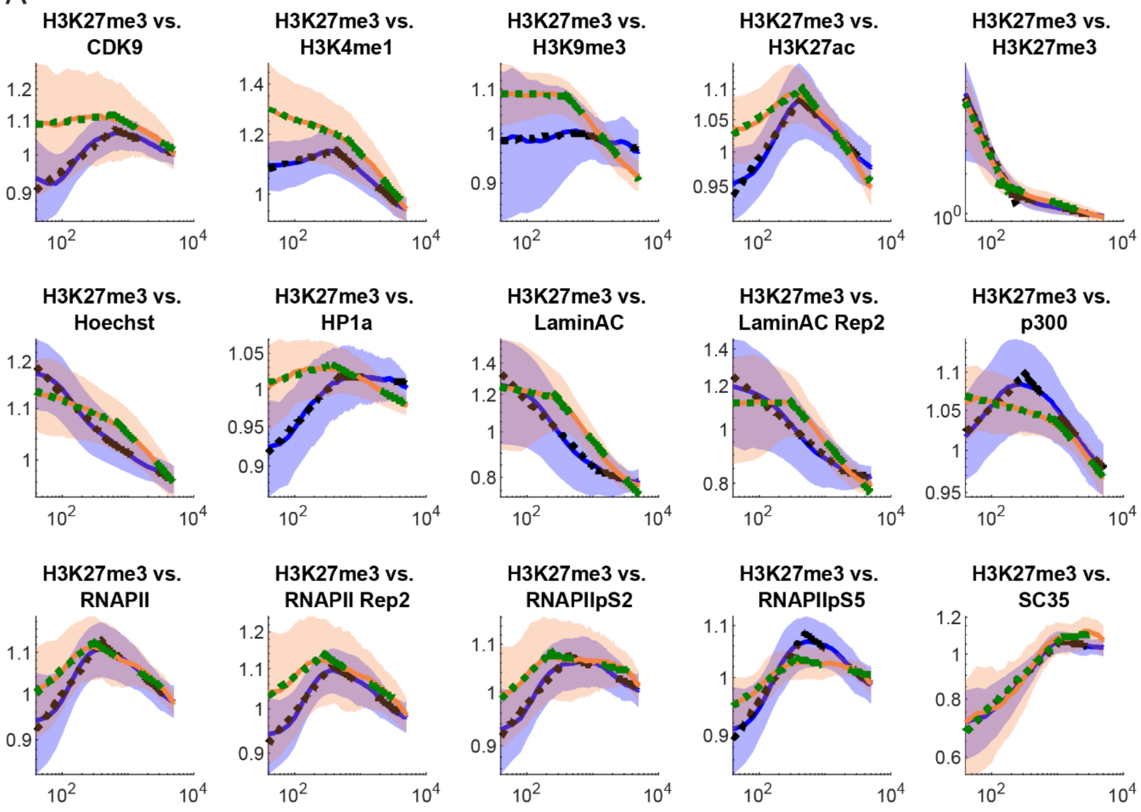

B

**Wilcoxon rank sum control vs. TSA: H3K27me3 vs.**

| | Hinge | Slope 1 | Slope 2 |
|---|---|---|---|
| **CDK9** | 0.566989 | 9.81E-05 | 0.375186 |
| **H3K4me1** | 0.004204 | 5.90E-08 | 0.016803 |
| **H3K9me3** | 0.108445 | 0.507782 | 0.001274 |
| **H3K27Ac** | 0.375186 | 1.04E-06 | 0.016803 |
| **H3K27me3** | 8.31E-07 | 0.221118 | 0.00079 |
| **Hoechst** | 0.522274 | 8.58E-08 | 0.000288 |
| **HP1a** | 0.094411 | 4.91E-06 | 0.009513 |
| **LaminAC** | 0.000288 | 3.19E-06 | 2.32E-05 |
| **LaminAC Rep2** | 0.001178 | 1.03E-05 | 1.71E-05 |
| **p300** | 4.91E-06 | 2.28E-07 | 0.991044 |
| **RNAPII** | 0.020145 | 0.004204 | 0.141421 |
| **RNAPII Rep2** | 0.022682 | 0.010152 | 0.081872 |
| **RNAPIIpS2** | 0.074289 | 0.03203 | 0.094411 |
| **RNAPIIpS5** | 0.074289 | 1.90E-05 | 0.00891 |
| **SC35** | 0.113474 | 0.582291 | 0.247596 |

**Figure S28. H3K27me3 vs. all, TSA vs. Control PCCF.**

(A) Normalized PCCF curves versus random (blue indicates control, orange indicates TSA). (B) Wilcoxon rank sum p-values for each pair of comparisons.

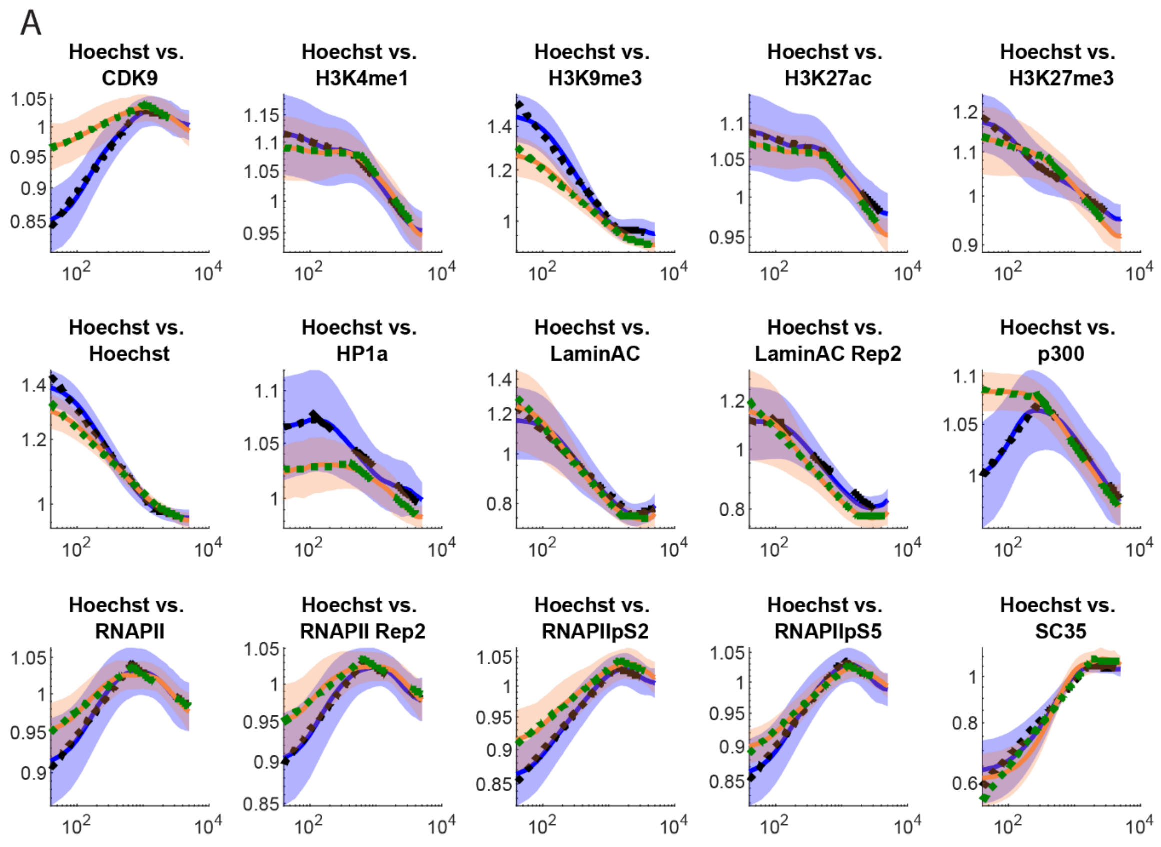

B

**Wilcoxon rank sum control vs. TSA: Hoechst vs.**

| | Hinge | Slope 1 | Slope 2 |
|---|---|---|---|
| **CDK9** | 0.247596 | 3.56E-08 | 0.046935 |
| **H3K4me1** | 0.057817 | 0.016803 | 0.328765 |
| **H3K9me3** | 0.030274 | 0.00079 | 0.276217 |
| **H3K27Ac** | 0.135445 | 0.037831 | 0.090071 |
| **H3K27me3** | 0.937369 | 2.90E-07 | 0.00218 |
| **Hoechst** | 0.566989 | 0.001005 | 0.266436 |
| **HP1a** | 0.375186 | 0.778992 | 0.727851 |
| **LaminAC** | 0.256896 | 0.039953 | 0.901728 |
| **LaminAC Rep2** | 0.955241 | 0.018971 | 0.41253 |
| **p300** | 0.425452 | 2.75E-08 | 0.645345 |
| **RNAPII** | 0.937369 | 6.75E-06 | 0.991044 |
| **RNAPII Rep2** | 0.645345 | 4.61E-05 | 0.778992 |
| **RNAPIIpS2** | 0.399844 | 0.000222 | 0.566989 |
| **RNAPIIpS5** | 0.016803 | 2.05E-06 | 0.21276 |
| **SC35** | 0.813641 | 0.955241 | 0.28624 |

**Figure S29. Hoechst vs. all, TSA vs. Control PCCF.**

(A) Normalized PCCF curves versus random (blue indicates control, orange indicates TSA). (B) Wilcoxon rank sum p-values for each pair of comparisons.

A

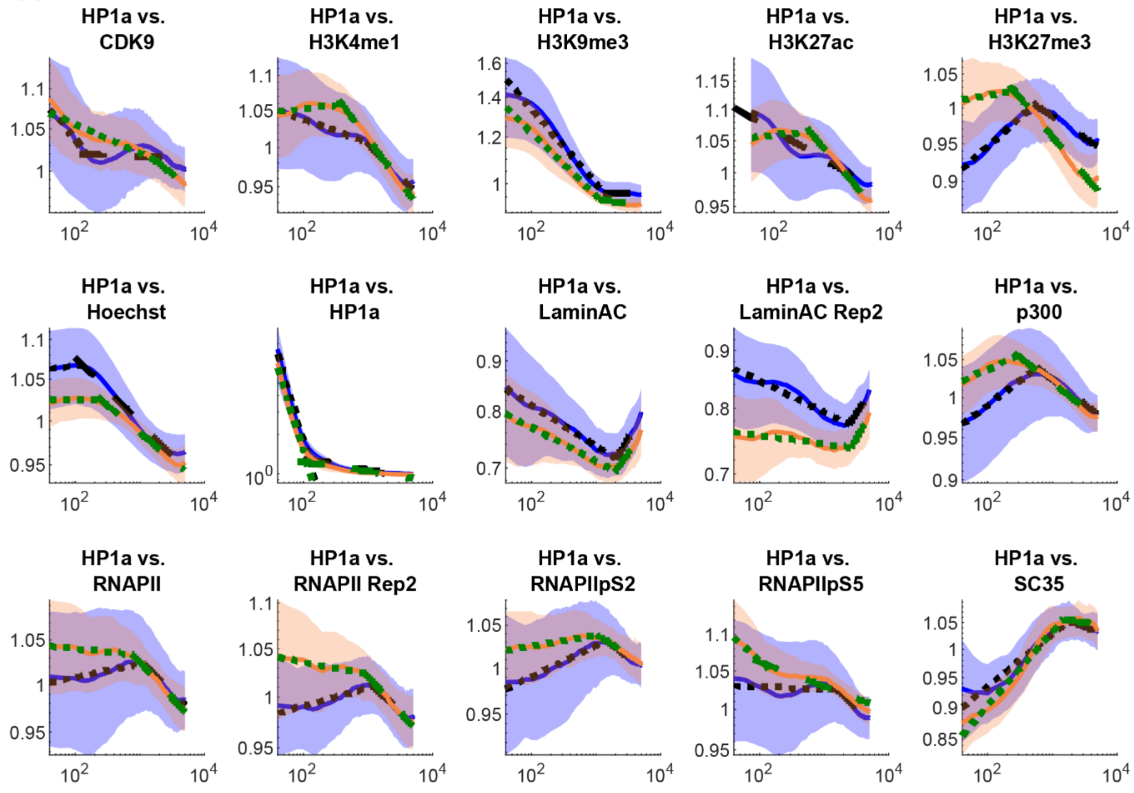

B

**Wilcoxon rank sum control vs. TSA: HP1α vs.**

| | Hinge | Slope 1 | Slope 2 |
|---|---|---|---|
| **CDK9** | 0.694403 | 0.566989 | 0.001005 |
| **H3K4me1** | 0.035805 | 0.000222 | 0.081872 |
| **H3K9me3** | 0.18906 | 0.307015 | 0.078006 |
| **H3K27Ac** | 0.18161 | 2.05E-06 | 0.000373 |
| **H3K27me3** | 6.07E-06 | 0.000373 | 0.002022 |
| **Hoechst** | 0.054917 | 0.711058 | 0.711058 |
| **HP1a** | 0.008341 | 0.919528 | 0.074289 |
| **LaminAC** | 0.507782 | 0.813641 | 0.973136 |
| **LaminAC Rep2** | 0.761826 | 0.057817 | 0.597784 |
| **p300** | 0.000406 | 0.022682 | 0.937369 |
| **RNAPII** | 0.266436 | 0.307015 | 0.129666 |
| **RNAPII Rep2** | 0.108445 | 0.108445 | 0.438607 |
| **RNAPIIpS2** | 0.991044 | 0.012306 | 0.317768 |
| **RNAPIIpS5** | 0.629316 | 0.044502 | 0.438607 |
| **SC35** | 0.060844 | 0.008341 | 0.090071 |

**Figure S30. HP1α vs. all, TSA vs. Control PCCF.**

(A) Normalized PCCF curves versus random (blue indicates control, orange indicates TSA). (B) Wilcoxon rank sum p-values for each pair of comparisons.

A

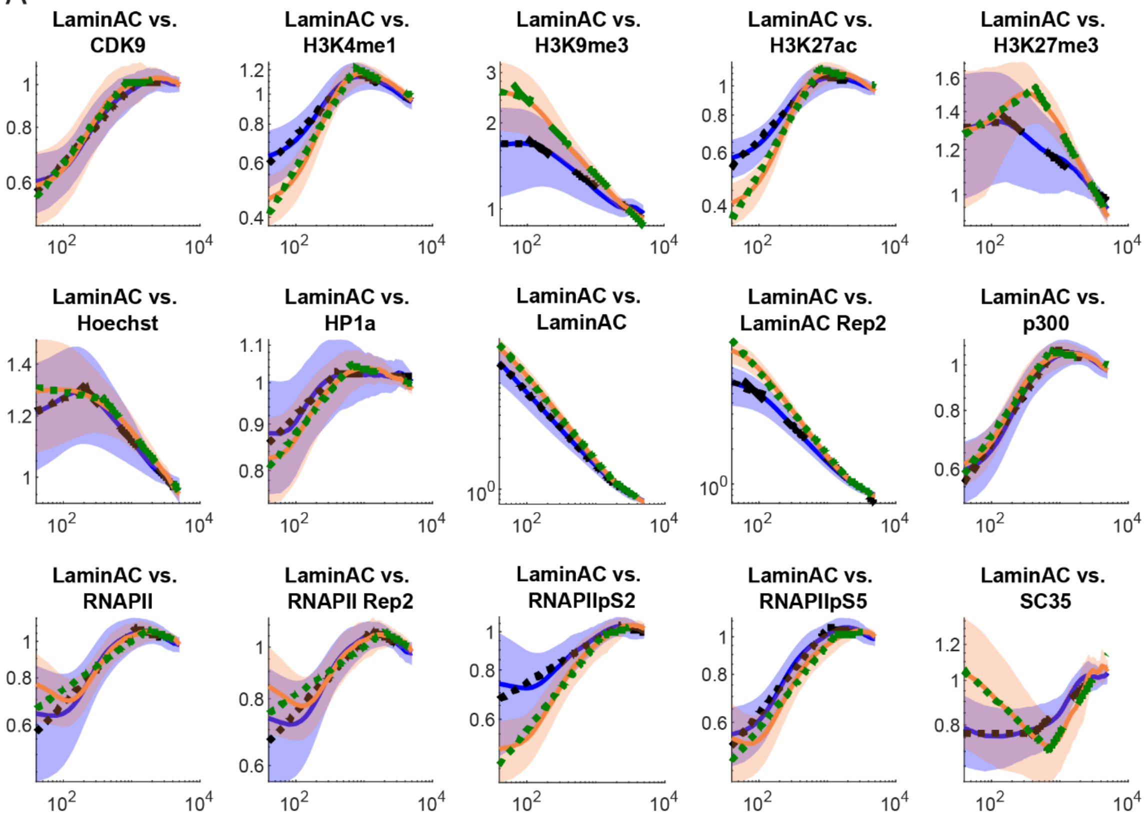

B

**Wilcoxon rank sum control vs. TSA: Lamin A/C vs.**

| | Hinge | Slope 1 | Slope 2 |
|---|---|---|---|
| **CDK9** | 0.340006 | 0.037831 | 0.866284 |
| **H3K4me1** | 0.629316 | 1.55E-05 | 0.761826 |
| **H3K9me3** | 0.256896 | 0.039953 | 0.009513 |
| **H3K27Ac** | 0.694403 | 2.05E-06 | 0.147598 |
| **H3K27me3** | 0.039953 | 0.001737 | 0.000242 |
| **Hoechst** | 0.113474 | 0.0073 | 0.831108 |
| **HP1a** | 0.06729 | 0.21276 | 0.085892 |
| **LaminAC** | 0.004204 | 0.160567 | 0.991044 |
| **LaminAC Rep2** | 0.465606 | 0.007805 | 0.21276 |
| **p300** | 0.042175 | 0.465606 | 0.141421 |
| **RNAPII** | 0.438607 | 0.037831 | 0.35149 |
| **RNAPII Rep2** | 0.796266 | 0.022682 | 0.551882 |
| **RNAPIIpS2** | 0.536975 | 8.94E-05 | 0.387395 |
| **RNAPIIpS5** | 0.108445 | 0.901728 | 0.018971 |
| **SC35** | 0.000442 | 2.56E-06 | 0.000523 |

**Figure S31. Lamin A/C vs. all, TSA vs. Control PCCF.**

(A) Normalized PCCF curves versus random (blue indicates control, orange indicates TSA). (B) Wilcoxon rank sum p-values for each pair of comparisons.

A

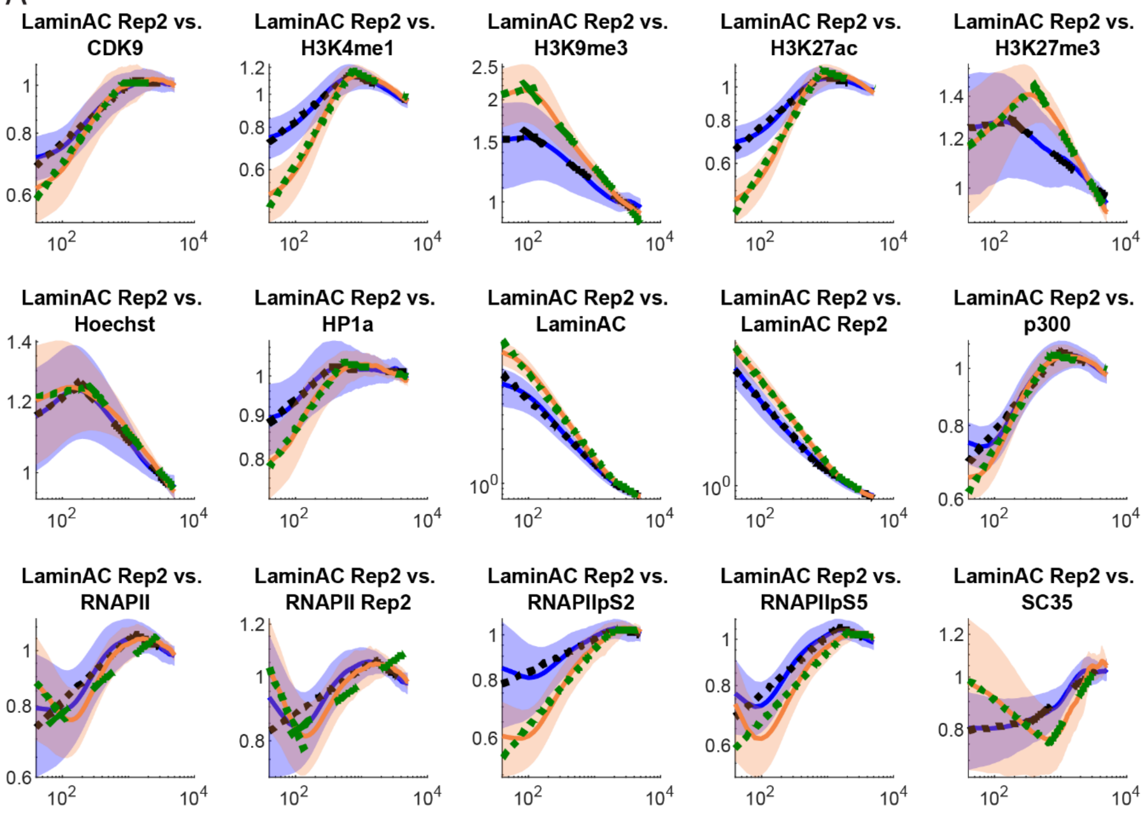

B

**Wilcoxon rank sum control vs. TSA: Lamin A/C repeat vs.**

| | Hinge | Slope 1 | Slope 2 |
|---|---|---|---|
| **CDK9** | 0.266436 | 5.58E-05 | 0.247596 |
| **H3K4me1** | 0.866284 | 2.28E-07 | 0.937369 |
| **H3K9me3** | 0.000671 | 0.238534 | 0.001378 |
| **H3K27Ac** | 0.536975 | 1.59E-07 | 0.317768 |
| **H3K27me3** | 0.070719 | 8.94E-05 | 9.81E-05 |
| **Hoechst** | 0.399844 | 0.057817 | 0.761826 |
| **HP1a** | 0.030274 | 0.003645 | 0.030274 |
| **LaminAC** | 0.937369 | 0.006824 | 0.118684 |
| **LaminAC Rep2** | 0.001089 | 0.41253 | 0.536975 |
| **p300** | 0.000856 | 3.79E-05 | 0.010152 |
| **RNAPII** | 0.522274 | 0.064 | 0.039953 |
| **RNAPII Rep2** | 0.507782 | 0.022682 | 0.018971 |
| **RNAPIIpS2** | 0.848658 | 0.000728 | 0.465606 |
| **RNAPIIpS5** | 0.147598 | 0.438607 | 0.00218 |
| **SC35** | 0.000288 | 7.42E-05 | 8.94E-05 |

**Figure S32. Lamin A/C vs. all, TSA vs. Control PCCF.**

(A) Normalized PCCF curves versus random (blue indicates control, orange indicates TSA). (B) Wilcoxon rank sum p-values for each pair of comparisons.

A

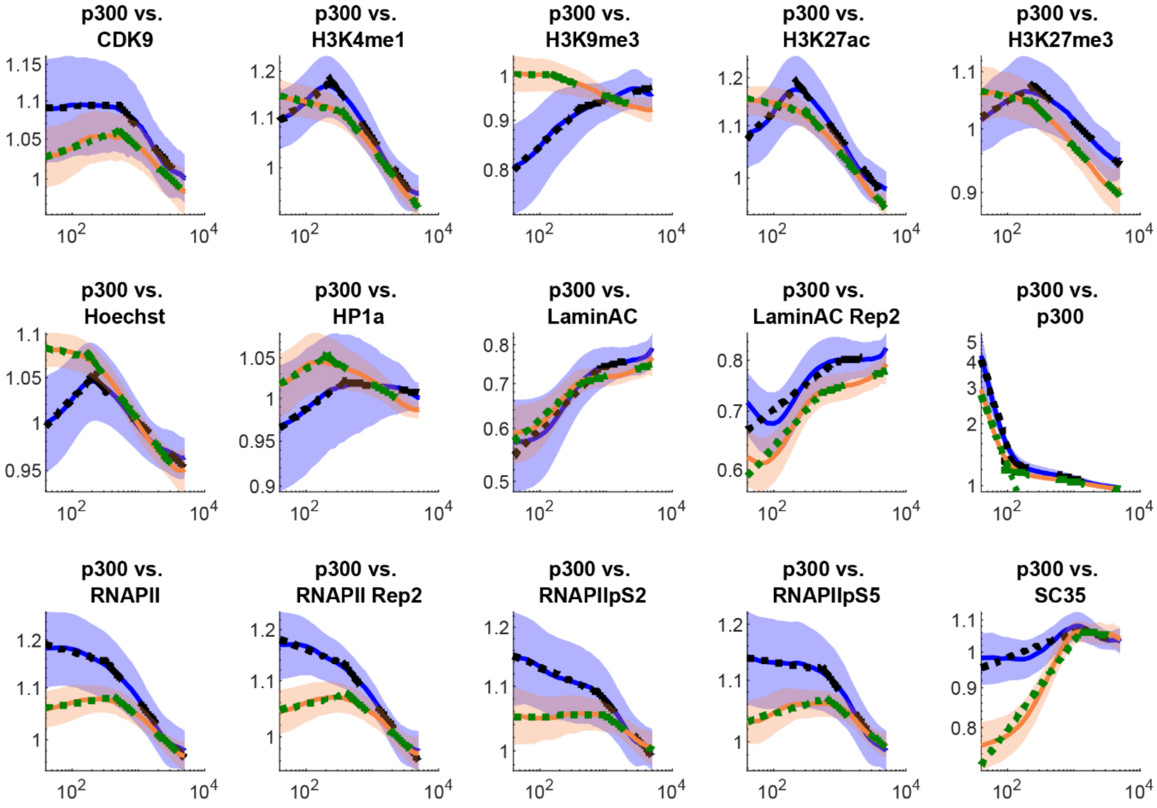

B

**Wilcoxon rank sum control vs. TSA: p300 vs.**

| | Hinge | Slope 1 | Slope 2 |
|---|---|---|---|
| **CDK9** | 0.35149 | 0.011547 | 0.221118 |
| **H3K4me1** | 0.000129 | 2.75E-08 | 0.135445 |
| **H3K9me3** | 0.21276 | 2.75E-08 | 0.033872 |
| **H3K27Ac** | 0.015802 | 2.75E-08 | 0.848658 |
| **H3K27me3** | 0.629316 | 3.26E-07 | 0.054917 |
| **Hoechst** | 0.340006 | 2.75E-08 | 0.012306 |
| **HP1a** | 0.013109 | 0.12408 | 0.003392 |
| **LaminAC** | 0.340006 | 0.21276 | 0.848658 |
| **LaminAC Rep2** | 0.070719 | 0.001005 | 0.18161 |
| **p300** | 7.41E-07 | 0.103594 | 9.81E-05 |
| **RNAPII** | 0.17438 | 5.46E-06 | 0.011547 |
| **RNAPII Rep2** | 0.744776 | 1.64E-06 | 0.002933 |
| **RNAPIIpS2** | 0.387395 | 0.00017 | 0.052139 |
| **RNAPIIpS5** | 0.425452 | 1.03E-05 | 0.001378 |
| **SC35** | 0.522274 | 2.75E-08 | 0.991044 |

**Figure S33. p300 vs. all, TSA vs. Control PCCF.**

(A) Normalized PCCF curves versus random (blue indicates control, orange indicates TSA). (B) Wilcoxon rank sum p-values for each pair of comparisons.

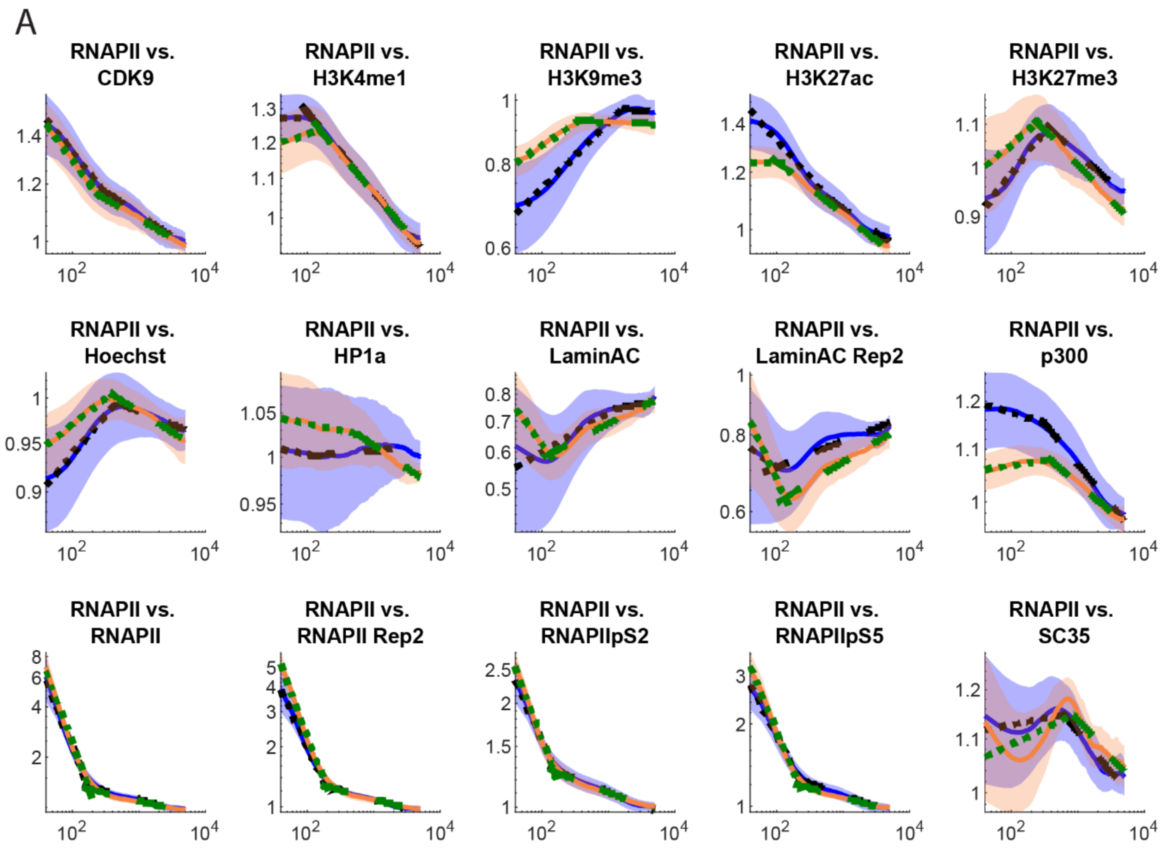

B

**Wilcoxon rank sum control vs. TSA: RNAPII vs.**

| | Hinge | Slope 1 | Slope 2 |
|---|---|---|---|
| **CDK9** | 0.796266 | 0.18906 | 0.340006 |
| **H3K4me1** | 0.03203 | 2.10E-05 | 0.479445 |
| **H3K9me3** | 0.00079 | 0.015802 | 0.536975 |
| **H3K27Ac** | 2.56E-06 | 1.10E-07 | 0.001609 |
| **H3K27me3** | 0.000203 | 0.008341 | 0.094411 |
| **Hoechst** | 0.256896 | 0.00017 | 0.507782 |
| **HP1a** | 0.796266 | 0.991044 | 0.00235 |
| **LaminAC** | 0.025494 | 0.000142 | 0.020145 |
| **LaminAC Rep2** | 0.465606 | 8.33E-06 | 0.003392 |
| **p300** | 0.238534 | 3.44E-05 | 0.009513 |
| **RNAPII** | 0.479445 | 0.17438 | 0.024052 |
| **RNAPII Rep2** | 0.973136 | 0.000728 | 0.238534 |
| **RNAPIIpS2** | 0.276217 | 0.00556 | 0.147598 |
| **RNAPIIpS5** | 0.238534 | 0.00017 | 0.003392 |
| **SC35** | 0.025494 | 0.629316 | 0.937369 |

**Figure S34. RNAPII vs. all, TSA vs. Control PCCF.**

(A) Normalized PCCF curves versus random (blue indicates control, orange indicates TSA). (B) Wilcoxon rank sum p-values for each pair of comparisons.

A

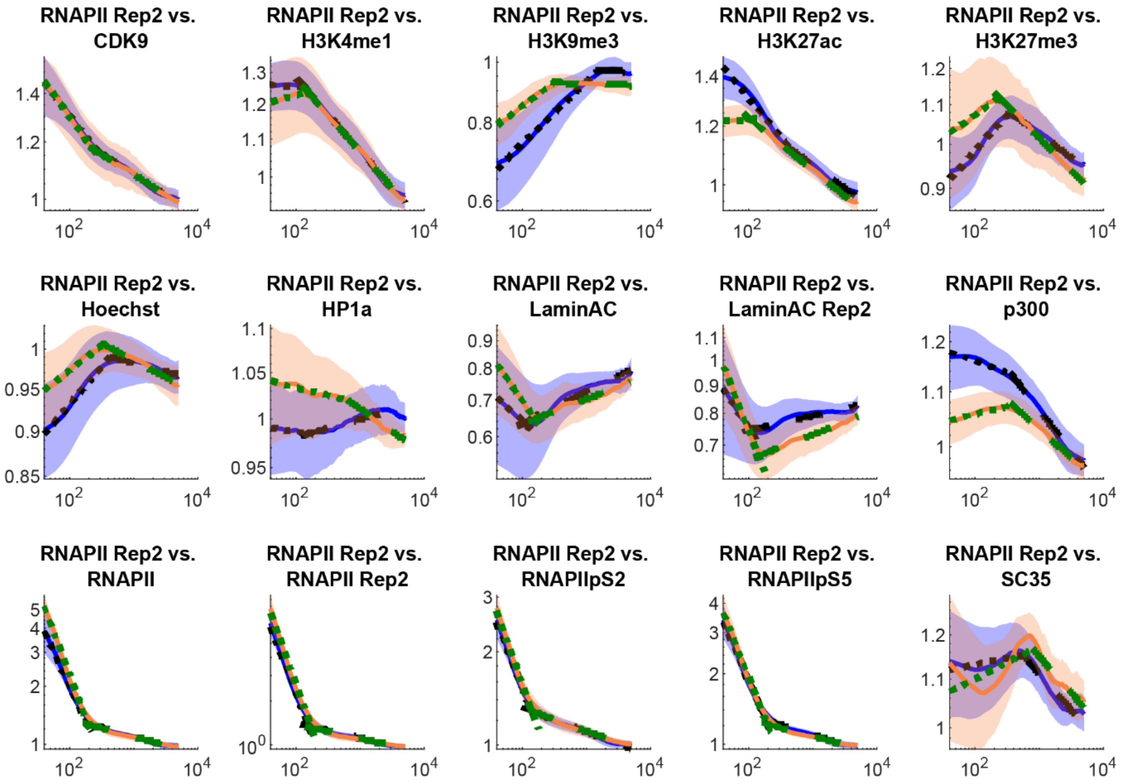

B

**Wilcoxon rank sum control vs. TSA: RNAPII repeat vs.**

| | Hinge | Slope 1 | Slope 2 |
|---|---|---|---|
| **CDK9** | 0.493505 | 0.522274 | 0.66154 |
| **H3K4me1** | 0.081872 | 3.12E-05 | 0.493505 |
| **H3K9me3** | 0.000264 | 0.025494 | 0.465606 |
| **H3K27Ac** | 9.25E-06 | 1.59E-07 | 0.006377 |
| **H3K27me3** | 2.32E-05 | 0.025494 | 0.030274 |
| **Hoechst** | 0.074289 | 5.58E-05 | 0.238534 |
| **HP1a** | 0.296506 | 0.866284 | 0.002933 |
| **LaminAC** | 0.375186 | 0.00891 | 0.052139 |
| **LaminAC Rep2** | 0.677895 | 0.00556 | 0.003155 |
| **p300** | 0.991044 | 6.07E-06 | 0.0073 |
| **RNAPII** | 1 | 0.000569 | 0.399844 |
| **RNAPII Rep2** | 0.007805 | 0.18906 | 0.094411 |
| **RNAPIIpS2** | 0.12408 | 0.28624 | 0.340006 |
| **RNAPIIpS5** | 0.052139 | 0.221118 | 0.003155 |
| **SC35** | 0.003645 | 0.167366 | 0.566989 |

**Figure S35. RNAPII repeat vs. all, TSA vs. Control PCCF.**

(A) Normalized PCCF curves versus random (blue indicates control, orange indicates TSA). (B) Wilcoxon rank sum p-values for each pair of comparisons.

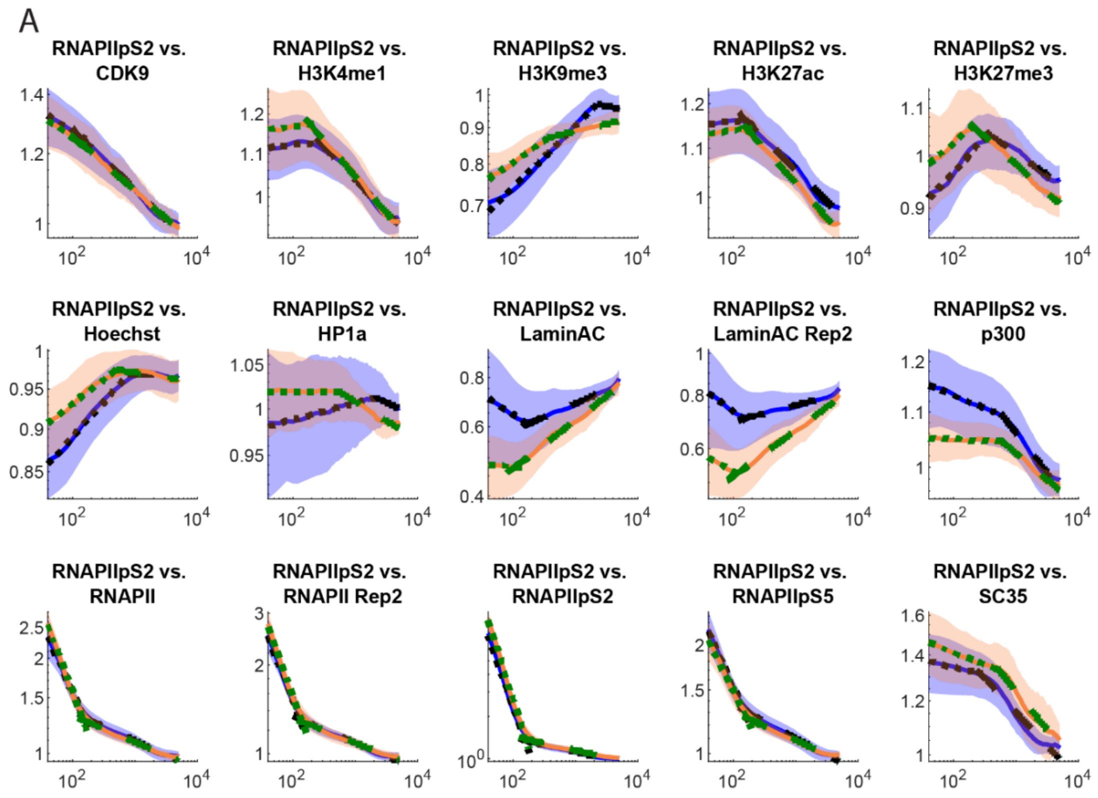

B

**Wilcoxon rank sum control vs. TSA: RNAPII pS2 vs.**

| | Hinge | Slope 1 | Slope 2 |
|---|---|---|---|
| **CDK9** | 0.694403 | 0.66154 | 0.582291 |
| **H3K4me1** | 0.06729 | 0.778992 | 0.363217 |
| **H3K9me3** | 0.015802 | 0.094411 | 0.761826 |
| **H3K27Ac** | 0.761826 | 0.438607 | 0.317768 |
| **H3K27me3** | 0.000155 | 0.135445 | 0.221118 |
| **Hoechst** | 0.147598 | 8.94E-05 | 0.727851 |
| **HP1a** | 0.744776 | 0.387395 | 0.000928 |
| **LaminAC** | 0.566989 | 0.002531 | 0.030274 |
| **LaminAC Rep2** | 0.118684 | 0.247596 | 4.61E-05 |
| **p300** | 0.937369 | 0.00079 | 0.012306 |
| **RNAPII** | 0.399844 | 0.004511 | 0.21276 |
| **RNAPII Rep2** | 0.135445 | 0.276217 | 0.307015 |
| **RNAPIIpS2** | 0.052139 | 0.021381 | 0.744776 |
| **RNAPIIpS5** | 0.060844 | 0.141421 | 0.009513 |
| **SC35** | 0.03203 | 0.221118 | 0.108445 |

**Figure S36. RNAPII pS2 vs. all, TSA vs. Control PCCF.**

(A) Normalized PCCF curves versus random (blue indicates control, orange indicates TSA). (B) Wilcoxon rank sum p-values for each pair of comparisons.

A

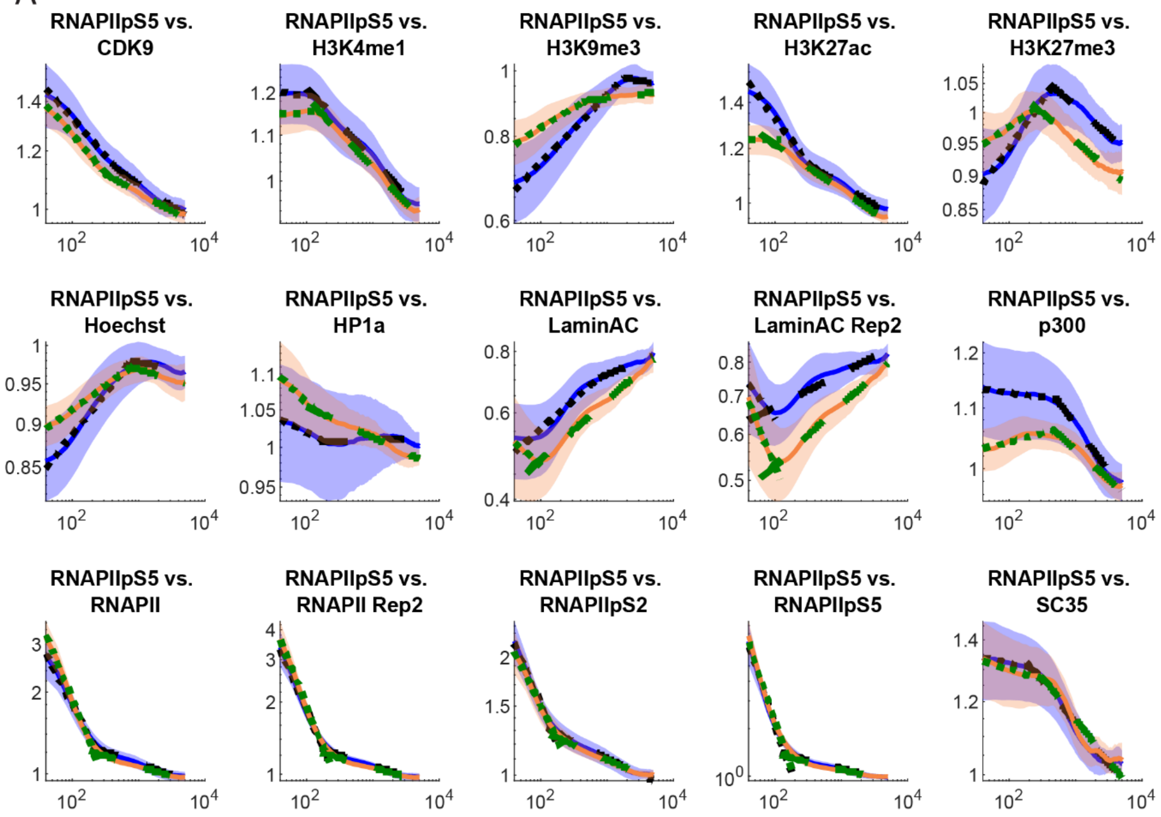

B

**Wilcoxon rank sum control vs. TSA: RNAPII pS5 vs.**

| | Hinge | Slope 1 | Slope 2 |
|---|---|---|---|
| CDK9 | 0.363217 | 0.694403 | 0.221118 |
| H3K4me1 | 0.035805 | 0.000129 | 0.160567 |
| H3K9me3 | 0.113474 | 0.002725 | 0.479445 |
| H3K27Ac | 0.451993 | 3.67E-07 | 0.118684 |
| H3K27me3 | 1.64E-06 | 0.000118 | 0.761826 |
| Hoechst | 0.973136 | 6.60E-07 | 0.66154 |
| HP1a | 0.057817 | 0.025494 | 0.000342 |
| LaminAC | 0.013958 | 0.010152 | 0.000242 |
| LaminAC Rep2 | 0.238534 | 0.001178 | 1.55E-05 |
| p300 | 0.28624 | 7.41E-07 | 0.002531 |
| RNAPII | 0.247596 | 0.000185 | 0.015802 |
| RNAPII Rep2 | 0.035805 | 0.196734 | 0.011547 |
| RNAPIIpS2 | 0.064 | 0.12408 | 0.042175 |
| RNAPIIpS5 | 0.196734 | 0.566989 | 0.033872 |
| SC35 | 0.094411 | 0.711058 | 0.566989 |

**Figure S37. RNAPII pS5 vs. all, TSA vs. Control PCCF.**

(A) Normalized PCCF curves versus random (blue indicates control, orange indicates TSA). (B) Wilcoxon rank sum p-values for each pair of comparisons.

A

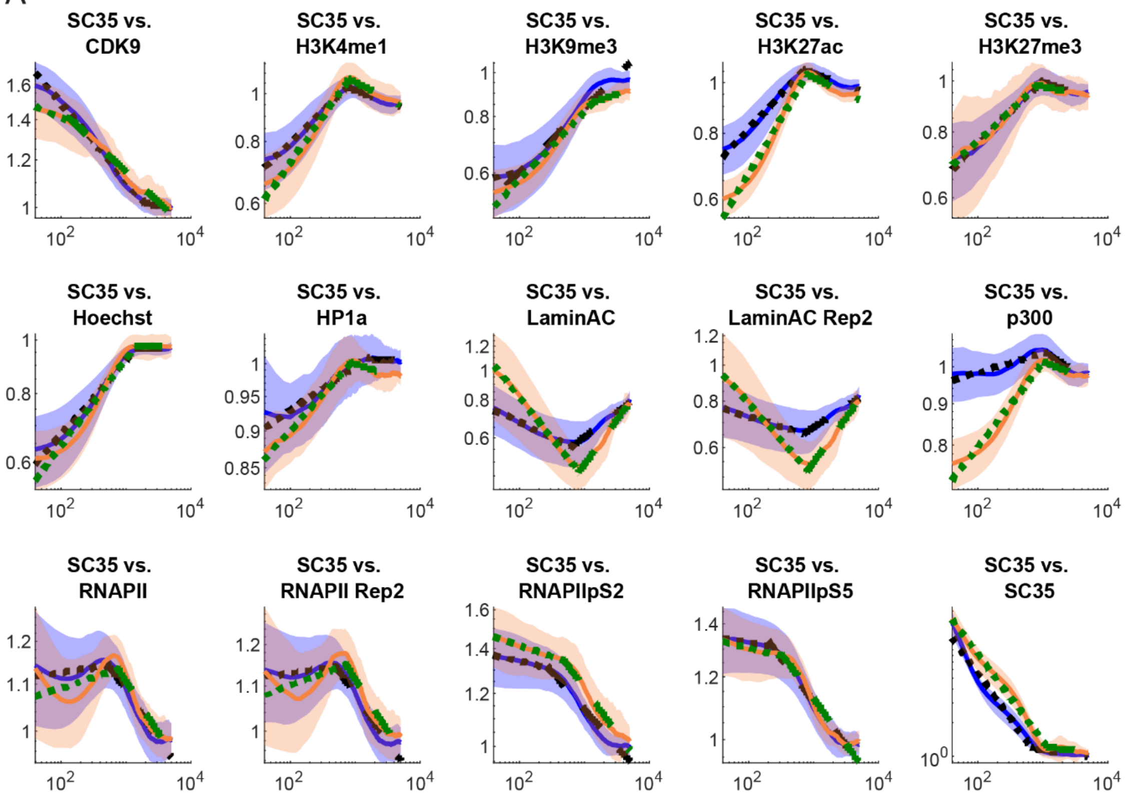

B

**Wilcoxon rank sum control vs. TSA: SC-35 vs.**

| | Hinge | Slope 1 | Slope 2 |
|---|---|---|---|
| **CDK9** | 0.044502 | 0.000288 | 0.042175 |
| **H3K4me1** | 0.196734 | 1.71E-05 | 0.064 |
| **H3K9me3** | 0.493505 | 0.014855 | 0.451993 |
| **H3K27Ac** | 0.118684 | 9.71E-08 | 0.090071 |
| **H3K27me3** | 0.094411 | 0.866284 | 0.147598 |
| **Hoechst** | 0.831108 | 0.451993 | 0.955241 |
| **HP1a** | 0.901728 | 0.003915 | 0.044502 |
| **LaminAC** | 0.003155 | 1.79E-07 | 0.000242 |
| **LaminAC Rep2** | 0.108445 | 3.67E-07 | 0.000155 |
| **p300** | 0.937369 | 2.75E-08 | 0.108445 |
| **RNAPII** | 0.000523 | 0.796266 | 0.507782 |
| **RNAPII Rep2** | 0.000856 | 0.276217 | 0.677895 |
| **RNAPIIpS2** | 0.012306 | 0.167366 | 0.044502 |
| **RNAPIIpS5** | 0.015802 | 0.831108 | 0.465606 |
| **SC35** | 2.10E-05 | 0.399844 | 0.387395 |

**Figure S38. SC-35 vs. all, TSA vs. Control PCCF.**

(A) Normalized PCCF curves versus random (blue indicates control, orange indicates TSA). (B) Wilcoxon rank sum p-values for each pair of comparisons.

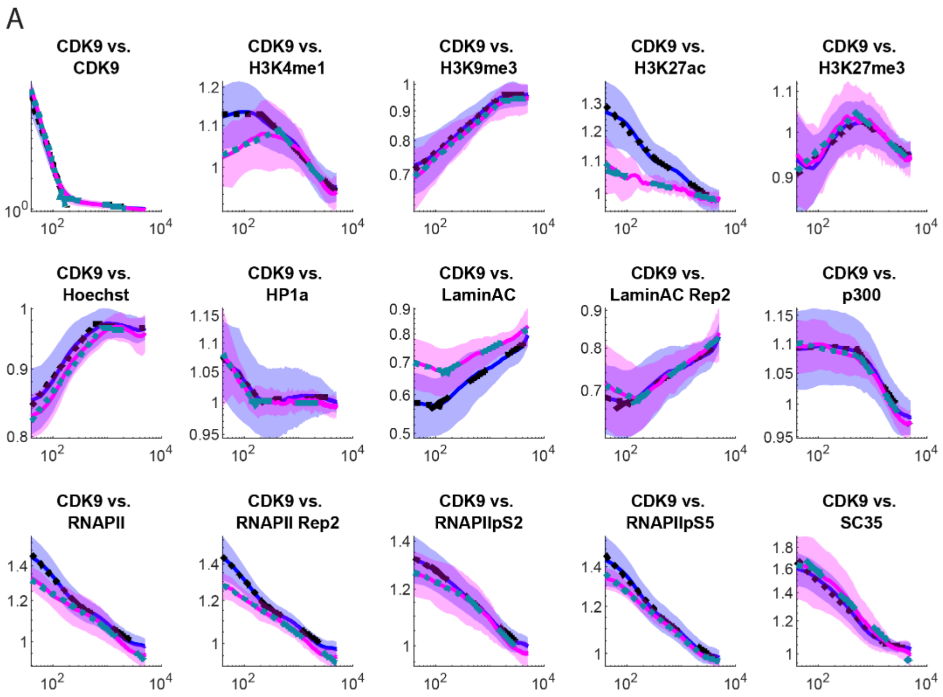

B

**Wilcoxon rank sum control vs. A485: CDK9 vs.**

| | Hinge | Slope 1 | Slope 2 |
|---|---|---|---|
| **CDK9** | 0.307015 | 0.000373 | 0.18161 |
| **H3K4me1** | 0.919528 | 0.000222 | 0.328765 |
| **H3K9me3** | 0.438607 | 0.296506 | 0.991044 |
| **H3K27Ac** | 0.597784 | 0.153979 | 0.000129 |
| **H3K27me3** | 0.387395 | 0.919528 | 0.536975 |
| **Hoechst** | 0.035805 | 0.694403 | 0.866284 |
| **HP1a** | 0.221118 | 0.973136 | 0.28624 |
| **LaminAC** | 0.340006 | 0.017858 | 0.761826 |
| **LaminAC Rep2** | 0.451993 | 0.085892 | 0.694403 |
| **p300** | 0.451993 | 0.108445 | 0.375186 |
| **RNAPII** | 0.41253 | 0.00891 | 0.831108 |
| **RNAPII Rep2** | 0.937369 | 0.000373 | 0.18906 |
| **RNAPIIpS2** | 0.296506 | 0.052139 | 0.375186 |
| **RNAPIIpS5** | 0.866284 | 0.002531 | 0.551882 |
| **SC35** | 0.831108 | 0.167366 | 0.479445 |

**Figure S39. CDK9 vs. all, A-485 vs. Control PCCF.**

(A) Normalized PCCF curves versus random (blue indicates control, magenta indicates A-485). (B) Wilcoxon rank sum p-values for each pair of comparisons.

A

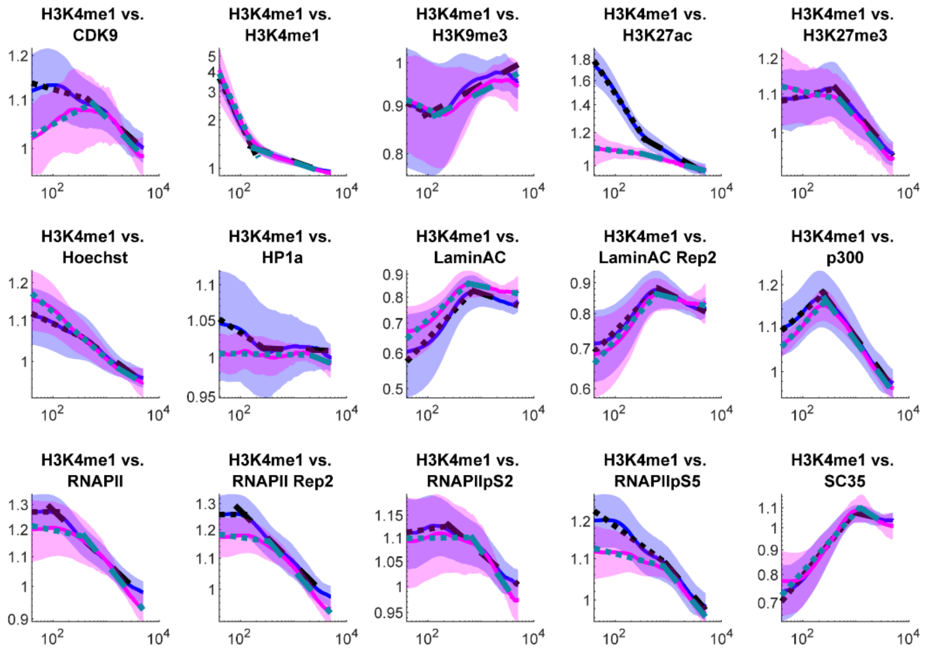

B

Wilcoxon rank sum control vs. A485: H3K4me1 vs.

| | Hinge | Slope 1 | Slope 2 |
|---|---|---|---|
| CDK9 | 0.66154 | 0.000288 | 0.582291 |
| H3K4me1 | 0.004204 | 0.536975 | 0.375186 |
| H3K9me3 | 0.41253 | 0.129666 | 0.081872 |
| H3K27Ac | 0.883977 | 1.17E-06 | 4.41E-06 |
| H3K27me3 | 0.057817 | 0.002531 | 0.479445 |
| Hoechst | 0.629316 | 0.01083 | 0.317768 |
| HP1a | 0.831108 | 0.001609 | 0.479445 |
| LaminAC | 0.296506 | 0.18161 | 0.866284 |
| LaminAC Rep2 | 0.451993 | 0.796266 | 0.340006 |
| p300 | 0.039953 | 0.507782 | 0.536975 |
| RNAPII | 0.001005 | 0.536975 | 0.085892 |
| RNAPII Rep2 | 0.00556 | 0.973136 | 0.094411 |
| RNAPIIpS2 | 0.18161 | 0.41253 | 0.113474 |
| RNAPIIpS5 | 0.866284 | 0.0073 | 0.17438 |
| SC35 | 0.003645 | 0.238534 | 0.022682 |

**Figure S40. H3K4me1 vs. all, A-485 vs. Control PCCF.**

(A) Normalized PCCF curves versus random (blue indicates control, magenta indicates A-485).
(B) Wilcoxon rank sum p-values for each pair of comparisons.

A

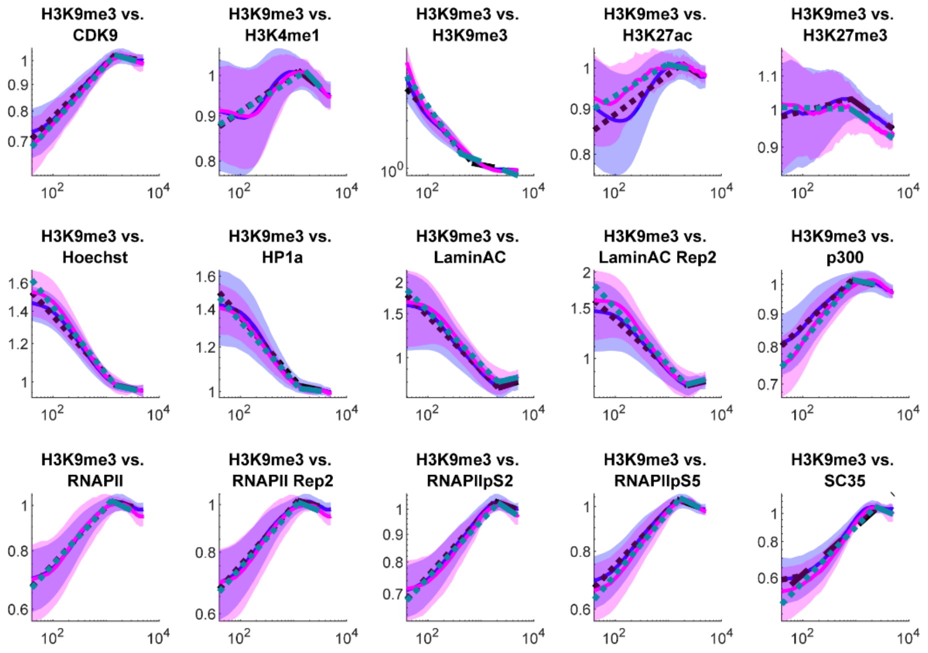

B

**Wilcoxon rank sum control vs. A485: H3K9me3 vs.**

| | Hinge | Slope 1 | Slope 2 |
|---|---|---|---|
| **CDK9** | 0.831108 | 0.21276 | 0.831108 |
| **H3K4me1** | 0.883977 | 0.167366 | 0.937369 |
| **H3K9me3** | 0.074289 | 0.006824 | 0.196734 |
| **H3K27Ac** | 0.094411 | 0.153979 | 0.507782 |
| **H3K27me3** | 0.054917 | 0.085892 | 0.582291 |
| **Hoechst** | 0.744776 | 0.238534 | 0.597784 |
| **HP1a** | 0.597784 | 0.479445 | 0.613461 |
| **LaminAC** | 0.629316 | 0.973136 | 0.328765 |
| **LaminAC Rep2** | 0.582291 | 0.883977 | 0.727851 |
| **p300** | 0.629316 | 0.013958 | 0.778992 |
| **RNAPII** | 0.727851 | 0.629316 | 0.024052 |
| **RNAPII Rep2** | 0.677895 | 0.727851 | 0.027009 |
| **RNAPIIpS2** | 0.465606 | 0.425452 | 0.438607 |
| **RNAPIIpS5** | 0.247596 | 0.276217 | 0.18161 |
| **SC35** | 0.229708 | 0.238534 | 0.363217 |

**Figure S41. H3K9me3 vs. all, A-485 vs. Control PCCF.**

(A) Normalized PCCF curves versus random (blue indicates control, magenta indicates A-485).
(B) Wilcoxon rank sum p-values for each pair of comparisons.

A

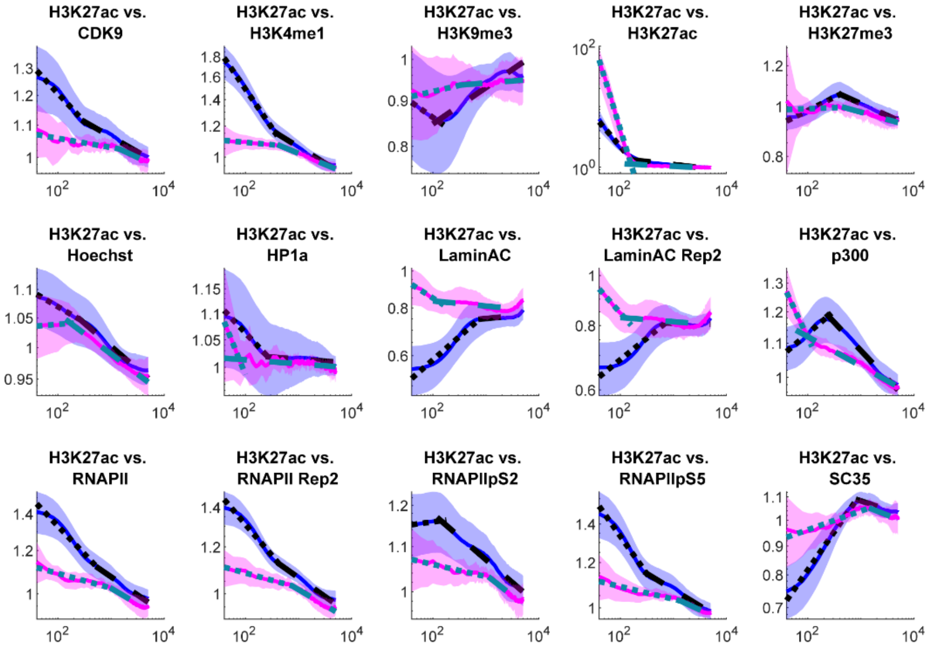

B

**Wilcoxon rank sum control vs. A485: H3K27ac vs.**

| | Hinge | Slope 1 | Slope 2 |
|---|---|---|---|
| **CDK9** | 0.831108 | 0.001737 | 0.027009 |
| **H3K4me1** | 0.06729 | 2.75E-08 | 0.004204 |
| **H3K9me3** | 0.761826 | 0.014855 | 0.18161 |
| **H3K27Ac** | 0.037831 | 2.75E-08 | 9.71E-08 |
| **H3K27me3** | 0.247596 | 0.141421 | 0.001875 |
| **Hoechst** | 0.266436 | 0.000442 | 0.727851 |
| **HP1a** | 0.129666 | 0.613461 | 0.507782 |
| **LaminAC** | 0.387395 | 3.19E-06 | 0.883977 |
| **LaminAC Rep2** | 0.551882 | 9.71E-08 | 0.098918 |
| **p300** | 0.000523 | 2.75E-08 | 5.58E-05 |
| **RNAPII** | 0.973136 | 0.003915 | 0.582291 |
| **RNAPII Rep2** | 0.937369 | 1.31E-06 | 0.160567 |
| **RNAPIIpS2** | 0.901728 | 0.340006 | 0.018971 |
| **RNAPIIpS5** | 0.522274 | 8.94E-05 | 0.007805 |
| **SC35** | 0.307015 | 2.02E-07 | 0.35149 |

**Figure S42. H3K27ac vs. all, A-485 vs. Control PCCF.**

(A) Normalized PCCF curves versus random (blue indicates control, magenta indicates A-485). (B) Wilcoxon rank sum p-values for each pair of comparisons.

A

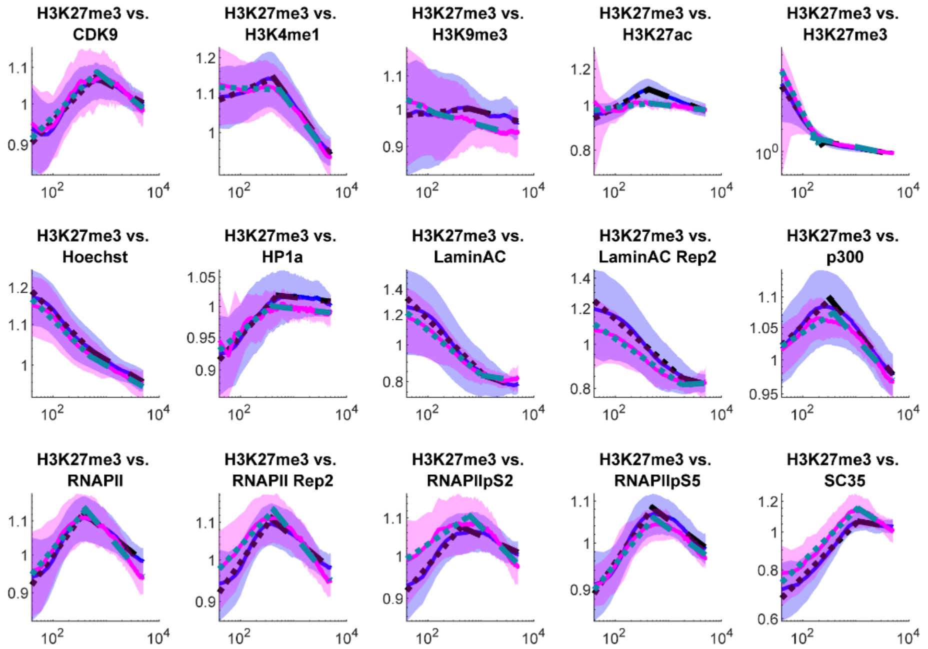

B

**Wilcoxon rank sum control vs. A485: H3K27me3 vs.**

| | Hinge | Slope 1 | Slope 2 |
|---|---|---|---|
| **CDK9** | 0.694403 | 0.711058 | 0.727851 |
| **H3K4me1** | 0.010152 | 0.001875 | 0.41253 |
| **H3K9me3** | 0.03203 | 0.04948 | 0.66154 |
| **H3K27Ac** | 0.001489 | 0.103594 | 4.91E-06 |
| **H3K27me3** | 0.016803 | 0.070719 | 0.129666 |
| **Hoechst** | 0.18161 | 0.566989 | 0.991044 |
| **HP1a** | 0.039953 | 0.883977 | 0.536975 |
| **LaminAC** | 0.919528 | 0.727851 | 0.629316 |
| **LaminAC Rep2** | 0.204633 | 0.266436 | 0.694403 |
| **p300** | 0.451993 | 0.296506 | 0.645345 |
| **RNAPII** | 0.479445 | 0.375186 | 0.17438 |
| **RNAPII Rep2** | 0.399844 | 0.28624 | 0.020145 |
| **RNAPIIpS2** | 0.229708 | 0.021381 | 0.013109 |
| **RNAPIIpS5** | 0.613461 | 0.046935 | 0.901728 |
| **SC35** | 0.761826 | 0.597784 | 0.001737 |

**Figure S43. H3K27me3 vs. all, A-485 vs. Control PCCF.**

(A) Normalized PCCF curves versus random (blue indicates control, magenta indicates A-485). (B) Wilcoxon rank sum p-values for each pair of comparisons.

A

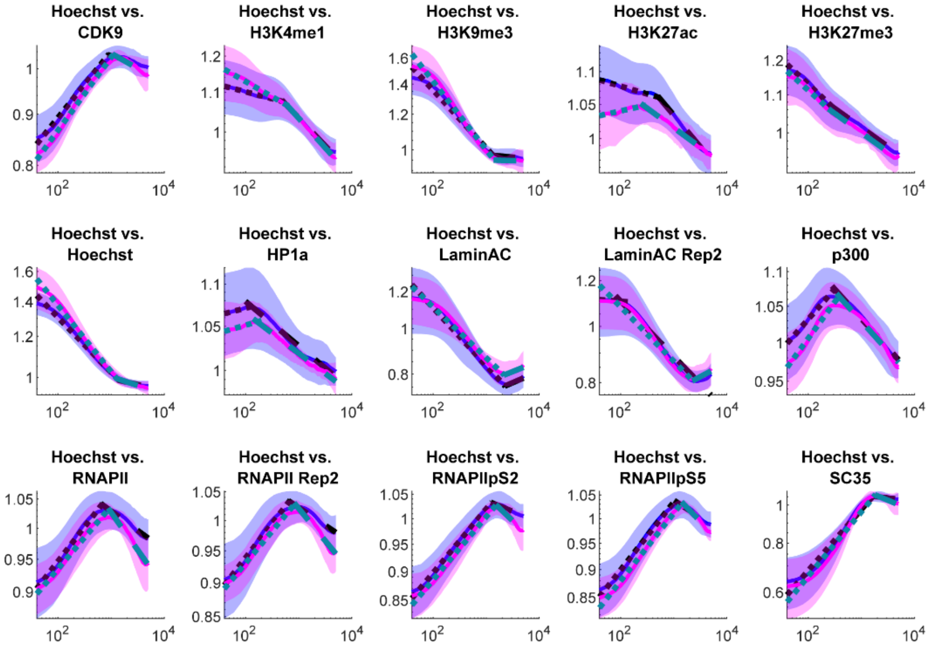

B

**Wilcoxon rank sum control vs. A485: Hoechst vs.**

| | Hinge | Slope 1 | Slope 2 |
|---|---|---|---|
| **CDK9** | 0.03203 | 0.135445 | 0.147598 |
| **H3K4me1** | 0.813641 | 0.012306 | 0.66154 |
| **H3K9me3** | 0.160567 | 0.027009 | 0.744776 |
| **H3K27Ac** | 0.070719 | 0.0073 | 0.00891 |
| **H3K27me3** | 0.937369 | 0.451993 | 0.645345 |
| **Hoechst** | 0.629316 | 0.04948 | 0.937369 |
| **HP1a** | 0.629316 | 0.613461 | 0.375186 |
| **LaminAC** | 0.796266 | 0.677895 | 0.694403 |
| **LaminAC Rep2** | 0.12408 | 0.831108 | 0.866284 |
| **p300** | 0.044502 | 0.052139 | 0.919528 |
| **RNAPII** | 0.001489 | 0.317768 | 0.000406 |
| **RNAPII Rep2** | 0.007805 | 0.613461 | 0.001005 |
| **RNAPIIpS2** | 0.004511 | 0.66154 | 0.00556 |
| **RNAPIIpS5** | 0.001005 | 0.937369 | 0.017858 |
| **SC35** | 0.037831 | 0.761826 | 0.008341 |

**Figure S44. Hoechst vs. all, A-485 vs. Control PCCF.**

(A) Normalized PCCF curves versus random (blue indicates control, magenta indicates A-485). (B) Wilcoxon rank sum p-values for each pair of comparisons.

A

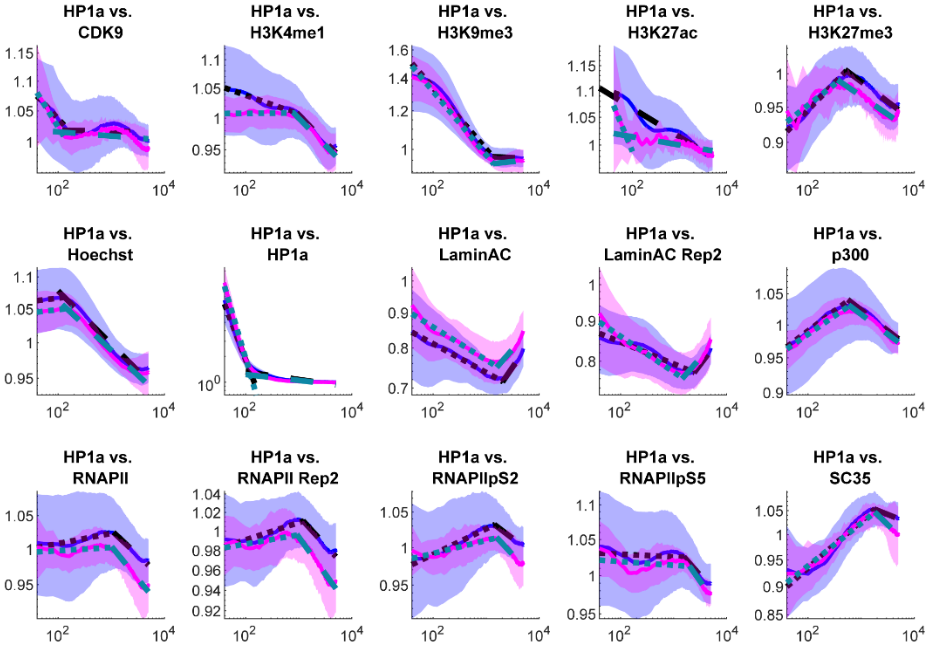

B

**Wilcoxon rank sum control vs. A485: HP1α vs.**

| | Hinge | Slope 1 | Slope 2 |
|---|---|---|---|
| CDK9 | 0.28624 | 0.848658 | 0.21276 |
| H3K4me1 | 0.694403 | 0.001005 | 0.973136 |
| H3K9me3 | 0.18906 | 0.167366 | 0.221118 |
| H3K27Ac | 0.006377 | 0.711058 | 0.17438 |
| H3K27me3 | 0.054917 | 0.778992 | 0.479445 |
| Hoechst | 0.507782 | 0.866284 | 0.991044 |
| HP1a | 0.238534 | 1.90E-05 | 0.060844 |
| LaminAC | 0.778992 | 0.551882 | 0.677895 |
| LaminAC Rep2 | 0.060844 | 0.060844 | 0.046935 |
| p300 | 0.677895 | 0.711058 | 0.991044 |
| RNAPII | 0.66154 | 0.465606 | 0.042175 |
| RNAPII Rep2 | 0.866284 | 0.831108 | 0.028601 |
| RNAPIIpS2 | 0.597784 | 0.013958 | 0.340006 |
| RNAPIIpS5 | 0.848658 | 1 | 0.41253 |
| SC35 | 0.399844 | 0.866284 | 0.103594 |

**Figure S45. HP1α vs. all, A-485 vs. Control PCCF.**

(A) Normalized PCCF curves versus random (blue indicates control, magenta indicates A-485). (B) Wilcoxon rank sum p-values for each pair of comparisons.

A

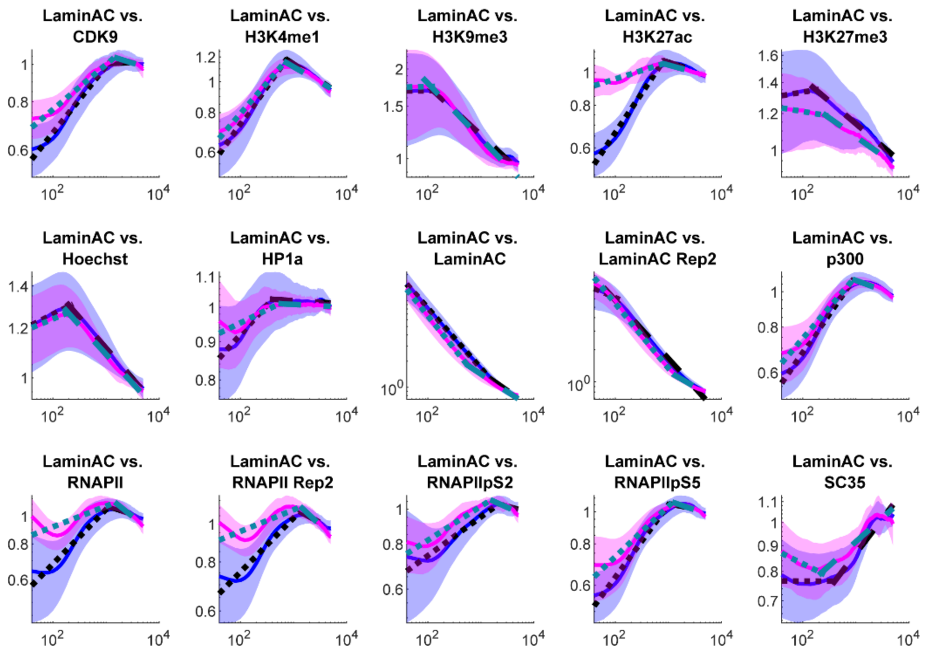

B

**Wilcoxon rank sum control vs. A485: Lamin A/C vs.**

| | Hinge | Slope 1 | Slope 2 |
|---|---|---|---|
| **CDK9** | 0.399844 | 0.006824 | 0.479445 |
| **H3K4me1** | 0.317768 | 0.070719 | 0.247596 |
| **H3K9me3** | 0.937369 | 0.973136 | 0.507782 |
| **H3K27Ac** | 0.229708 | 8.31E-07 | 0.296506 |
| **H3K27me3** | 0.090071 | 0.387395 | 1 |
| **Hoechst** | 0.28624 | 0.438607 | 0.238534 |
| **HP1a** | 0.129666 | 0.008341 | 0.451993 |
| **LaminAC** | 0.399844 | 0.141421 | 0.052139 |
| **LaminAC Rep2** | 0.060844 | 0.006824 | 0.002725 |
| **p300** | 0.613461 | 0.06729 | 0.522274 |
| **RNAPII** | 0.003915 | 0.000118 | 0.003915 |
| **RNAPII Rep2** | 0.28624 | 0.00235 | 0.002933 |
| **RNAPIIpS2** | 0.375186 | 0.582291 | 0.007805 |
| **RNAPIIpS5** | 0.160567 | 0.013958 | 0.074289 |
| **SC35** | 0.017858 | 0.018971 | 0.022682 |

**Figure S46. Lamin A/C vs. all, A-485 vs. Control PCCF.**

(A) Normalized PCCF curves versus random (blue indicates control, magenta indicates A-485). (B) Wilcoxon rank sum p-values for each pair of comparisons.

A

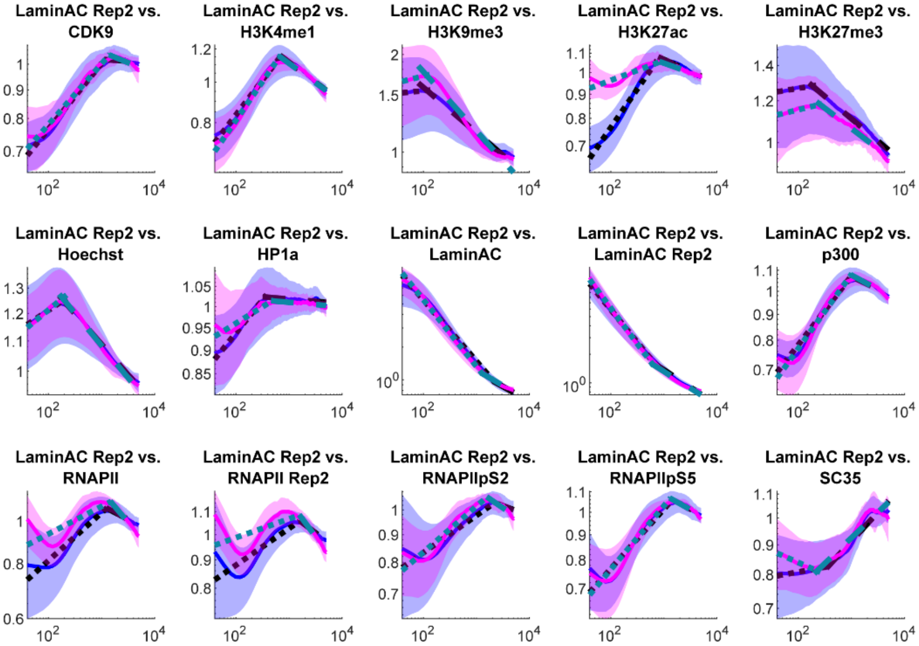

B

**Wilcoxon rank sum control vs. A485: Lamin A/C repeat vs.**

| | Hinge | Slope 1 | Slope 2 |
|---|---|---|---|
| **CDK9** | 0.266436 | 0.761826 | 0.796266 |
| **H3K4me1** | 0.153979 | 0.35149 | 0.507782 |
| **H3K9me3** | 0.129666 | 0.256896 | 0.613461 |
| **H3K27Ac** | 0.266436 | 9.32E-07 | 0.103594 |
| **H3K27me3** | 0.645345 | 0.135445 | 0.813641 |
| **Hoechst** | 0.247596 | 0.645345 | 0.582291 |
| **HP1a** | 0.35149 | 0.035805 | 0.613461 |
| **LaminAC** | 0.167366 | 0.085892 | 0.028601 |
| **LaminAC Rep2** | 0.387395 | 0.204633 | 0.796266 |
| **p300** | 0.613461 | 0.493505 | 0.399844 |
| **RNAPII** | 0.317768 | 0.044502 | 0.022682 |
| **RNAPII Rep2** | 0.044502 | 0.266436 | 0.005188 |
| **RNAPIIpS2** | 0.118684 | 0.21276 | 0.00235 |
| **RNAPIIpS5** | 0.866284 | 0.522274 | 0.465606 |
| **SC35** | 0.039953 | 0.001875 | 0.266436 |

**Figure S47. Lamin A/C repeat vs. all, A-485 vs. Control PCCF.**

(A) Normalized PCCF curves versus random (blue indicates control, magenta indicates A-485). (B) Wilcoxon rank sum p-values for each pair of comparisons.

A

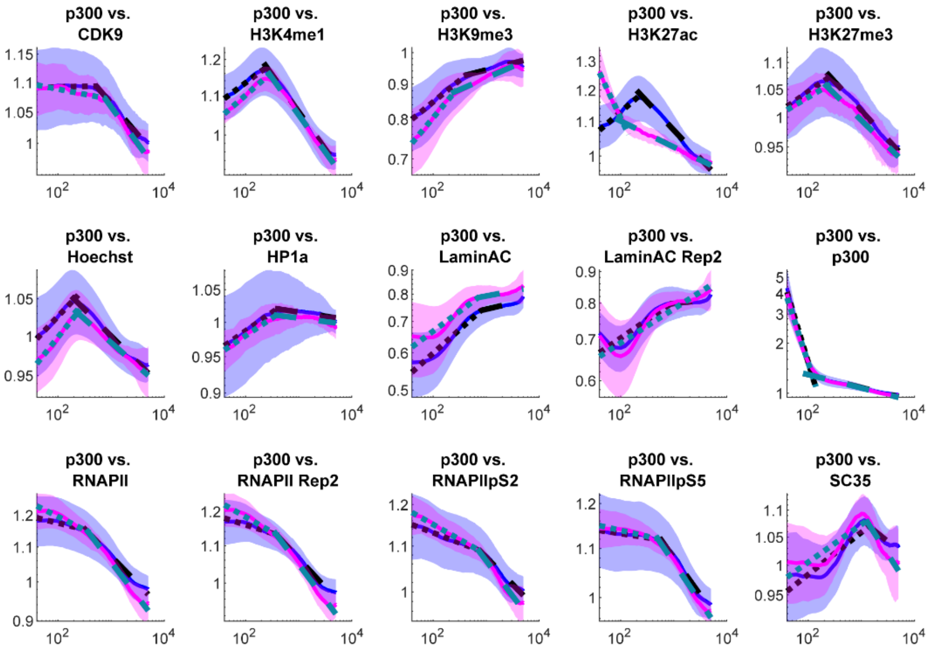

B

**Wilcoxon rank sum control vs. A485: p300 vs.**

| | Hinge | Slope 1 | Slope 2 |
|---|---|---|---|
| CDK9 | 0.135445 | 0.06729 | 0.991044 |
| H3K4me1 | 0.013958 | 0.507782 | 0.761826 |
| H3K9me3 | 0.919528 | 0.037831 | 0.937369 |
| H3K27Ac | 0.000928 | 2.75E-08 | 5.88E-07 |
| H3K27me3 | 0.711058 | 0.522274 | 0.761826 |
| Hoechst | 0.074289 | 0.113474 | 0.438607 |
| HP1a | 0.711058 | 0.991044 | 0.266436 |
| LaminAC | 0.147598 | 0.296506 | 0.328765 |
| LaminAC Rep2 | 0.438607 | 0.778992 | 0.694403 |
| p300 | 0.645345 | 0.387395 | 0.256896 |
| RNAPII | 1 | 0.01083 | 0.037831 |
| RNAPII Rep2 | 0.363217 | 0.008341 | 0.037831 |
| RNAPIIpS2 | 0.848658 | 0.030274 | 0.078006 |
| RNAPIIpS5 | 0.778992 | 0.328765 | 0.221118 |
| SC35 | 0.901728 | 0.582291 | 0.013109 |

**Figure S48. p300 vs. all, A-485 vs. Control PCCF.**

(A) Normalized PCCF curves versus random (blue indicates control, magenta indicates A-485). (B) Wilcoxon rank sum p-values for each pair of comparisons.

A

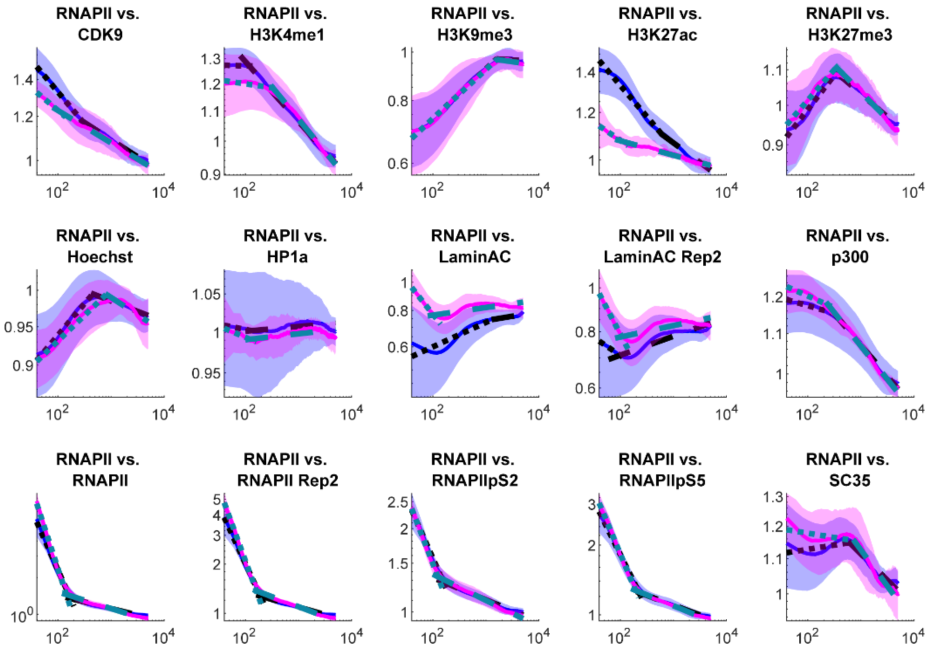

B

**Wilcoxon rank sum control vs. A485: RNAPII vs.**

| | Hinge | Slope 1 | Slope 2 |
|---|---|---|---|
| **CDK9** | 0.566989 | 0.016803 | 0.425452 |
| **H3K4me1** | 0.000288 | 0.744776 | 0.919528 |
| **H3K9me3** | 0.566989 | 0.813641 | 0.196734 |
| **H3K27Ac** | 0.001378 | 0.015802 | 0.276217 |
| **H3K27me3** | 0.761826 | 0.566989 | 0.551882 |
| **Hoechst** | 0.03203 | 0.17438 | 0.153979 |
| **HP1a** | 0.866284 | 0.848658 | 0.465606 |
| **LaminAC** | 0.03203 | 6.75E-05 | 0.21276 |
| **LaminAC Rep2** | 0.153979 | 0.000108 | 0.727851 |
| **p300** | 0.147598 | 0.160567 | 0.141421 |
| **RNAPII** | 0.000185 | 0.000118 | 0.001274 |
| **RNAPII Rep2** | 4.61E-05 | 0.000481 | 8.94E-05 |
| **RNAPIIpS2** | 5.23E-07 | 0.046935 | 0.060844 |
| **RNAPIIpS5** | 0.000185 | 0.003915 | 0.017858 |
| **SC35** | 0.129666 | 0.003392 | 0.221118 |

**Figure S49. RNAPII vs. all, A-485 vs. Control PCCF.**

(A) Normalized PCCF curves versus random (blue indicates control, magenta indicates A-485). (B) Wilcoxon rank sum p-values for each pair of comparisons.

A

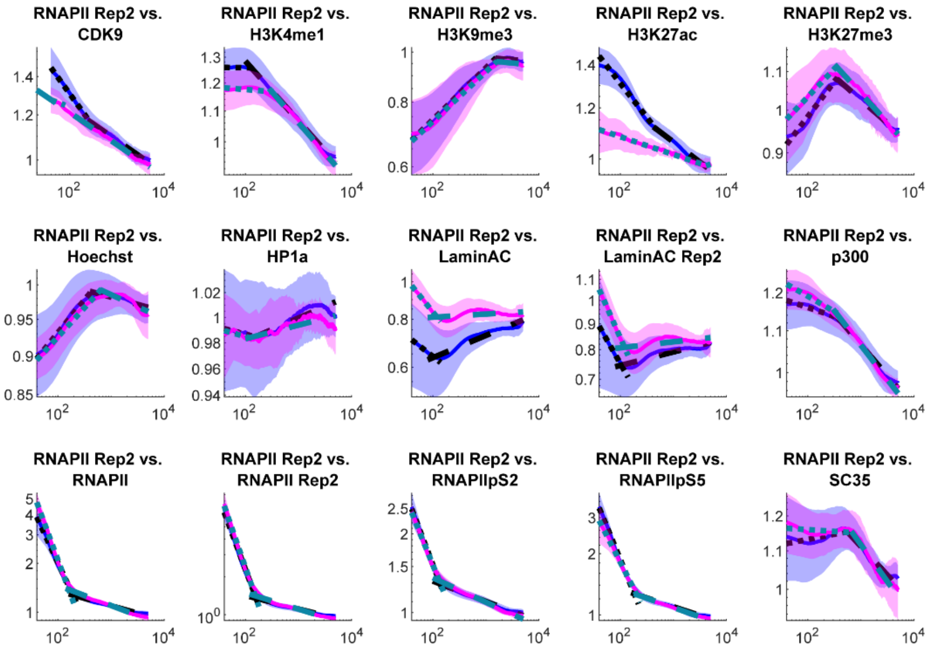

B

**Wilcoxon rank sum control vs. A485: RNAPII repeat vs.**

| | Hinge | Slope 1 | Slope 2 |
|---|---|---|---|
| CDK9 | 0.901728 | 0.000203 | 0.727851 |
| H3K4me1 | 0.000481 | 0.761826 | 0.761826 |
| H3K9me3 | 0.901728 | 0.18161 | 0.551882 |
| H3K27Ac | 0.465606 | 8.31E-07 | 0.020145 |
| H3K27me3 | 0.919528 | 0.41253 | 0.12408 |
| Hoechst | 0.037831 | 0.479445 | 0.118684 |
| HP1a | 0.582291 | 0.831108 | 0.536975 |
| LaminAC | 0.027009 | 0.001737 | 0.12408 |
| LaminAC Rep2 | 0.141421 | 0.03203 | 0.883977 |
| p300 | 0.629316 | 0.020145 | 0.21276 |
| RNAPII | 6.14E-05 | 0.000523 | 0.00017 |
| RNAPII Rep2 | 0.465606 | 0.35149 | 0.003915 |
| RNAPIIpS2 | 0.399844 | 0.108445 | 0.03203 |
| RNAPIIpS5 | 0.027009 | 0.012306 | 0.507782 |
| SC35 | 0.645345 | 0.135445 | 0.307015 |

**Figure S50. RNAPII repeat vs. all, A-485 vs. Control PCCF.**

(A) Normalized PCCF curves versus random (blue indicates control, magenta indicates A-485). (B) Wilcoxon rank sum p-values for each pair of comparisons.

A

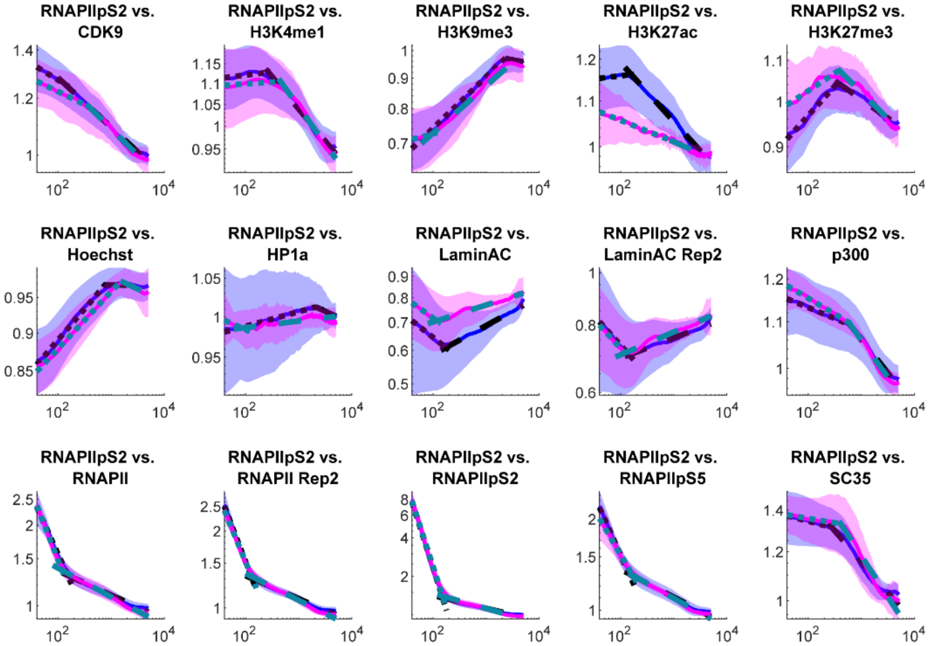

B

**Wilcoxon rank sum control vs. A485: RNAPII pS2 vs.**

| | Hinge | Slope 1 | Slope 2 |
|---|---|---|---|
| **CDK9** | 0.522274 | 0.042175 | 0.831108 |
| **H3K4me1** | 0.006824 | 0.425452 | 0.866284 |
| **H3K9me3** | 0.098918 | 0.013958 | 0.41253 |
| **H3K27Ac** | 0.796266 | 0.493505 | 3.96E-06 |
| **H3K27me3** | 0.866284 | 0.039953 | 0.276217 |
| **Hoechst** | 0.221118 | 0.167366 | 0.937369 |
| **HP1a** | 0.451993 | 0.18161 | 0.711058 |
| **LaminAC** | 0.017858 | 0.566989 | 0.01083 |
| **LaminAC Rep2** | 0.375186 | 0.991044 | 0.387395 |
| **p300** | 0.479445 | 0.007805 | 0.153979 |
| **RNAPII** | 4.65E-07 | 0.027009 | 0.078006 |
| **RNAPII Rep2** | 0.17438 | 0.153979 | 0.011547 |
| **RNAPIIpS2** | 0.677895 | 0.536975 | 0.108445 |
| **RNAPIIpS5** | 0.00218 | 0.001875 | 0.438607 |
| **SC35** | 0.276217 | 0.66154 | 0.229708 |

**Figure S51. RNAPII pS2 vs. all, A-485 vs. Control PCCF.**

(A) Normalized PCCF curves versus random (blue indicates control, magenta indicates A-485). (B) Wilcoxon rank sum p-values for each pair of comparisons.

A

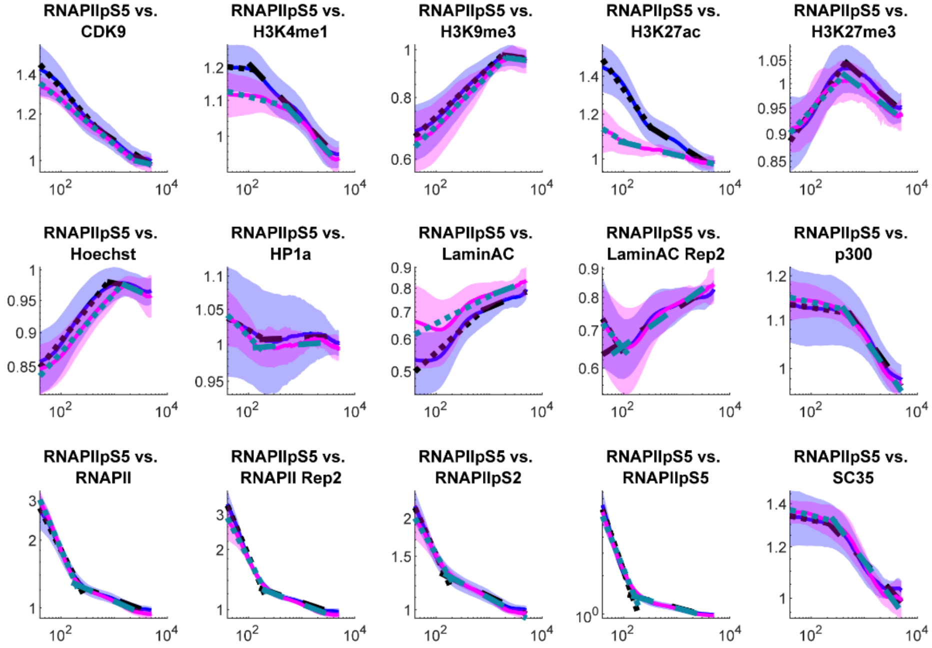

B

**Wilcoxon rank sum control vs. A485: RNAPII pS5 vs.**

| | Hinge | Slope 1 | Slope 2 |
|---|---|---|---|
| **CDK9** | 0.317768 | 0.004204 | 0.901728 |
| **H3K4me1** | 0.238534 | 0.238534 | 0.677895 |
| **H3K9me3** | 0.866284 | 0.677895 | 0.883977 |
| **H3K27Ac** | 0.017858 | 0.000242 | 6.75E-06 |
| **H3K27me3** | 0.387395 | 0.064 | 0.465606 |
| **Hoechst** | 0.016803 | 0.221118 | 0.256896 |
| **HP1a** | 0.479445 | 0.153979 | 0.813641 |
| **LaminAC** | 0.039953 | 0.00218 | 0.17438 |
| **LaminAC Rep2** | 0.937369 | 0.744776 | 0.17438 |
| **p300** | 0.196734 | 0.41253 | 0.276217 |
| **RNAPII** | 0.000222 | 0.003392 | 0.003155 |
| **RNAPII Rep2** | 0.037831 | 0.014855 | 0.153979 |
| **RNAPIIpS2** | 0.002933 | 0.001737 | 0.276217 |
| **RNAPIIpS5** | 0.0073 | 0.035805 | 0.613461 |
| **SC35** | 0.991044 | 0.160567 | 0.118684 |

**Figure S52. RNAPII pS5 vs. all, A-485 vs. Control PCCF.**

(A) Normalized PCCF curves versus random (blue indicates control, magenta indicates A-485). (B) Wilcoxon rank sum p-values for each pair of comparisons.

A

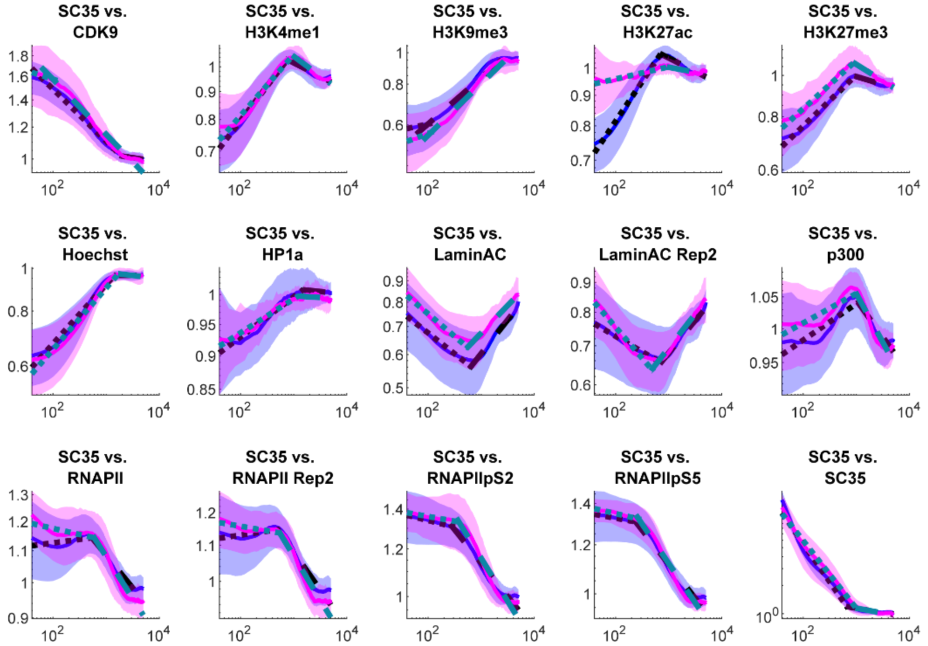

B

**Wilcoxon rank sum control vs. A485: SC-35 vs.**

| | Hinge | Slope 1 | Slope 2 |
|---|---|---|---|
| **CDK9** | 0.645345 | 0.078006 | 0.493505 |
| **H3K4me1** | 0.003392 | 0.141421 | 0.340006 |
| **H3K9me3** | 0.901728 | 0.901728 | 0.465606 |
| **H3K27Ac** | 0.35149 | 2.86E-06 | 5.58E-05 |
| **H3K27me3** | 0.597784 | 0.328765 | 0.005188 |
| **Hoechst** | 0.796266 | 0.317768 | 0.711058 |
| **HP1a** | 0.018971 | 0.677895 | 0.582291 |
| **LaminAC** | 0.375186 | 0.973136 | 0.090071 |
| **LaminAC Rep2** | 0.008341 | 0.000671 | 0.438607 |
| **p300** | 0.582291 | 0.098918 | 0.025494 |
| **RNAPII** | 0.425452 | 0.003645 | 0.522274 |
| **RNAPII Rep2** | 0.438607 | 0.035805 | 0.41253 |
| **RNAPIIpS2** | 0.35149 | 0.582291 | 0.375186 |
| **RNAPIIpS5** | 0.955241 | 0.18906 | 0.196734 |
| **SC35** | 0.399844 | 0.090071 | 0.813641 |

**Figure S53. SC-35 vs. all, A-485 vs. Control PCCF.**

(A) Normalized PCCF curves versus random (blue indicates control, magenta indicates A-485). (B) Wilcoxon rank sum p-values for each pair of comparisons.

A

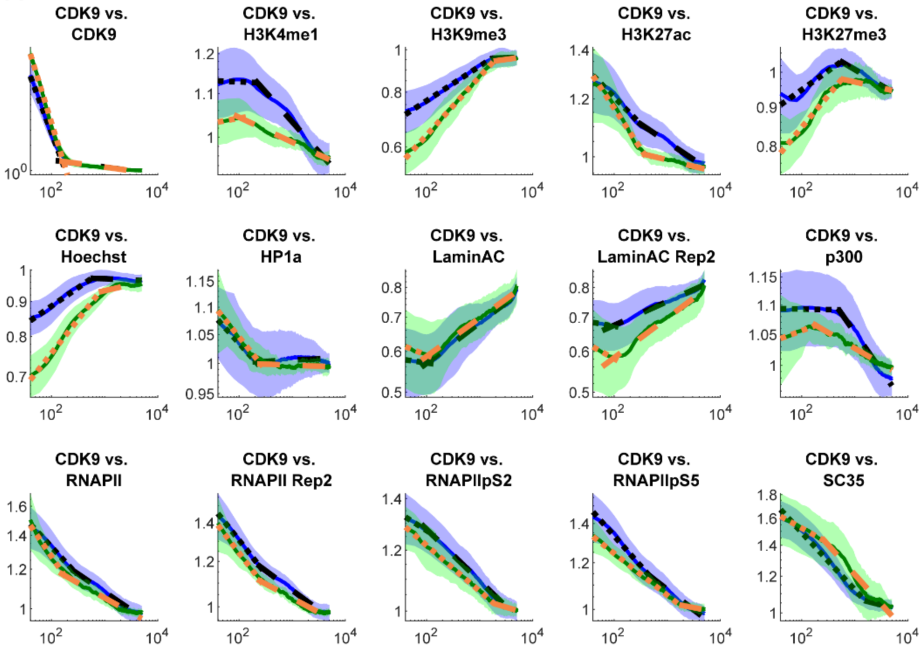

B

**Wilcoxon rank sum control vs. αAM: CDK9 vs.**

| | Hinge | Slope 1 | Slope 2 |
|---|---|---|---|
| **CDK9** | 0.129666 | 7.41E-07 | 0.677895 |
| **H3K4me1** | 0.113474 | 0.307015 | 3.12E-05 |
| **H3K9me3** | 0.438607 | 0.006824 | 0.129666 |
| **H3K27Ac** | 0.317768 | 0.005188 | 1.90E-05 |
| **H3K27me3** | 0.796266 | 0.006824 | 0.004204 |
| **Hoechst** | 0.147598 | 1.40E-07 | 0.000569 |
| **HP1a** | 0.813641 | 0.597784 | 0.35149 |
| **LaminAC** | 0.465606 | 0.307015 | 0.28624 |
| **LaminAC Rep2** | 0.387395 | 0.479445 | 0.052139 |
| **p300** | 0.01083 | 0.090071 | 2.32E-05 |
| **RNAPII** | 0.046935 | 0.081872 | 0.135445 |
| **RNAPII Rep2** | 0.629316 | 0.796266 | 0.060844 |
| **RNAPIIpS2** | 0.813641 | 0.66154 | 0.06729 |
| **RNAPIIpS5** | 0.17438 | 0.247596 | 0.153979 |
| **SC35** | 0.387395 | 0.001737 | 0.070719 |

**Figure S54. CDK9 vs. all, αAM vs. Control PCCF.**

(A) Normalized PCCF curves versus random (blue indicates control, green indicates αAM). (B) Wilcoxon rank sum p-values for each pair of comparisons.

A

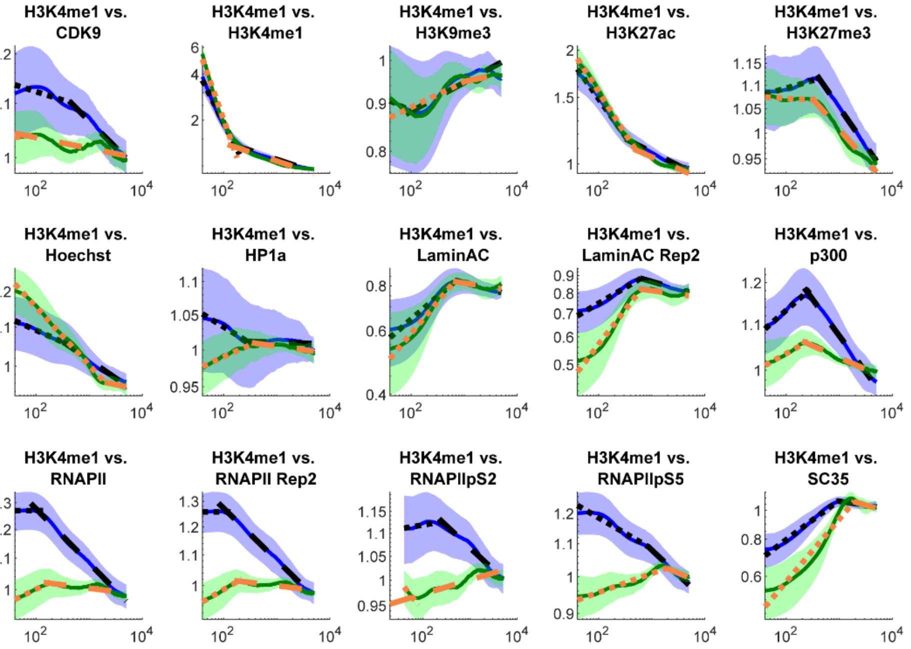

B

**Wilcoxon rank sum control vs. αAM: H3K4me1 vs.**

| | Hinge | Slope 1 | Slope 2 |
|---|---|---|---|
| CDK9 | 0.12408 | 0.744776 | 2.56E-05 |
| H3K4me1 | 0.103594 | 1.26E-05 | 0.66154 |
| H3K9me3 | 0.522274 | 0.778992 | 0.901728 |
| H3K27Ac | 0.042175 | 0.094411 | 0.003645 |
| H3K27me3 | 0.006824 | 0.014855 | 0.094411 |
| Hoechst | 0.328765 | 1.84E-06 | 0.566989 |
| HP1a | 0.35149 | 6.07E-06 | 0.247596 |
| LaminAC | 0.831108 | 0.028601 | 0.551882 |
| LaminAC Rep2 | 0.955241 | 4.41E-06 | 0.12408 |
| p300 | 0.340006 | 0.078006 | 2.02E-07 |
| RNAPII | 0.021381 | 2.05E-06 | 7.42E-05 |
| RNAPII Rep2 | 0.094411 | 6.60E-07 | 3.19E-06 |
| RNAPIIpS2 | 0.901728 | 0.493505 | 2.10E-05 |
| RNAPIIpS5 | 0.340006 | 3.55E-06 | 0.000314 |
| SC35 | 0.866284 | 0.919528 | 0.35149 |

**Figure S55. H3K4me1 vs. all, αAM vs. Control PCCF.**

(A) Normalized PCCF curves versus random (blue indicates control, green indicates αAM). (B) Wilcoxon rank sum p-values for each pair of comparisons.

A

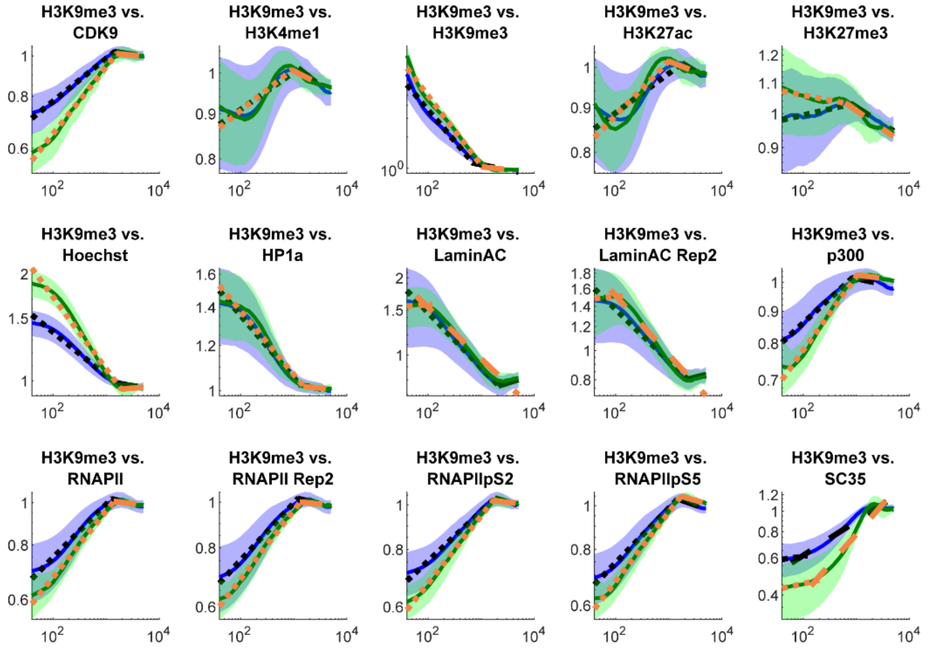

B

**Wilcoxon rank sum control vs. αAM: H3K9me3 vs.**

| | Hinge | Slope 1 | Slope 2 |
|---|---|---|---|
| CDK9 | 1 | 0.001737 | 0.28624 |
| H3K4me1 | 0.328765 | 0.238534 | 0.778992 |
| H3K9me3 | 0.035805 | 0.113474 | 0.387395 |
| H3K27Ac | 0.256896 | 0.070719 | 0.937369 |
| H3K27me3 | 0.582291 | 0.042175 | 0.991044 |
| Hoechst | 0.317768 | 0.002725 | 0.778992 |
| HP1a | 0.66154 | 0.522274 | 0.425452 |
| LaminAC | 0.551882 | 0.135445 | 0.238534 |
| LaminAC Rep2 | 0.016803 | 0.078006 | 0.078006 |
| p300 | 0.044502 | 0.000342 | 0.340006 |
| RNAPII | 0.399844 | 0.008341 | 0.057817 |
| RNAPII Rep2 | 0.597784 | 0.003915 | 0.090071 |
| RNAPIIpS2 | 0.66154 | 0.001274 | 0.438607 |
| RNAPIIpS5 | 0.03203 | 0.002022 | 0.761826 |
| SC35 | 0.848658 | 0.113474 | 1.71E-05 |

**Figure S56. H3K9me3 vs. all, αAM vs. Control PCCF.**

(A) Normalized PCCF curves versus random (blue indicates control, green indicates αAM). (B) Wilcoxon rank sum p-values for each pair of comparisons.

A

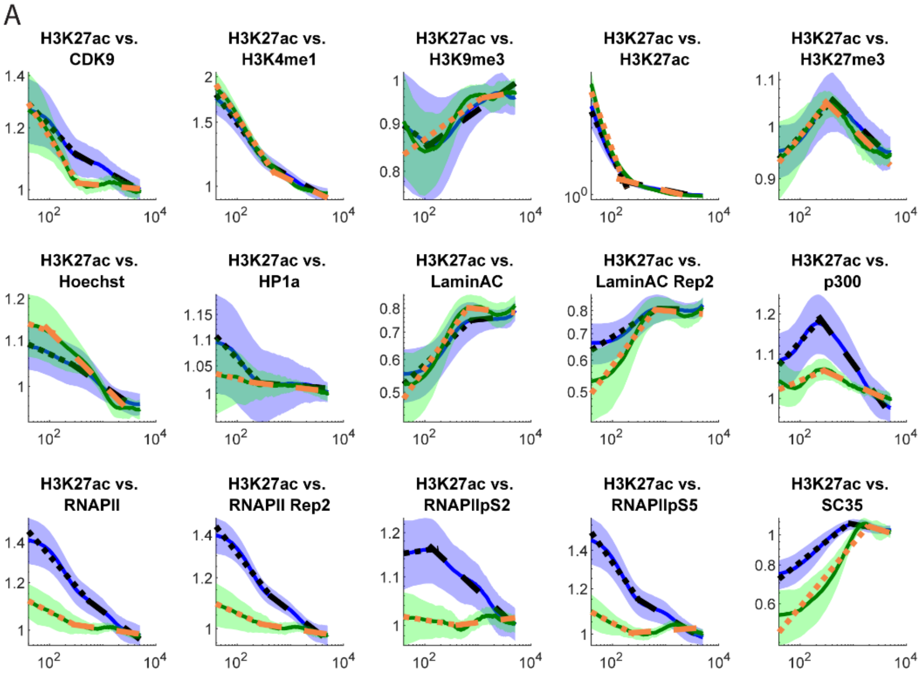

B

Wilcoxon rank sum control vs. αAM: H3K27ac vs.

| | Hinge | Slope 1 | Slope 2 |
|---|---|---|---|
| CDK9 | 0.955241 | 0.015802 | 2.05E-06 |
| H3K4me1 | 0.039953 | 0.118684 | 0.001005 |
| H3K9me3 | 0.831108 | 0.613461 | 0.153979 |
| H3K27Ac | 0.551882 | 0.00218 | 0.015802 |
| H3K27me3 | 0.000155 | 0.387395 | 0.955241 |
| Hoechst | 0.147598 | 0.848658 | 0.340006 |
| HP1a | 0.021381 | 0.000222 | 0.135445 |
| LaminAC | 0.317768 | 0.014855 | 0.090071 |
| LaminAC Rep2 | 0.438607 | 1.90E-05 | 0.340006 |
| p300 | 0.167366 | 0.001378 | 9.71E-08 |
| RNAPII | 0.070719 | 3.12E-05 | 0.000222 |
| RNAPII Rep2 | 0.566989 | 1.47E-06 | 8.15E-05 |
| RNAPIIpS2 | 0.66154 | 0.831108 | 1.03E-05 |
| RNAPIIpS5 | 0.033872 | 3.19E-06 | 1.04E-06 |
| SC35 | 0.831108 | 0.796266 | 0.204633 |

**Figure S57. H3K27ac vs. all, αAM vs. Control PCCF.**

(A) Normalized PCCF curves versus random (blue indicates control, green indicates αAM). (B) Wilcoxon rank sum p-values for each pair of comparisons.

A

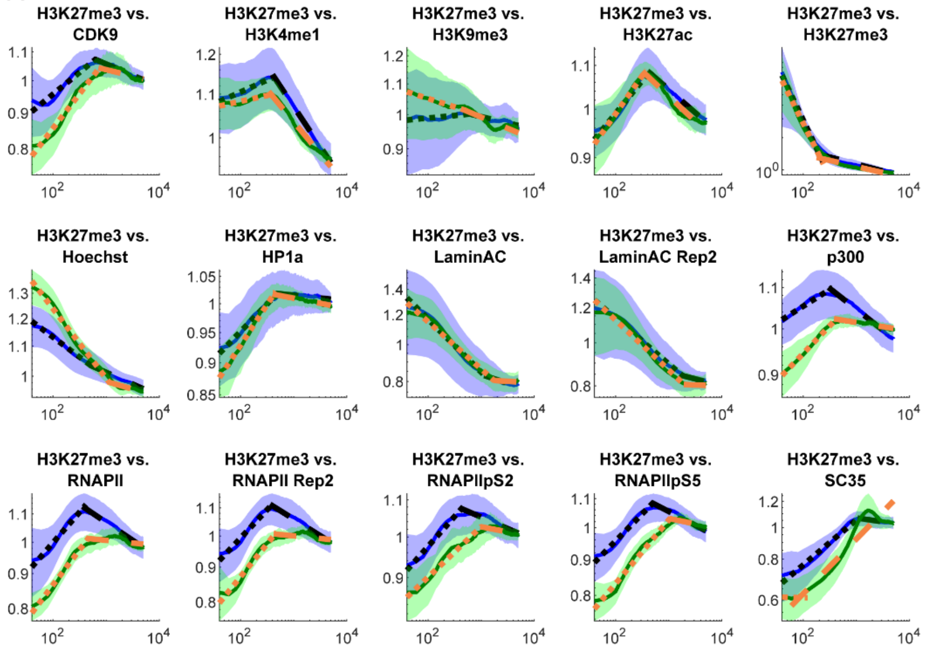

B

**Wilcoxon rank sum control vs. αAM: H3K27me3 vs.**

| | Hinge | Slope 1 | Slope 2 |
|---|---|---|---|
| **CDK9** | 0.113474 | 0.001609 | 0.060844 |
| **H3K4me1** | 0.081872 | 0.070719 | 0.081872 |
| **H3K9me3** | 0.35149 | 0.256896 | 0.451993 |
| **H3K27Ac** | 0.000406 | 0.28624 | 0.901728 |
| **H3K27me3** | 0.831108 | 0.597784 | 0.582291 |
| **Hoechst** | 0.629316 | 4.58E-08 | 0.613461 |
| **HP1a** | 0.129666 | 0.001609 | 0.296506 |
| **LaminAC** | 0.296506 | 0.551882 | 0.711058 |
| **LaminAC Rep2** | 0.831108 | 0.645345 | 0.328765 |
| **p300** | 0.085892 | 0.001378 | 1.84E-06 |
| **RNAPII** | 0.070719 | 0.493505 | 0.000373 |
| **RNAPII Rep2** | 0.848658 | 0.070719 | 1.71E-05 |
| **RNAPIIpS2** | 0.085892 | 0.901728 | 0.005956 |
| **RNAPIIpS5** | 0.003392 | 0.001274 | 0.003392 |
| **SC35** | 0.727851 | 0.118684 | 0.090071 |

**Figure S58. H3K27me3 vs. all, αAM vs. Control PCCF.**

(A) Normalized PCCF curves versus random (blue indicates control, green indicates αAM). (B) Wilcoxon rank sum p-values for each pair of comparisons.

A

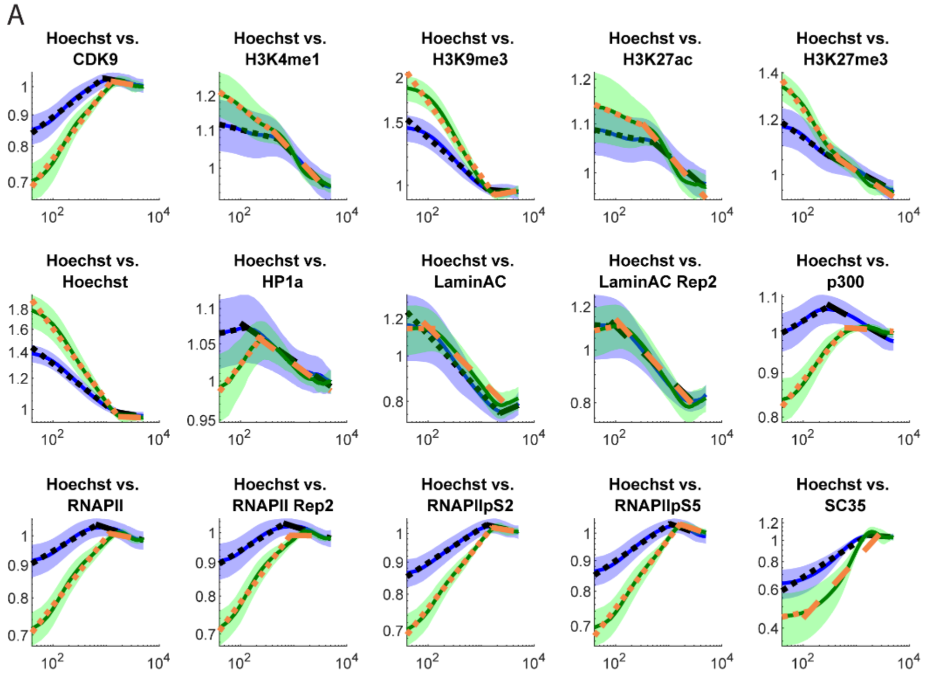

B

**Wilcoxon rank sum control vs. αAM: Hoechst vs.**

| | Hinge | Slope 1 | Slope 2 |
|---|---|---|---|
| CDK9 | 0.006377 | 1.10E-07 | 0.507782 |
| H3K4me1 | 0.21276 | 6.07E-06 | 0.744776 |
| H3K9me3 | 0.013958 | 4.18E-05 | 0.438607 |
| H3K27Ac | 0.03203 | 0.387395 | 0.118684 |
| H3K27me3 | 0.094411 | 2.75E-08 | 0.677895 |
| Hoechst | 0.003392 | 2.75E-08 | 0.01083 |
| HP1a | 0.711058 | 1.17E-06 | 0.711058 |
| LaminAC | 0.831108 | 0.090071 | 0.744776 |
| LaminAC Rep2 | 0.021381 | 0.002725 | 0.013958 |
| p300 | 1.84E-06 | 9.71E-08 | 1.17E-06 |
| RNAPII | 0.005956 | 2.75E-08 | 0.025494 |
| RNAPII Rep2 | 0.167366 | 3.13E-08 | 0.021381 |
| RNAPIIpS2 | 0.00556 | 7.58E-08 | 0.493505 |
| RNAPIIpS5 | 0.000928 | 5.90E-08 | 0.813641 |
| SC35 | 0.000671 | 0.001178 | 6.07E-06 |

**Figure S59. Hoechst vs. all, αAM vs. Control PCCF.**

(A) Normalized PCCF curves versus random (blue indicates control, green indicates αAM). (B) Wilcoxon rank sum p-values for each pair of comparisons.

A

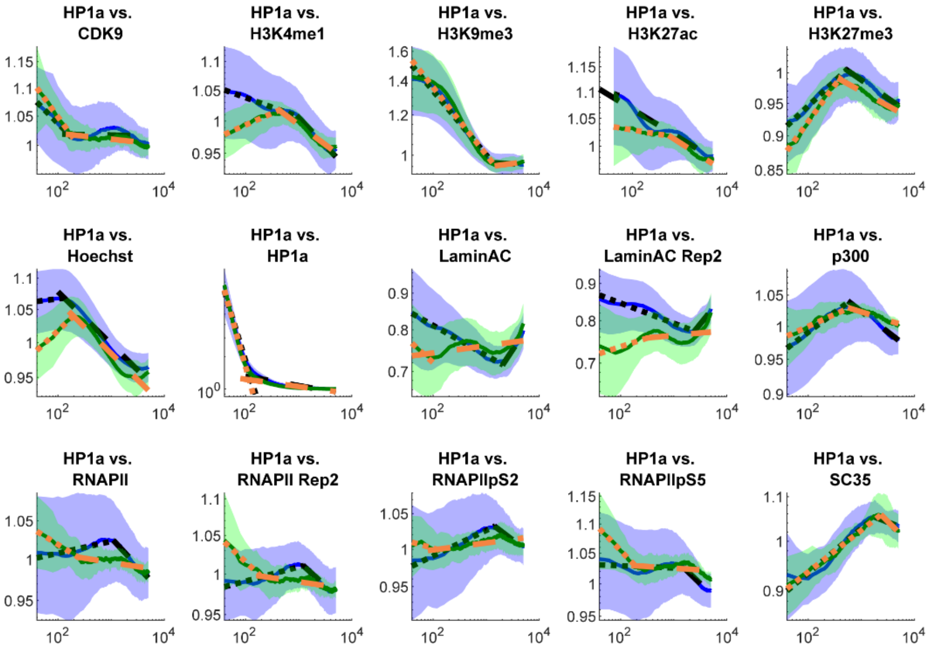

B

**Wilcoxon rank sum control vs. αAM: HP1α vs.**

| | Hinge | Slope 1 | Slope 2 |
|---|---|---|---|
| **CDK9** | 0.340006 | 0.147598 | 0.41253 |
| **H3K4me1** | 0.221118 | 1.31E-06 | 0.147598 |
| **H3K9me3** | 0.12408 | 0.778992 | 0.375186 |
| **H3K27Ac** | 0.266436 | 0.000222 | 0.375186 |
| **H3K27me3** | 0.000142 | 0.0073 | 0.153979 |
| **Hoechst** | 0.317768 | 2.28E-07 | 0.17438 |
| **HP1a** | 0.018971 | 0.004839 | 0.064 |
| **LaminAC** | 0.042175 | 0.06729 | 0.081872 |
| **LaminAC Rep2** | 0.425452 | 0.21276 | 0.919528 |
| **p300** | 0.266436 | 0.113474 | 0.000618 |
| **RNAPII** | 0.002725 | 0.000728 | 0.04948 |
| **RNAPII Rep2** | 0.00556 | 0.000203 | 0.003155 |
| **RNAPIIpS2** | 0.001489 | 0.017858 | 0.06729 |
| **RNAPIIpS5** | 0.022682 | 0.025494 | 0.118684 |
| **SC35** | 0.41253 | 0.813641 | 0.256896 |

**Figure S60. HP1α vs. all, αAM vs. Control PCCF.**

(A) Normalized PCCF curves versus random (blue indicates control, green indicates αAM). (B) Wilcoxon rank sum p-values for each pair of comparisons.

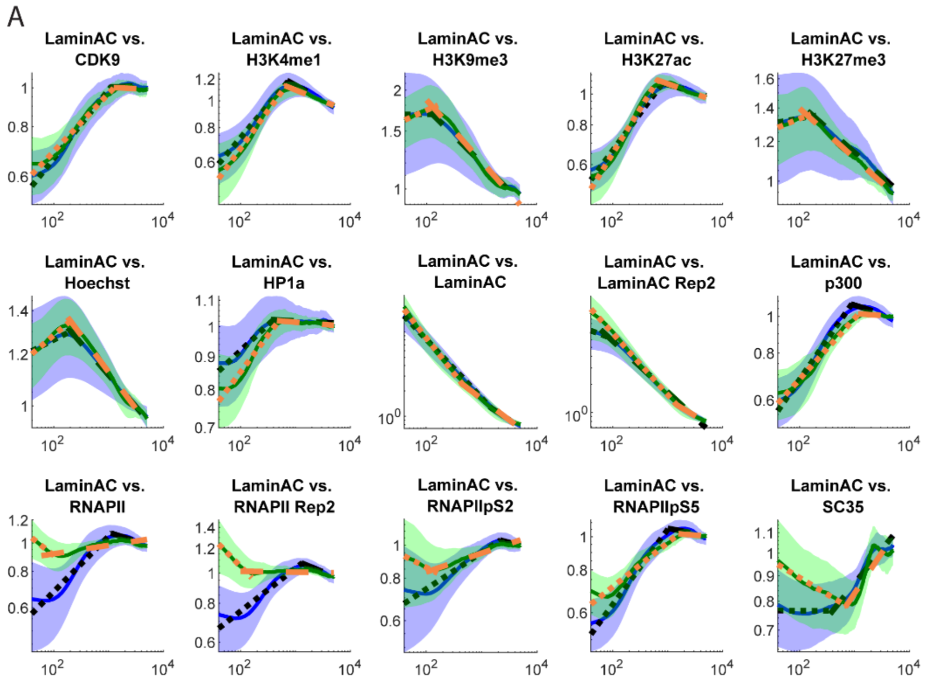

B

**Wilcoxon rank sum control vs. αAM: Lamin A/C vs.**

| | Hinge | Slope 1 | Slope 2 |
|---|---|---|---|
| **CDK9** | 0.387395 | 0.35149 | 0.937369 |
| **H3K4me1** | 0.06729 | 0.06729 | 0.054917 |
| **H3K9me3** | 0.03203 | 0.074289 | 0.296506 |
| **H3K27Ac** | 0.00891 | 0.081872 | 0.645345 |
| **H3K27me3** | 0.011547 | 0.052139 | 0.613461 |
| **Hoechst** | 0.901728 | 0.147598 | 0.778992 |
| **HP1a** | 0.35149 | 0.016803 | 0.582291 |
| **LaminAC** | 0.831108 | 0.015802 | 0.493505 |
| **LaminAC Rep2** | 0.238534 | 0.005188 | 0.221118 |
| **p300** | 0.039953 | 0.006377 | 0.001737 |
| **RNAPII** | 0.135445 | 1.59E-07 | 0.03203 |
| **RNAPII Rep2** | 0.001489 | 4.14E-07 | 0.002933 |
| **RNAPIIpS2** | 0.551882 | 0.013109 | 0.813641 |
| **RNAPIIpS5** | 0.493505 | 3.12E-05 | 0.025494 |
| **SC35** | 0.00891 | 0.001609 | 0.238534 |

**Figure S61. Lamin A/C vs. all, αAM vs. Control PCCF.**

(A) Normalized PCCF curves versus random (blue indicates control, green indicates αAM). (B) Wilcoxon rank sum p-values for each pair of comparisons.

A

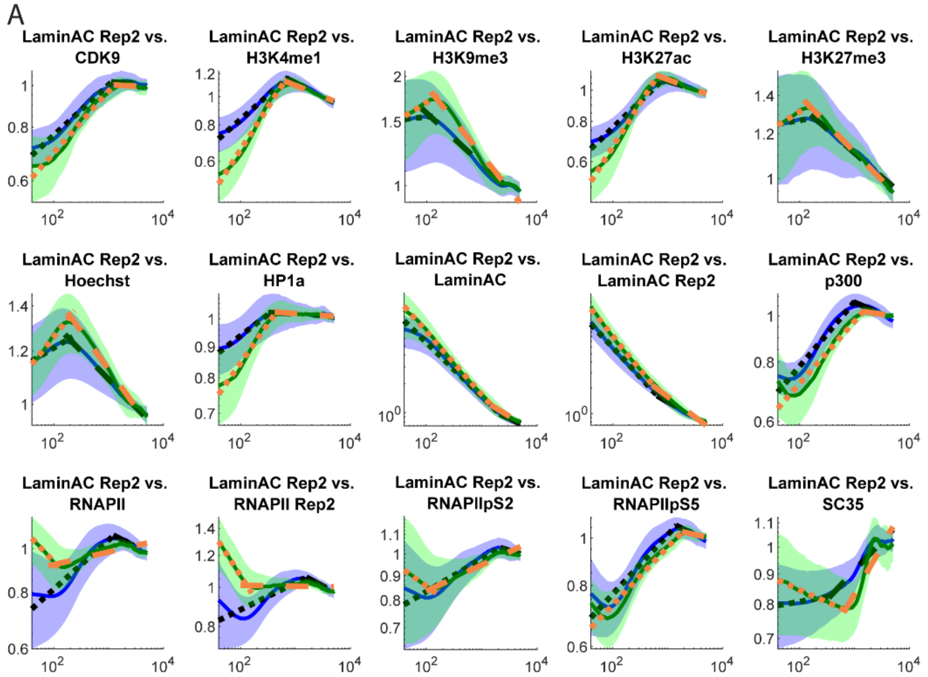

B

**Wilcoxon rank sum control vs. αAM: Lamin A/C repeat vs.**

| | Hinge | Slope 1 | Slope 2 |
|---|---|---|---|
| **CDK9** | 0.551882 | 0.014855 | 0.813641 |
| **H3K4me1** | 0.030274 | 7.50E-06 | 0.18906 |
| **H3K9me3** | 0.090071 | 0.060844 | 0.085892 |
| **H3K27Ac** | 0.057817 | 2.32E-05 | 0.645345 |
| **H3K27me3** | 0.399844 | 0.221118 | 0.973136 |
| **Hoechst** | 0.973136 | 0.005956 | 0.094411 |
| **HP1a** | 0.129666 | 0.001005 | 0.727851 |
| **LaminAC** | 0.613461 | 0.04948 | 0.831108 |
| **LaminAC Rep2** | 0.744776 | 0.18906 | 0.06729 |
| **p300** | 0.016803 | 0.796266 | 0.000481 |
| **RNAPII** | 0.167366 | 2.10E-05 | 0.002531 |
| **RNAPII Rep2** | 0.018971 | 0.000373 | 0.042175 |
| **RNAPIIpS2** | 0.340006 | 0.098918 | 0.425452 |
| **RNAPIIpS5** | 0.711058 | 0.66154 | 0.018971 |
| **SC35** | 0.0073 | 0.000442 | 0.00891 |

**Figure S62. Lamin A/C repeat vs. all, αAM vs. Control PCCF.**

(A) Normalized PCCF curves versus random (blue indicates control, green indicates αAM). (B) Wilcoxon rank sum p-values for each pair of comparisons.

A

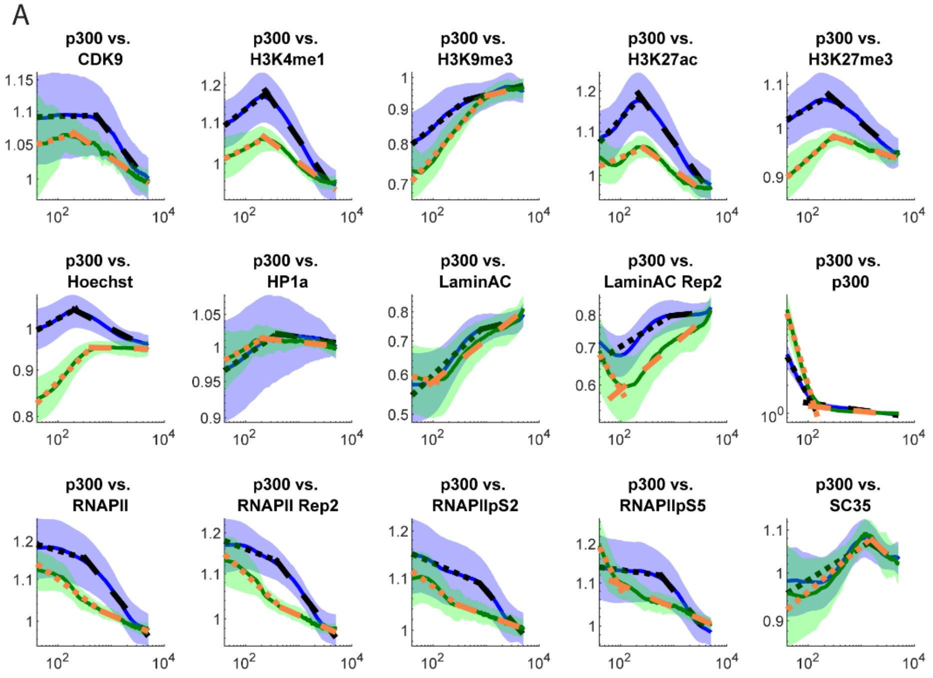

B

**Wilcoxon rank sum control vs. αAM: p300 vs.**

| | Hinge | Slope 1 | Slope 2 |
|---|---|---|---|
| **CDK9** | 0.025494 | 0.296506 | 0.001274 |
| **H3K4me1** | 0.204633 | 0.090071 | 1.84E-06 |
| **H3K9me3** | 0.18161 | 0.014855 | 0.493505 |
| **H3K27Ac** | 0.090071 | 0.001178 | 3.19E-06 |
| **H3K27me3** | 0.21276 | 0.00218 | 6.75E-06 |
| **Hoechst** | 1.04E-06 | 3.19E-06 | 8.31E-07 |
| **HP1a** | 0.003392 | 0.813641 | 0.694403 |
| **LaminAC** | 0.118684 | 0.013109 | 0.001875 |
| **LaminAC Rep2** | 0.054917 | 0.005188 | 6.75E-06 |
| **p300** | 0.000373 | 9.71E-08 | 0.000671 |
| **RNAPII** | 0.094411 | 0.000118 | 1.31E-06 |
| **RNAPII Rep2** | 0.866284 | 0.000288 | 3.26E-07 |
| **RNAPIIpS2** | 0.196734 | 0.118684 | 8.33E-06 |
| **RNAPIIpS5** | 0.074289 | 8.31E-07 | 3.12E-05 |
| **SC35** | 0.677895 | 0.425452 | 0.17438 |

**Figure S63.** p**300 vs. all, αAM vs. Control PCCF.**

(A) Normalized PCCF curves versus random (blue indicates control, green indicates αAM). (B) Wilcoxon rank sum p-values for each pair of comparisons.

A

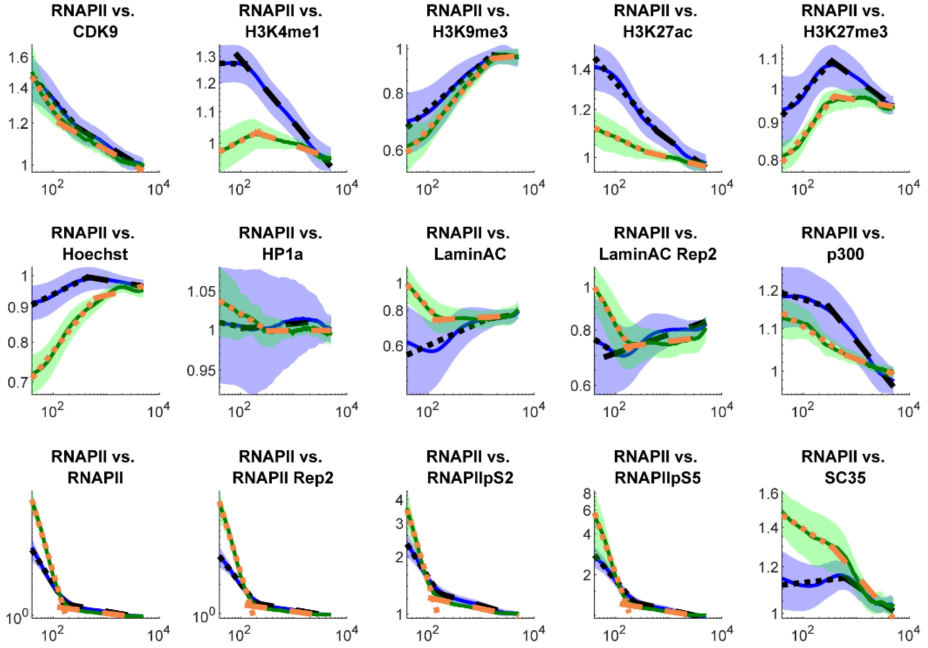

B

Wilcoxon rank sum control vs. αAM: RNAPII vs.

| | Hinge | Slope 1 | Slope 2 |
|---|---|---|---|
| CDK9 | 0.057817 | 0.108445 | 0.645345 |
| H3K4me1 | 0.010152 | 1.10E-07 | 7.41E-07 |
| H3K9me3 | 0.613461 | 0.010152 | 0.103594 |
| H3K27Ac | 0.001737 | 0.001378 | 0.160567 |
| H3K27me3 | 0.113474 | 0.536975 | 0.000129 |
| Hoechst | 0.039953 | 3.13E-08 | 0.00891 |
| HP1a | 0.582291 | 0.006377 | 0.613461 |
| LaminAC | 0.135445 | 6.07E-06 | 0.991044 |
| LaminAC Rep2 | 0.317768 | 1.47E-06 | 0.21276 |
| p300 | 0.016803 | 0.000108 | 8.31E-07 |
| RNAPII | 0.042175 | 2.75E-08 | 3.44E-05 |
| RNAPII Rep2 | 3.96E-06 | 2.75E-08 | 7.42E-05 |
| RNAPIIpS2 | 7.41E-07 | 7.58E-08 | 0.000373 |
| RNAPIIpS5 | 0.001005 | 5.07E-05 | 4.18E-05 |
| SC35 | 0.001737 | 6.07E-06 | 0.070719 |

**Figure S64. RNAPII vs. all, αAM vs. Control PCCF.**

(A) Normalized PCCF curves versus random (blue indicates control, green indicates αAM). (B) Wilcoxon rank sum p-values for each pair of comparisons.

A

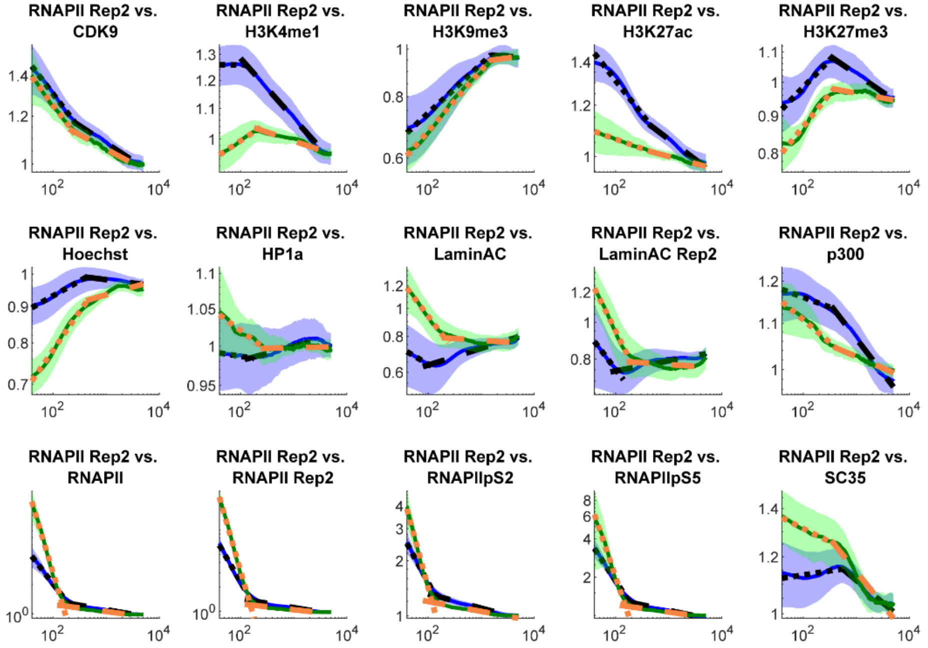

B

Wilcoxon rank sum control vs. αAM: RNAPII repeat vs.

| | Hinge | Slope 1 | Slope 2 |
|---|---|---|---|
| CDK9 | 0.28624 | 0.937369 | 0.522274 |
| H3K4me1 | 0.001089 | 7.58E-08 | 5.90E-08 |
| H3K9me3 | 0.597784 | 0.078006 | 0.141421 |
| H3K27Ac | 0.438607 | 2.56E-06 | 0.00556 |
| H3K27me3 | 0.438607 | 0.04948 | 2.56E-06 |
| Hoechst | 0.196734 | 3.13E-08 | 0.003392 |
| HP1a | 0.081872 | 0.016803 | 0.296506 |
| LaminAC | 0.465606 | 1.03E-05 | 0.070719 |
| LaminAC Rep2 | 0.919528 | 0.000928 | 0.387395 |
| p300 | 0.522274 | 0.002933 | 2.90E-07 |
| RNAPII | 3.19E-06 | 2.75E-08 | 0.000242 |
| RNAPII Rep2 | 0.28624 | 2.75E-08 | 2.86E-06 |
| RNAPIIpS2 | 2.32E-05 | 3.56E-08 | 2.56E-05 |
| RNAPIIpS5 | 0.375186 | 0.108445 | 9.32E-07 |
| SC35 | 0.030274 | 2.32E-05 | 0.256896 |

**Figure S65. RNAPII repeat vs. all, αAM vs. Control PCCF.**

(A) Normalized PCCF curves versus random (blue indicates control, green indicates αAM). (B) Wilcoxon rank sum p-values for each pair of comparisons.

A

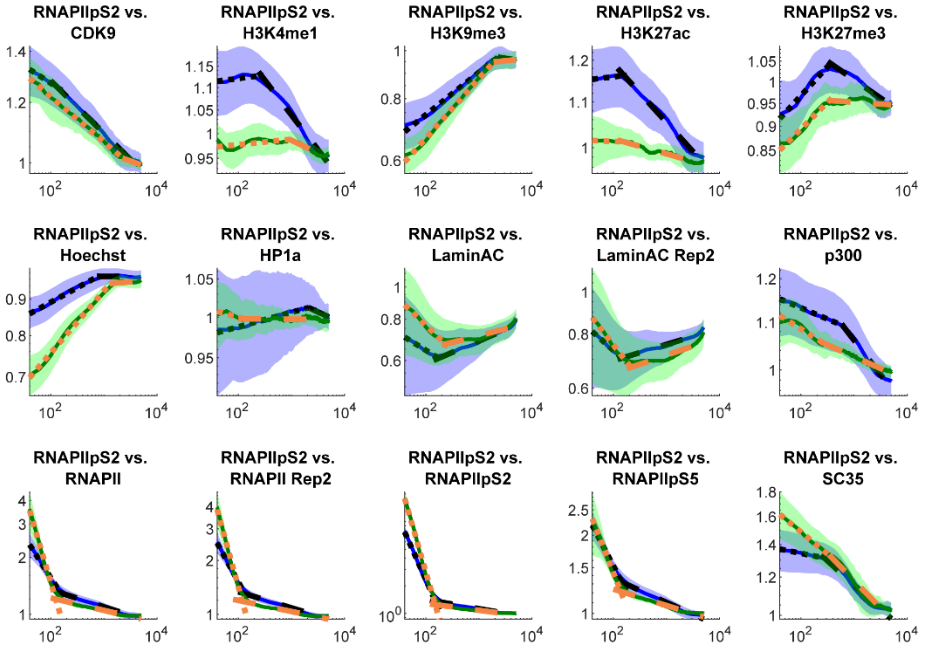

B

**Wilcoxon rank sum control vs. αAM: RNAPII pS2 vs.**

| | Hinge | Slope 1 | Slope 2 |
|---|---|---|---|
| **CDK9** | 0.901728 | 0.551882 | 0.167366 |
| **H3K4me1** | 0.991044 | 0.629316 | 2.28E-07 |
| **H3K9me3** | 0.955241 | 0.000264 | 0.645345 |
| **H3K27Ac** | 0.028601 | 0.991044 | 1.31E-06 |
| **H3K27me3** | 0.677895 | 0.536975 | 7.42E-05 |
| **Hoechst** | 0.098918 | 6.69E-08 | 0.135445 |
| **HP1a** | 0.060844 | 0.340006 | 0.363217 |
| **LaminAC** | 0.848658 | 0.027009 | 0.465606 |
| **LaminAC Rep2** | 0.046935 | 0.317768 | 0.465606 |
| **p300** | 0.645345 | 0.113474 | 3.19E-06 |
| **RNAPII** | 2.29E-06 | 5.90E-08 | 8.94E-05 |
| **RNAPII Rep2** | 0.000222 | 3.56E-08 | 3.55E-06 |
| **RNAPIIpS2** | 0.883977 | 2.75E-08 | 0.000373 |
| **RNAPIIpS5** | 0.919528 | 0.247596 | 4.41E-06 |
| **SC35** | 0.00218 | 4.41E-06 | 0.465606 |

**Figure S66. RNAPII pS2 vs. all, αAM vs. Control PCCF.**

(A) Normalized PCCF curves versus random (blue indicates control, green indicates αAM). (B) Wilcoxon rank sum p-values for each pair of comparisons.

A

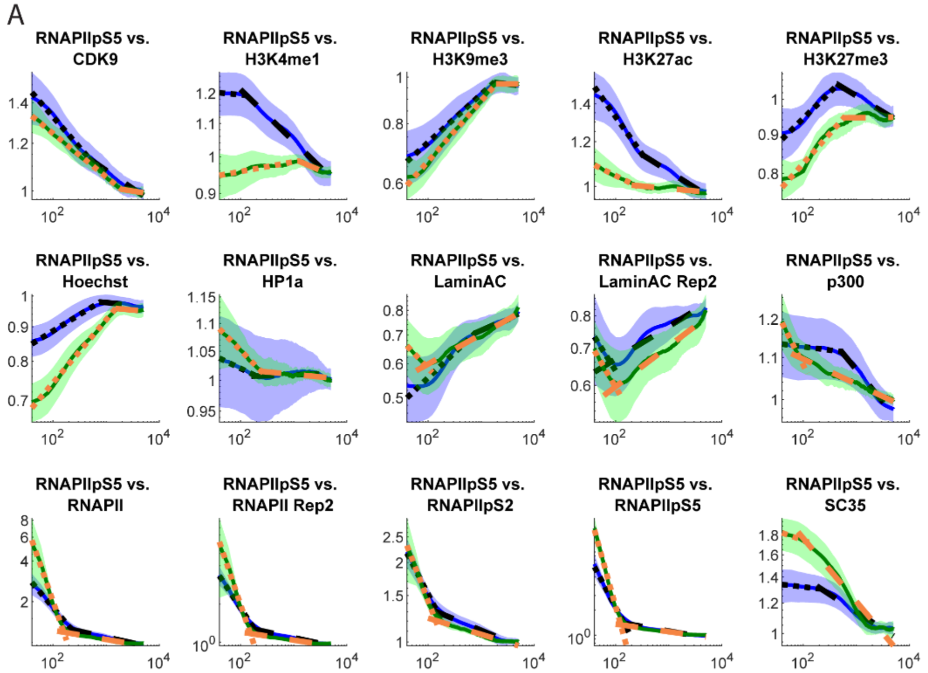

B

**Wilcoxon rank sum control vs. αAM: RNAPII pS5 vs.**

| | Hinge | Slope 1 | Slope 2 |
|---|---|---|---|
| **CDK9** | 0.03203 | 0.18906 | 0.711058 |
| **H3K4me1** | 0.044502 | 2.32E-05 | 4.91E-06 |
| **H3K9me3** | 0.597784 | 0.035805 | 0.238534 |
| **H3K27Ac** | 0.340006 | 8.33E-06 | 1.47E-06 |
| **H3K27me3** | 0.141421 | 0.06729 | 3.79E-05 |
| **Hoechst** | 0.046935 | 2.05E-06 | 0.425452 |
| **HP1a** | 0.694403 | 0.017858 | 0.074289 |
| **LaminAC** | 0.000856 | 5.23E-07 | 0.000314 |
| **LaminAC Rep2** | 0.955241 | 0.103594 | 0.000118 |
| **p300** | 0.004839 | 5.46E-06 | 5.46E-06 |
| **RNAPII** | 0.002022 | 3.79E-05 | 0.000222 |
| **RNAPII Rep2** | 0.399844 | 0.103594 | 1.47E-06 |
| **RNAPIIpS2** | 0.866284 | 0.221118 | 5.07E-05 |
| **RNAPIIpS5** | 1.71E-05 | 2.75E-08 | 0.006377 |
| **SC35** | 0.078006 | 4.61E-05 | 0.064 |

**Figure S67. RNAPII pS5 vs. all, αAM vs. Control PCCF.**

(A) Normalized PCCF curves versus random (blue indicates control, green indicates αAM). (B) Wilcoxon rank sum p-values for each pair of comparisons.

A

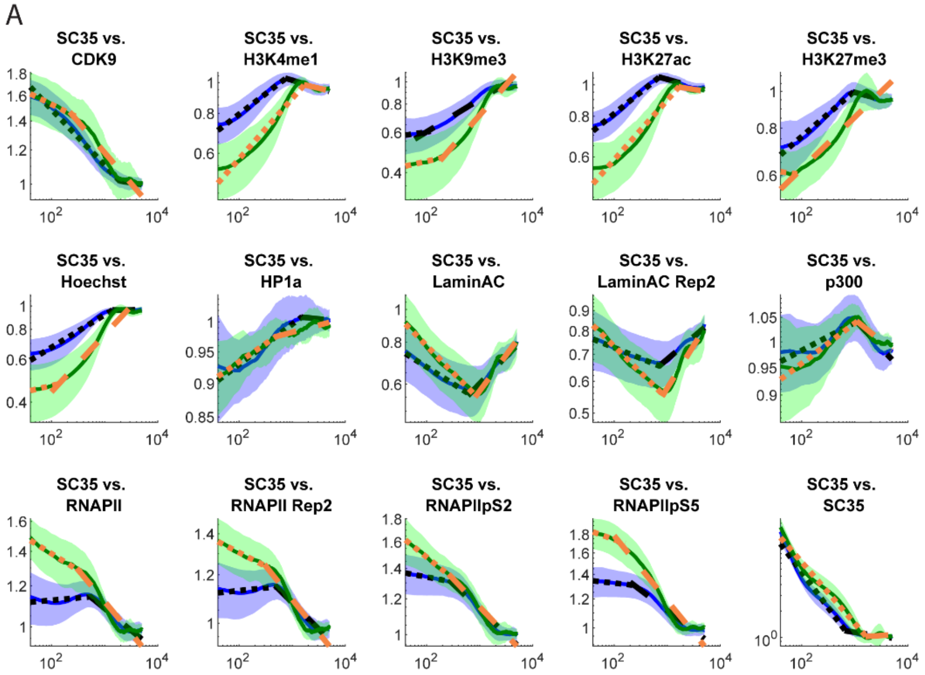

B

**Wilcoxon rank sum control vs. αAM: SC-35 vs.**

| | Hinge | Slope 1 | Slope 2 |
|---|---|---|---|
| **CDK9** | 0.081872 | 0.008341 | 0.004839 |
| **H3K4me1** | 0.007805 | 0.28624 | 0.085892 |
| **H3K9me3** | 0.694403 | 0.340006 | 0.000185 |
| **H3K27Ac** | 0.000856 | 0.196734 | 0.108445 |
| **H3K27me3** | 0.425452 | 0.078006 | 0.011547 |
| **Hoechst** | 0.000569 | 0.000728 | 1.71E-05 |
| **HP1a** | 0.028601 | 0.831108 | 0.17438 |
| **LaminAC** | 0.022682 | 0.002933 | 0.141421 |
| **LaminAC Rep2** | 0.129666 | 1.90E-05 | 0.002531 |
| **p300** | 0.479445 | 0.307015 | 0.317768 |
| **RNAPII** | 0.024052 | 5.46E-06 | 0.160567 |
| **RNAPII Rep2** | 0.001005 | 9.25E-06 | 0.108445 |
| **RNAPIIpS2** | 0.081872 | 2.86E-06 | 0.883977 |
| **RNAPIIpS5** | 0.0073 | 0.000129 | 0.002531 |
| **SC35** | 0.167366 | 0.266436 | 0.35149 |

**Figure S68. SC-35 vs. all, αAM vs. Control PCCF.**

(A) Normalized PCCF curves versus random (blue indicates control, green indicates αAM). (B) Wilcoxon rank sum p-values for each pair of comparisons.

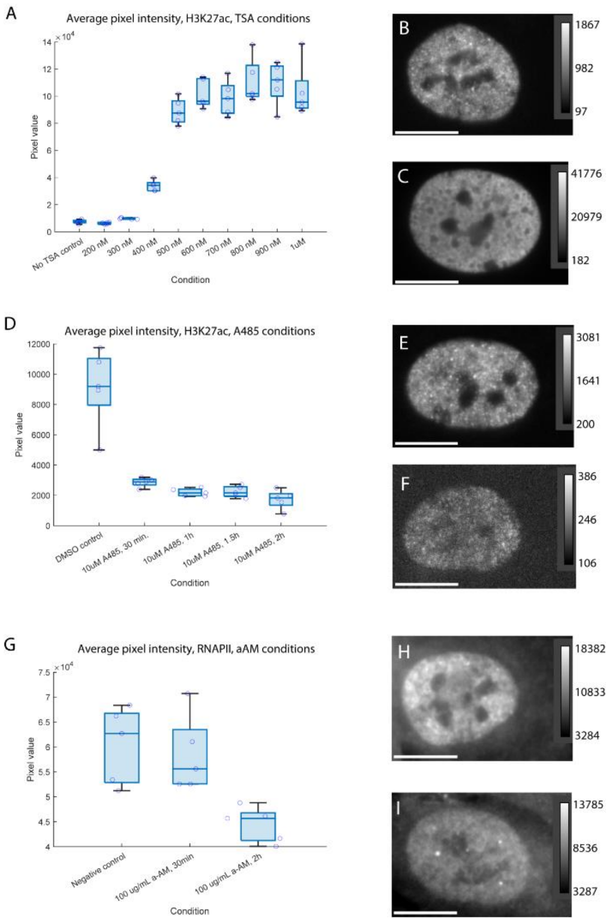

**Figure S69. Widefield imaging and quantification of perturbation effects.**

(A) Average pixel intensity of H3K27ac within nuclear masks after labeling with anti-rabbit FluoTag 488 (FT488) at varying concentrations of Trichostatin A (TSA), treated for twenty hours immediately prior to fixation. Nuclear masks were generated with manual thresholding of DAPI costain. Points represent individual fields of view (n=5). (B) Representative nucleus labeled with H3K27ac-FT488 on an untreated sample. Scale bar represents 10 µm. (C) Representative nucleus labeled with H3K27ac-FT488 on a sample treated with 1 µM TSA for twenty hours. Scale bar represents 10 µm. (D) Average pixel intensity of H3K27ac within nuclear masks after labeling with anti-rabbit FluoTag 488 (FT488) for varying time treatments with 10 µM A-485, treated immediately prior to fixation. Points represent individual fields of view (n=5). (E) Representative nucleus labeled with H3K27ac-FT488 on a vehicle (DMSO) control sample. Scale bar represents 10 µm. (F) Representative nucleus labeled with H3K27ac-FT488 on a sample treated with 10 µM A-485 for two hours. Scale bar represents 10 µm. (G) Average pixel intensity within nuclear masks of RNA Polymerase II (RNAPII) after labeling with anti-mouse FluoTag 488 (FT488) at varying concentrations, treated for two hours immediately prior to fixation with a-amanitin. Points represent individual fields of view (n=5). (H) Representative nucleus labeled with RNAPII-FT488 on a vehicle (DMSO) control sample. Scale bar represents 10 µm. (I) Representative nucleus labeled with RNAPII-FT488 on a sample treated with 100 µg/mL α-amanitin for two hours. Scale bar represents 10 µm.

| Target | Manufacturer | Catalog Number | Primary antibody working concent-ration (μg/mL) | PAINT Probe ID | Docking Strand Sequence | Imager Strand Sequence | Imager Strand Fluorophore | Imager strand concentration |
|---|---|---|---|---|---|---|---|---|
| **CDK9** | Abcam | 76320 | 0.36 | P5 | TTTCAATGTAT | TACATTGA | ATTO655 | 5 nM |
| **H3K4me1** | Abcam | 8895 | 0.34 | P13 | TTATAGAGAAG | CTTCTCTAT | ATTO655 | 1 nM |
| **H3K9me3** | Abcam | 176916 | 3.77 | P8 | TTATGTTAATG | CATTAACAT | Cy3B | 0.25 nM |
| **H3K27ac** | Abcam | 177178 | 4.69 | P4 | TTATGAATCTA | AGATTCAT | Cy3B | 2.5 nM (control) or less, as needed for single molecule localization |
| **H3K27me3** | Cell Signaling Technologies | 9733 | 0.34 | P9 | TTAATTAGGAT | TCCTAATT | ATTO655 | 1 nM |
| **HP1a** | Abcam | 109028 | 1.03 | P3 | TTTCTTCATTA | AATGAAGA | Cy3B | 2.5 nM |
| **Lamin A/C** | Santa Cruz | 376248 | 0.667 | P1 | TTATACATCTA | AGATGTAT | Cy3B | 2.5 nM |
| **p300** | Abcam | 275378 | 1 | P17 | TTCCTAAAATT | AATTTTAGG | Cy3B | 2.5 nM |
| **RNA Polymerase II (RNAPII)** | Santa Cruz | 47701 | 0.667 | P24 | TTATAATCATG | CATGATTAT | ATTO655 | 1 nM |
| **RNAPII S2P** | Abcam | 5095 | 3.33 | P19 | TTATATGATCT | GATCATAT | ATTO655 | 1 nM |
| **RNAPII S5P** | Abcam | 5131 | 3 | P16 | TTCGTTTAATT | ATTAAACG | Cy3B | 2.5 nM |
| **SC-35** | Abcam | 11826 | 5 | P20 | TTTATTAAGCT | GCTTAATA | ATTO655 | 2.5 nM |

**Table S1. Primary antibody and DNA-PAINT probe characteristics.**

| **Hinge point (mean)** | | | | | | | | | | | | | | | |
|---|---|---|---|---|---|---|---|---|---|---|---|---|---|---|---|
| | **CDK9** | **H3K4me1** | **H3K9me3** | **H3K27Ac** | **H3K27me3** | **Hoechst** | **HP1a** | **LaminAC** | **LaminAC_** | **p300** | **RNAPII** | **RNAPII_Repeat** | **RNAPIIpS2** | **RNAPIIpS5** | **SC35** |
| **CDK9** | 158.5138 | 526.1587 | 946.5467 | 543.602 | 566.58594 | 589.7036 | 317.5664 | 718.8677 | 572.7211 | 516.5918 | 419.5546567 | 950.1423892 | 750.11529 | 827.7827044 | 1175.658 |
| **H3K4me1** | 824.081 | 172.8786 | 731.8097 | 379.0874 | 432.92194 | 763.1269 | 841.2191 | 647.7429 | 663.4339 | 253.5937 | 383.3860524 | 374.6436864 | 630.90448 | 1085.891128 | 916.4533 |
| **H3K9me3** | 1418.788 | 932.4586 | 755.436 | 1335.976 | 843.61921 | 1205.802 | 1101.184 | 987.9534 | 1150.19 | 838.3301 | 1258.768341 | 1230.814699 | 4739.6308 | 1390.988407 | 765.079 |
| **H3K27Ac** | 565.1892 | 455.9915 | 1955.03 | 173.1408 | 413.01066 | 746.3368 | 351.866 | 575.3034 | 810.6623 | 243.8073 | 672.2783112 | 612.943143 | 622.35403 | 357.0785321 | 805.9193 |
| **H3K27me3** | 629.4212 | 443.1834 | 830.4978 | 469.9164 | 233.34868 | 752.1509 | 564.7868 | 958.5516 | 1100.275 | 417.9154 | 397.2817171 | 413.453337 | 541.96367 | 519.5978447 | 1340.625 |
| **Hoechst** | 886.4922 | 705.6366 | 1158.019 | 672.497 | 477.78572 | 1137.137 | 1275.519 | 823.5046 | 1091.017 | 331.9883 | 713.2760976 | 716.7584998 | 1217.2707 | 1073.861232 | 1358.498 |
| **HP1a** | 436.3834 | 719.2316 | 1177.04 | 504.4498 | 591.4897 | 555.5315 | 126.162 | 1249.202 | 1174.685 | 675.3701 | 1215.775369 | 1187.37794 | 1046.3962 | 930.4469648 | 1275.072 |
| **LaminAC** | 1194.244 | 783.3544 | 741.7939 | 1225.724 | 607.43405 | 333.6157 | 606.7121 | 949.2696 | 947.2901 | 943.9657 | 1099.911826 | 1166.513377 | 27307.681 | 1098.605981 | 475.9424 |
| **LaminAC_Repeat** | 1060.883 | 816.309 | 989.9858 | 809.532 | 550.74656 | 375.1444 | 609.4829 | 872.8962 | 889.5143 | 976.3542 | 1152.352194 | 1005.586478 | 2802.868 | 1293.236511 | 471.3379 |
| **p300** | 575.2883 | 257.0014 | 669.4739 | 241.8823 | 264.48088 | 214.2021 | 889.9371 | 737.5314 | 1006.559 | 121.7829 | 473.6120482 | 524.942198 | 772.25162 | 571.5567233 | 1297.707 |
| **RNAPII** | 361.6851 | 230.0492 | 950.7614 | 1148.47 | 362.96607 | 529.6629 | 524.7414 | 796.347 | 741.094 | 375.9723 | 180.7635042 | 198.3278659 | 163.4091 | 230.3599005 | 721.0136 |
| **RNAPII_Repeat** | 570.6201 | 206.0653 | 936.2083 | 879.3821 | 372.83075 | 504.9066 | 801.8293 | 754.5444 | 1109.03 | 495.9953 | 197.9652727 | 162.1193571 | 141.40935 | 188.5222895 | 678.9064 |
| **RNAPIIpS2** | 713.6228 | 375.4726 | 986.216 | 363.4274 | 394.22179 | 859.7128 | 616.9924 | 385.2078 | 490.2727 | 708.6814 | 158.2294479 | 138.5735396 | 151.73624 | 152.090612 | 510.6515 |
| **RNAPIIpS5** | 815.1765 | 524.7762 | 2362.735 | 356.1561 | 472.39064 | 852.6721 | 399.833 | 1127.959 | 561.6754 | 541.3253 | 228.6675802 | 187.3377436 | 155.51802 | 150.2309195 | 468.3111 |
| **SC35** | 1009.038 | 747.0323 | 938.338 | 686.0205 | 865.71052 | 1326.198 | 1053.524 | 704.7616 | 801.1276 | 979.1081 | 573.5578884 | 575.3545117 | 360.80548 | 382.9472847 | 716.0968 |
| **Hinge point (std)** | | | | | | | | | | | | | | | |
| | **CDK9** | **H3K4me1** | **H3K9me3** | **H3K27Ac** | **H3K27me3** | **Hoechst** | **HP1a** | **LaminAC** | **LaminAC_** | **p300** | **RNAPII** | **RNAPII_Repeat** | **RNAPIIpS2** | **RNAPIIpS5** | **SC35** |
| **CDK9** | 13.42878 | 438.6285 | 746.3281 | 666.8347 | 234.44137 | 235.881 | 446.1475 | 775.2112 | 573.0066 | 204.6105 | 470.1579556 | 1620.273959 | 822.93588 | 815.2400908 | 849.3367 |
| **H3K4me1** | 556.6855 | 37.70591 | 689.5495 | 84.39743 | 113.45993 | 668.9765 | 926.5296 | 245.2848 | 254.893 | 40.30187 | 613.2093315 | 494.2375254 | 549.74243 | 863.6186196 | 296.8133 |
| **H3K9me3** | 492.4887 | 554.6687 | 456.2442 | 2098.514 | 505.65324 | 531.1881 | 510.2525 | 829.3655 | 911.3562 | 521.1011 | 391.1999644 | 489.6692544 | 17974.026 | 467.5810835 | 850.0496 |
| **H3K27Ac** | 669.124 | 433.2078 | 8277.014 | 29.65163 | 83.382667 | 715.3552 | 336.5565 | 222.4326 | 509.707 | 36.89457 | 652.8341408 | 592.358041 | 760.18129 | 259.6091517 | 277.6247 |
| **H3K27me3** | 292.2437 | 106.351 | 735.8558 | 226.2279 | 44.117356 | 482.1277 | 305.2264 | 655.7601 | 874.4877 | 381.041 | 124.7842267 | 143.6735213 | 310.54148 | 148.0921454 | 1780.894 |
| **Hoechst** | 261.1377 | 856.1042 | 476.5196 | 545.59 | 344.2653 | 508.2054 | 2123.817 | 913.417 | 1038.812 | 113.7859 | 230.9011726 | 265.0576437 | 393.12511 | 279.6183797 | 618.0539 |
| **HP1a** | 580.3056 | 547.8843 | 449.5102 | 535.5631 | 255.44154 | 802.3727 | 12.47089 | 909.712 | 1048.695 | 399.4626 | 2025.142466 | 1105.807109 | 700.13791 | 987.4119019 | 1577.244 |
| **LaminAC** | 1099.666 | 246.8306 | 877.8068 | 2474.991 | 863.80601 | 405.433 | 668.9011 | 804.5573 | 929.9717 | 397.2277 | 392.778479 | 577.6458735 | 144968.92 | 426.2050285 | 254.9067 |
| **LaminAC_Repeat** | 590.8503 | 278.1482 | 1085.859 | 368.0899 | 651.08204 | 487.0653 | 757.7351 | 811.1309 | 743.6148 | 398.2791 | 456.6861041 | 778.8996935 | 7590.2691 | 539.9991085 | 342.6602 |
| **p300** | 273.0351 | 43.17401 | 629.9199 | 39.61298 | 76.272671 | 55.05968 | 1450.084 | 789.1029 | 979.8471 | 5.56424 | 411.7228617 | 451.4828474 | 569.57609 | 226.6148813 | 1123.609 |
| **RNAPII** | 375.1656 | 331.3589 | 723.0723 | 961.9112 | 108.85683 | 294.7478 | 438.6049 | 874.911 | 1120.388 | 319.2626 | 31.42917826 | 30.53753934 | 28.552717 | 26.95223841 | 366.4962 |
| **RNAPII_Repeat** | 745.3403 | 198.3881 | 763.4664 | 871.0474 | 109.51598 | 303.3581 | 866.2517 | 1117.628 | 2555.416 | 441.9175 | 29.39977809 | 19.11960969 | 19.356151 | 25.67800179 | 409.9308 |
| **RNAPIIpS2** | 816.1244 | 226.9797 | 753.0683 | 510.14 | 161.50214 | 563.0608 | 661.7757 | 370.1098 | 1003.406 | 913.5617 | 26.72580077 | 19.85915669 | 14.210964 | 25.66922147 | 631.375 |
| **RNAPIIpS5** | 765.9044 | 479.839 | 8712.832 | 201.3469 | 125.7775 | 418.2021 | 355.6436 | 911.4133 | 680.2985 | 152.3241 | 26.04898799 | 24.15016246 | 24.669658 | 21.57508528 | 588.7804 |
| **SC35** | 811.0409 | 195.9327 | 959.6523 | 211.38 | 306.05346 | 387.0055 | 885.852 | 475.2479 | 482.2887 | 385.3556 | 167.2543709 | 278.6518403 | 256.27254 | 545.8689731 | 367.3924 |

**Table S2. Hinge points of the PCCF curves for control cells.**

| Slope 1 (mean) | | | | | | | | | | | | | | | |
|---|---|---|---|---|---|---|---|---|---|---|---|---|---|---|---|
| | **CDK9** | **H3K4me1** | **H3K9me3** | **H3K27Ac** | **H3K27me3** | **Hoechst** | **HP1a** | **LaminAC** | **LaminAC_** | **p300** | **RNAPII** | **RNAPII_Repeat** | **RNAPIIpS2** | **RNAPIIpS5** | **SC35** |
| **CDK9** | -1.66325 | -0.00588 | 0.075166 | -0.06146 | 0.0528251 | 0.055342 | -0.05107 | 0.020865 | 0.012312 | 0.002754 | -0.113238965 | -0.117133323 | -0.054979 | -0.09859188 | -0.10704 |
| **H3K4me1** | -0.00394 | -0.69996 | 0.006483 | -0.1958 | 0.0138736 | -0.02291 | -0.01695 | 0.133109 | 0.093893 | 0.042057 | -0.013236177 | -0.012029549 | 0.0061355 | -0.02961592 | 0.13978 |
| **H3K9me3** | 0.098082 | 0.033791 | -0.64108 | 0.017131 | 0.0188199 | -0.11003 | -0.09796 | -0.1494 | -0.11793 | 0.08688 | 0.12823997 | 0.129472141 | 0.1028022 | 0.124794162 | 0.085162 |
| **H3K27Ac** | -0.06714 | -0.19722 | -0.0183 | -0.98809 | 0.0517735 | -0.01927 | -0.04305 | 0.115269 | 0.08481 | 0.055872 | -0.094075529 | -0.100725838 | -0.00107 | -0.12845549 | 0.142768 |
| **H3K27me3** | 0.065158 | 0.022754 | 0.009885 | 0.062471 | -0.933535 | -0.0544 | 0.048036 | -0.12122 | -0.08465 | 0.035788 | 0.093364326 | 0.085280193 | 0.0696955 | 0.07901549 | 0.140702 |
| **Hoechst** | 0.067804 | -0.01345 | -0.11659 | -0.00909 | -0.052742 | -0.0983 | 0.000435 | -0.07566 | -0.04717 | 0.036178 | 0.050428491 | 0.051493548 | 0.0566364 | 0.061657434 | 0.144609 |
| **HP1a** | -0.0444 | -0.01453 | -0.11638 | -0.03861 | 0.0419145 | -0.00107 | -1.57183 | -0.03983 | -0.01658 | 0.029471 | 0.003214097 | 0.006137136 | 0.0116556 | -0.01093069 | 0.015601 |
| **LaminAC** | 0.169634 | 0.241528 | -0.02113 | 0.233389 | 0.0144577 | 0.046332 | 0.083015 | -0.69092 | -0.37396 | 0.217754 | 0.207349252 | 0.132578016 | 0.0701598 | 0.226847877 | -0.005 |
| **LaminAC_Repeat** | 0.115228 | 0.168671 | -0.02627 | 0.162317 | 0.0168999 | 0.051219 | 0.07159 | -0.41236 | -0.70324 | 0.134219 | 0.113532884 | -0.003428486 | 0.0302814 | 0.11801005 | 0.011898 |
| **p300** | 0.006684 | 0.041238 | 0.072436 | 0.055518 | 0.0290843 | 0.031739 | 0.027066 | 0.099514 | 0.03861 | -1.02095 | -0.013833868 | -0.016249486 | -0.018361 | -0.0047838 | 0.022753 |
| **RNAPII** | -0.11127 | -0.00869 | 0.102118 | -0.09554 | 0.0843246 | 0.039447 | -0.00618 | 0.052346 | -0.01589 | -0.01278 | -0.984828177 | -0.699953234 | -0.435837 | -0.45963439 | 0.012815 |
| **RNAPII_Repeat** | -0.1148 | -0.00627 | 0.112584 | -0.10021 | 0.0773827 | 0.043199 | -0.00232 | -0.02021 | -0.11666 | -0.014 | -0.697831421 | -1.134326073 | -0.532478 | -0.6315299 | 0.01211 |
| **RNAPIIpS2** | -0.05274 | 0.005874 | 0.072758 | 0.001431 | 0.0632804 | 0.043059 | 0.005294 | -0.06787 | -0.08963 | -0.0164 | -0.437189648 | -0.535123204 | -1.308035 | -0.3889753 | -0.01428 |
| **RNAPIIpS5** | -0.09798 | -0.02216 | 0.094877 | -0.12634 | 0.0697464 | 0.049981 | -0.0194 | 0.121307 | -0.06926 | -0.00587 | -0.45999996 | -0.633053677 | -0.387273 | -1.19229799 | -0.01489 |
| **SC35** | -0.10387 | 0.132775 | 0.074195 | 0.140417 | 0.1266421 | 0.135057 | 0.016562 | -0.10481 | -0.05467 | 0.023825 | 0.011817806 | 0.009539867 | -0.012321 | -0.01335247 | -0.93782 |
| **Slope 1 (std)** | | | | | | | | | | | | | | | |
| | **CDK9** | **H3K4me1** | **H3K9me3** | **H3K27Ac** | **H3K27me3** | **Hoechst** | **HP1a** | **LaminAC** | **LaminAC_** | **p300** | **RNAPII** | **RNAPII_Repeat** | **RNAPIIpS2** | **RNAPIIpS5** | **SC35** |
| **CDK9** | 0.188697 | 0.030275 | 0.041977 | 0.039975 | 0.0496908 | 0.01438 | 0.02908 | 0.082809 | 0.042495 | 0.01589 | 0.031487976 | 0.0375811 | 0.0356253 | 0.035520229 | 0.0483 |
| **H3K4me1** | 0.030508 | 0.169237 | 0.057284 | 0.032509 | 0.0189738 | 0.012613 | 0.022654 | 0.081534 | 0.04292 | 0.015438 | 0.026380185 | 0.025943245 | 0.0200259 | 0.022415212 | 0.038306 |
| **H3K9me3** | 0.05715 | 0.058339 | 0.195726 | 0.069539 | 0.0431475 | 0.036094 | 0.049843 | 0.180655 | 0.144131 | 0.033861 | 0.051793881 | 0.056301028 | 0.0467932 | 0.046376722 | 0.090015 |
| **H3K27Ac** | 0.037896 | 0.03097 | 0.057392 | 0.302999 | 0.0200752 | 0.009903 | 0.023049 | 0.084072 | 0.043873 | 0.024522 | 0.035752168 | 0.032248497 | 0.0257987 | 0.036523961 | 0.034173 |
| **H3K27me3** | 0.050453 | 0.018137 | 0.047562 | 0.019882 | 0.3630755 | 0.015127 | 0.02289 | 0.072429 | 0.068618 | 0.020323 | 0.040218137 | 0.035409128 | 0.0278822 | 0.023163953 | 0.071185 |
| **Hoechst** | 0.014458 | 0.012251 | 0.036245 | 0.009727 | 0.0146498 | 0.026429 | 0.01432 | 0.082117 | 0.067778 | 0.012584 | 0.012676086 | 0.016108976 | 0.0119422 | 0.014027937 | 0.068476 |
| **HP1a** | 0.033268 | 0.018784 | 0.047109 | 0.024841 | 0.0246991 | 0.018246 | 0.302278 | 0.054076 | 0.037144 | 0.014521 | 0.017959854 | 0.018403594 | 0.0217851 | 0.018724085 | 0.08496 |
| **LaminAC** | 0.068252 | 0.090706 | 0.160286 | 0.084732 | 0.0632146 | 0.045432 | 0.04832 | 0.191264 | 0.254621 | 0.06846 | 0.116234852 | 0.122654652 | 0.1342385 | 0.065450702 | 0.057246 |
| **LaminAC_Repeat** | 0.04455 | 0.055182 | 0.144523 | 0.061871 | 0.0526409 | 0.043303 | 0.043585 | 0.267968 | 0.166659 | 0.046588 | 0.074653267 | 0.177676773 | 0.1143469 | 0.059675111 | 0.053035 |
| **p300** | 0.017597 | 0.015297 | 0.03544 | 0.024323 | 0.01932 | 0.011795 | 0.014888 | 0.081758 | 0.062131 | 0.266159 | 0.014496137 | 0.017724285 | 0.0205773 | 0.012580027 | 0.031483 |
| **RNAPII** | 0.03444 | 0.022722 | 0.059101 | 0.029902 | 0.039408 | 0.012533 | 0.019188 | 0.152827 | 0.102369 | 0.017089 | 0.252725846 | 0.166737444 | 0.1158732 | 0.086250811 | 0.038715 |
| **RNAPII_Repeat** | 0.037529 | 0.020649 | 0.060563 | 0.034231 | 0.0352679 | 0.013105 | 0.022001 | 0.135681 | 0.16507 | 0.018825 | 0.166985043 | 0.158397246 | 0.1193324 | 0.111054676 | 0.036288 |
| **RNAPIIpS2** | 0.033825 | 0.020167 | 0.04158 | 0.023919 | 0.0287128 | 0.010659 | 0.0249 | 0.105199 | 0.09079 | 0.021631 | 0.114757593 | 0.120384955 | 0.1252067 | 0.087784818 | 0.037518 |
| **RNAPIIpS5** | 0.035945 | 0.022105 | 0.042996 | 0.038646 | 0.0231398 | 0.013147 | 0.013822 | 0.052781 | 0.15964 | 0.013263 | 0.086242943 | 0.110120647 | 0.0862811 | 0.256643991 | 0.030181 |
| **SC35** | 0.051723 | 0.033706 | 0.072002 | 0.028689 | 0.0652917 | 0.049096 | 0.084096 | 0.049876 | 0.038216 | 0.029626 | 0.038597265 | 0.037439381 | 0.0359262 | 0.026303018 | 0.345915 |

**Table S3. Short length-scale slopes of the PCCF curves for control cells.**

| Slope 2 (mean) | | | | | | | | | | | | | | | |
|---|---|---|---|---|---|---|---|---|---|---|---|---|---|---|---|
| | **CDK9** | **H3K4me1** | **H3K9me3** | **H3K27Ac** | **H3K27me3** | **Hoechst** | **HP1a** | **LaminAC** | **LaminAC_** | **p300** | **RNAPII** | **RNAPII_Repeat** | **RNAPIIpS2** | **RNAPIIpS5** | **SC35** |
| **CDK9** | -0.08141 | -0.06387 | 0.031344 | -0.04685 | -0.043753 | -0.00588 | -0.00264 | 0.077607 | 0.056037 | -0.05369 | -0.062629929 | -0.046584668 | -0.053351 | -0.05862026 | -0.03679 |
| **H3K4me1** | -0.04662 | -0.11372 | -0.00385 | -0.06953 | -0.067439 | -0.03635 | -0.00446 | -0.03618 | -0.0451 | -0.0661 | -0.06807034 | -0.076034878 | -0.052111 | -0.06542402 | -0.03221 |
| **H3K9me3** | -0.02157 | -0.0386 | -0.09104 | -0.01128 | -0.052171 | -0.0329 | -0.01987 | -0.02154 | -0.00809 | -0.02385 | -0.035520154 | -0.034509202 | -0.015967 | -0.04165598 | 0.089397 |
| **H3K27Ac** | -0.04359 | -0.07613 | 0.031776 | -0.09706 | -0.046832 | -0.02893 | -0.00472 | 0.019603 | 0.00136 | -0.06803 | -0.053943426 | -0.05293965 | -0.054006 | -0.05614946 | -0.0365 |
| **H3K27me3** | -0.03523 | -0.07742 | -0.02298 | -0.04632 | -0.117366 | -0.0243 | -0.0069 | -0.04349 | -0.03859 | -0.0428 | -0.054895693 | -0.052228452 | -0.035273 | -0.04282287 | -0.02471 |
| **Hoechst** | -0.01788 | -0.05794 | -0.02189 | -0.04275 | -0.042615 | -0.03088 | -0.02561 | -0.02977 | -0.03139 | -0.03583 | -0.031317417 | -0.030307647 | -0.022943 | -0.0358973 | -0.00611 |
| **HP1a** | -0.00829 | -0.03612 | -0.00869 | -0.02297 | -0.029211 | -0.02533 | -0.08457 | 0.06967 | 0.039392 | -0.03231 | -0.022892634 | -0.023412355 | -0.019232 | -0.02159549 | -0.004 |
| **LaminAC** | 0.009513 | -0.11566 | -0.10203 | -0.04212 | -0.119336 | -0.10373 | -0.01071 | -0.44824 | -0.4192 | -0.05112 | -0.060536316 | -0.06496469 | 0.0089013 | -0.03928557 | 0.145105 |
| **LaminAC_Repeat** | -0.00958 | -0.10016 | -0.09951 | -0.0539 | -0.10309 | -0.09294 | -0.00988 | -0.38054 | -0.32472 | -0.05127 | -0.06379626 | -0.044070375 | -0.0049 | -0.05312397 | 0.102532 |
| **p300** | -0.04511 | -0.07846 | 0.001311 | -0.06867 | -0.044203 | -0.03099 | -0.00992 | 0.034987 | 0.030936 | -0.07619 | -0.066373979 | -0.064021884 | -0.052611 | -0.06539921 | -0.01984 |
| **RNAPII** | -0.05362 | -0.08199 | 0.005326 | -0.03163 | -0.055478 | -0.01715 | -0.00107 | 0.062866 | 0.02382 | -0.06575 | -0.105520782 | -0.092620083 | -0.079108 | -0.08356288 | -0.0676 |
| **RNAPII_Repeat** | -0.04322 | -0.08395 | 0.00417 | -0.04215 | -0.051664 | -0.01451 | -0.00108 | 0.034673 | 0.032049 | -0.06298 | -0.091360152 | -0.105466864 | -0.077237 | -0.08537642 | -0.07147 |
| **RNAPIIpS2** | -0.05443 | -0.06839 | 0.032655 | -0.05814 | -0.040896 | -0.00611 | -0.00251 | 0.083518 | 0.0527 | -0.05981 | -0.089535095 | -0.088353914 | -0.098901 | -0.09059894 | -0.09608 |
| **RNAPIIpS5** | -0.04992 | -0.07353 | 0.035455 | -0.05854 | -0.044377 | -0.01179 | -0.00085 | 0.034392 | 0.034446 | -0.06515 | -0.085651839 | -0.088939408 | -0.082316 | -0.1083987 | -0.08468 |
| **SC35** | -0.0575 | -0.05272 | 0.088709 | -0.04884 | -0.038578 | 0.004222 | -0.00278 | 0.182101 | 0.130197 | -0.05012 | -0.088779394 | -0.088887081 | -0.113614 | -0.10836718 | -0.08248 |
| **Slope 2 (std)** | | | | | | | | | | | | | | | |
| | **CDK9** | **H3K4me1** | **H3K9me3** | **H3K27Ac** | **H3K27me3** | **Hoechst** | **HP1a** | **LaminAC** | **LaminAC_** | **p300** | **RNAPII** | **RNAPII_Repeat** | **RNAPIIpS2** | **RNAPIIpS5** | **SC35** |
| **CDK9** | 0.022528 | 0.025081 | 0.051072 | 0.043741 | 0.0271038 | 0.016302 | 0.041294 | 0.062046 | 0.044856 | 0.027516 | 0.037664734 | 0.048334261 | 0.0509118 | 0.058740468 | 0.068819 |
| **H3K4me1** | 0.030783 | 0.026593 | 0.072547 | 0.022237 | 0.0308945 | 0.029919 | 0.036513 | 0.053036 | 0.036847 | 0.021343 | 0.036839484 | 0.036556162 | 0.0377924 | 0.052626994 | 0.04651 |
| **H3K9me3** | 0.043861 | 0.052114 | 0.098309 | 0.060685 | 0.0438422 | 0.051566 | 0.046769 | 0.185972 | 0.154874 | 0.031101 | 0.044678438 | 0.041033736 | 0.068446 | 0.049318144 | 0.140271 |
| **H3K27Ac** | 0.042959 | 0.035068 | 0.055021 | 0.021042 | 0.0214244 | 0.026469 | 0.03317 | 0.062283 | 0.052839 | 0.02228 | 0.039404052 | 0.03761816 | 0.0350291 | 0.022757581 | 0.040587 |
| **H3K27me3** | 0.029764 | 0.031027 | 0.058924 | 0.022821 | 0.039151 | 0.02628 | 0.020819 | 0.094054 | 0.077461 | 0.021661 | 0.030647622 | 0.029007477 | 0.0308236 | 0.026144471 | 0.071606 |
| **Hoechst** | 0.022934 | 0.028955 | 0.06001 | 0.020315 | 0.0269722 | 0.036569 | 0.029212 | 0.106369 | 0.083044 | 0.016461 | 0.023487518 | 0.02789636 | 0.0368786 | 0.028955193 | 0.088174 |
| **HP1a** | 0.043805 | 0.030864 | 0.056366 | 0.033975 | 0.0193255 | 0.028101 | 0.030444 | 0.076362 | 0.076902 | 0.02477 | 0.03568958 | 0.027919469 | 0.0408187 | 0.043943101 | 0.068315 |
| **LaminAC** | 0.081966 | 0.079913 | 0.167263 | 0.084333 | 0.084774 | 0.058924 | 0.046421 | 0.158615 | 0.172097 | 0.051736 | 0.096462592 | 0.080641569 | 0.1248246 | 0.057267726 | 0.06854 |
| **LaminAC_Repeat** | 0.066999 | 0.062403 | 0.140384 | 0.055035 | 0.0677673 | 0.042082 | 0.043472 | 0.164018 | 0.108146 | 0.046566 | 0.07300403 | 0.079230141 | 0.0997316 | 0.072357095 | 0.058894 |
| **p300** | 0.029077 | 0.022431 | 0.049288 | 0.022275 | 0.0195618 | 0.016493 | 0.026781 | 0.051412 | 0.099214 | 0.020068 | 0.02810449 | 0.025370708 | 0.0361881 | 0.032645459 | 0.077989 |
| **RNAPII** | 0.03235 | 0.025287 | 0.071874 | 0.055471 | 0.0301175 | 0.020057 | 0.036991 | 0.120888 | 0.051325 | 0.027916 | 0.019609565 | 0.017111814 | 0.020009 | 0.021010792 | 0.051989 |
| **RNAPII_Repeat** | 0.036791 | 0.026971 | 0.066259 | 0.043221 | 0.0251921 | 0.022536 | 0.038546 | 0.057978 | 0.104104 | 0.022536 | 0.01920964 | 0.021936448 | 0.0174698 | 0.015693689 | 0.049294 |
| **RNAPIIpS2** | 0.048526 | 0.026235 | 0.078053 | 0.030022 | 0.0255937 | 0.020981 | 0.036977 | 0.056325 | 0.045932 | 0.033592 | 0.020993471 | 0.018428091 | 0.0224987 | 0.019753135 | 0.046853 |
| **RNAPIIpS5** | 0.057678 | 0.029923 | 0.078333 | 0.024225 | 0.0220648 | 0.019704 | 0.037086 | 0.045018 | 0.050894 | 0.030715 | 0.02259424 | 0.016774151 | 0.0206524 | 0.022640425 | 0.044091 |
| **SC35** | 0.070083 | 0.029564 | 0.116359 | 0.031641 | 0.027873 | 0.032575 | 0.042961 | 0.08679 | 0.068982 | 0.040581 | 0.034236962 | 0.033876353 | 0.0414672 | 0.042803667 | 0.098058 |

**Table S4. Long length-scale slopes of the PCCF curves for control cells.**

| Target | Manufacturer | Catalog Number | Host Species and Clonality | Screening Results |
|---|---|---|---|---|
| **Brd4** | Sigma-Aldrich | HPA015055 | Rabbit polyclonal, affinity purified | Nuclear-localized for FluoTag screen with lower specificity than shown in manufacturer representative images, did not proceed with DNA-PAINT |
| **CDK9** | Abcam | 76320 | Rabbit monoclonal | Nuclear-localized with FluoTag and DNA-PAINT probes |
| **CTCF** | Abcam | 128873 | Rabbit monoclonal | Nuclear-localized with FluoTag and DNA-PAINT probes |
| **EZH2** | Abcam | 228697 | Rabbit polyclonal, affinity purified | Nonspecific stain with FluoTag screen, did not proceed with DNA-PAINT |
| **Fibrillarin** | Santa Cruz | 166001 | Mouse monoclonal | Nonspecific stain with FluoTag screen, did not proceed with DNA-PAINT |
| **Histone H1** | Santa Cruz | 393530 | Mouse monoclonal | Nonspecific stain with FluoTag screen, did not proceed with DNA-PAINT |
| **Histone H2A.Z** | Abcam | 4174 | Rabbit polyclonal, affinity purified | Nuclear-localized for FluoTag screen with lower specificity than shown in manufacturer representative images, did not proceed with DNA-PAINT |
| **Histone H3.3** | Abcam | 176840 | Rabbit monoclonal | Nonspecific stain with FluoTag screen, did not proceed with DNA-PAINT |
| **H3K4me1** | Abcam | 8895 | Rabbit polyclonal, affinity purified | Nuclear-localized with FluoTag and DNA-PAINT probes |
| **H3K9me3** | Abcam | 176916 | Rabbit monoclonal | Nuclear-localized with FluoTag and DNA-PAINT probes |
| **H3K9me3** | Abcam | 8898 | Rabbit polyclonal, affinity purified | Nuclear-localized with FluoTag and DNA-PAINT probes; varying optimal concentration between batches |
| **H3K27ac** | Abcam | 177178 | Rabbit monoclonal | Nuclear-localized with FluoTag and DNA-PAINT probes |
| **H3K27me3** | Cell Signaling Technology | 9733 | Rabbit monoclonal | Nuclear-localized with FluoTag and DNA-PAINT probes |
| **H3K27me3** | Abcam | 192985 | Rabbit monoclonal | Occasional nuclear-specific stain but inconsistent with FluoTag screen, did not proceed with DNA-PAINT |
| **H3K27me3** | Abcam | 6002 | Mouse monoclonal | Nonspecific stain with FluoTag screen, did not proceed with DNA-PAINT |
| **H3K36me3** | Abcam | 9050 | Rabbit polyclonal, affinity purified | Nuclear-localized with FluoTag but not with DNA-PAINT probe |
| **H3K36me3** | Cell Signaling Technology | 4909 | Rabbit monoclonal | Nonspecific stain with FluoTag screen, did not proceed with DNA-PAINT |
| **H4K20me1** | Abcam | 9051 | Rabbit polyclonal, affinity purified | Nuclear-localized for FluoTag and DNA-PAINT probes with slightly lower specificity than other primaries |
| **HP1a** | Abcam | 109028 | Rabbit monoclonal | Nuclear-localized with FluoTag and DNA-PAINT probes |
| **Lamin A/C** | Santa Cruz | 376248 | Mouse monoclonal | Nuclear-localized with FluoTag and DNA-PAINT probes |
| **Med14** | Abcam | 196624 | Rabbit polyclonal, affinity purified | Nonspecific stain with FluoTag screen, did not proceed with DNA-PAINT |
| **p300** | Abcam | 275378 | Rabbit monoclonal | Nuclear-localized with FluoTag and DNA-PAINT probes |
| **Rad21** | Abcam | 154769 | Rabbit polyclonal, affinity purified | Nonspecific stain with FluoTag screen, did not proceed with DNA-PAINT |

| | | | | |
|---|---|---|---|---|
| **RNA Polymerase II (RNAPII)** | Santa Cruz | 47701 | Mouse monoclonal | Nuclear-localized with FluoTag and DNA-PAINT probes |
| **RNAPII S2P** | Abcam | 5095 | Rabbit polyclonal, affinity purified | Nuclear-localized with FluoTag and DNA-PAINT probes |
| **RNAPII S5P** | Abcam | 5131 | Rabbit polyclonal, affinity purified | Nuclear-localized with FluoTag and DNA-PAINT probes |
| **SC-35** | Abcam | 11826 | Mouse monoclonal | Nuclear-localized with FluoTag and DNA-PAINT probes |
| **TRAP80 (Med17)** | Abcam | 155593 | Rabbit polyclonal, affinity purified | Nuclear-localized with FluoTag but not with DNA-PAINT probe |

**Table S5. Primary antibody screening results.** Green is recommended for nuclear DNA-PAINT; yellow indicates intermediate specificity or higher batch variability; and red indicates low nuclear specificity for primarily nuclear-localized targets with DNA-PAINT probe.

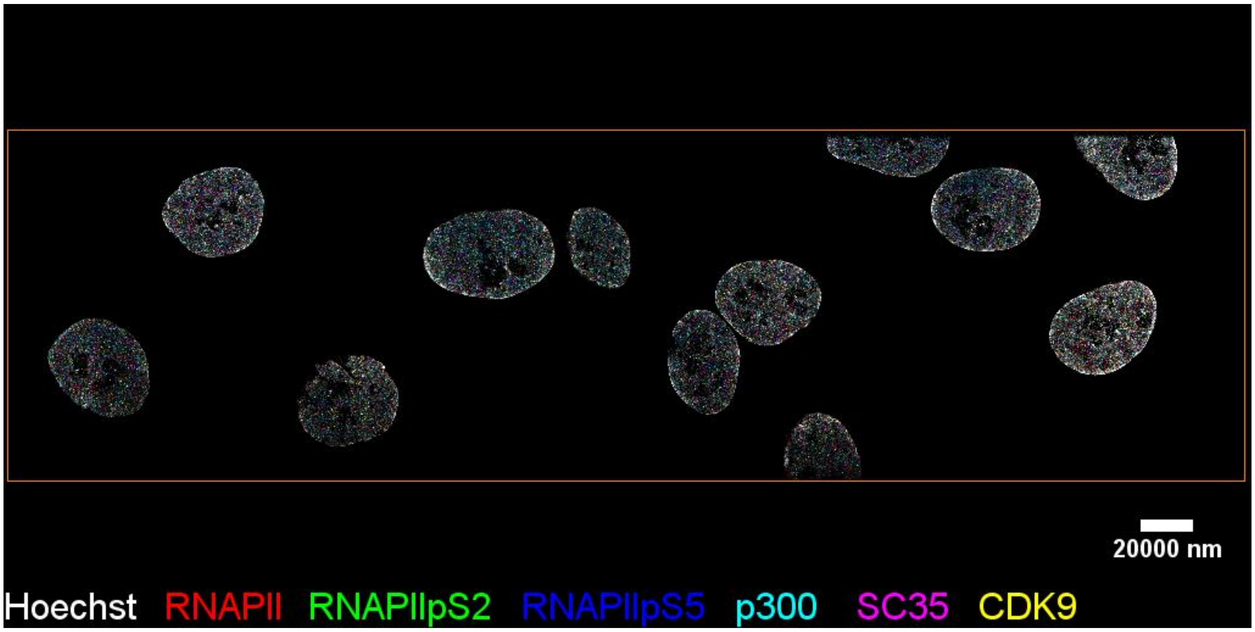

**Movie S1.** Representative multiplex rendering of a single HIST field of view

### Supplementary Note 1. Extended description of Pair Cross-Correlation Function (PCCF) implementation and output

Traditional image analysis methods such as the Pearson correlation coefficient[95] or the Mander's colocalization metric[96] are based on the intensity correlation or the spatial overlap of signal between two species and are commonly used in immunofluorescence microscopy. However, these approaches are less suitable to analyze the data presented here for the following reasons. First, our Exchange-PAINT localization precision approaches the size of the individual molecular targets, resulting in relatively low overlap in the reconstructed super-resolution images, even for two targets that are frequently colocalized together. Second, low signal overlap is further exacerbated due to the limited localization density that arises either from low antibody efficiency or simply from low target densities. And third, traditional co-localization metrics do not capture a length-scale dependence in correlation wherein two targets can be enriched at one length scale and partitioned at another.

For these reasons, we chose to use Ripley's statistics as the primary analysis tool in this manuscript. This method determines the spatial enrichment between two target molecules (or between one target with itself) across spatial scales. Because it averages over all localizations within a dataset, it is able to quantitatively detect subtle trends and differences in the localization patterns between two species even under incomplete labeling density or heterogeneous distributions (e.g. a mix of clustered and random). In contrast to K-means[97] or DBSCAN[98], Ripley's statistics do not require any *a priori* assumptions about the number of clusters present, the number of molecules per cluster, or the cluster size. As such, they are an attractive tool to explore localization datasets in an unbiased manner[99]. However, these same advantages can also pose challenges when interpreting the G(r) and PCCF(r) curves. For example, in contrast to clustering methods such as DBSCAN, G(r) does not identify or classify discrete clusters within a dataset. As such, it would not detect instances wherein two types of clusters (e.g. those in the nuclear interior vs. close to the lamina) might display different phenotypes.

To better understand the relationship between a given localization distribution and the G(r) curves, we used simulated datasets. We first started with a single target distributed within a ten-micron diameter spherical domain of similar size to a cell nucleus (**Fig S21A**). Within this domain, we simulated ten thousand points that were subject to varying spatial constraints. In a given simulation, a fraction of the ten thousand points were localized into spatial clusters of a given diameter while the remainder were randomly distributed within the domain. Within each cluster, the points were randomly distributed. We then individually varied the fraction clustered, the cluster diameter, and the number of clusters while keeping the other parameters constant. The G(r) curve for each "clustered" distribution was then normalized by the G(r) curve for a purely random distribution within the domain and this process was repeated for ten independent replicates to capture the simulated variability. As expected, we found that value of the G(r) curves increased corresponding to the fraction of clustered points (**Fig S21B**). Since the points within a cluster are randomly distributed, each curve also showed a plateau below the cluster diameter. The inflection point at which the G(r) curve deviated from complete spatial randomness (G(r) = 1) tracked with the cluster diameter (**Fig S21C**). Finally, as the number of clusters increased, we found that the value of the G(r) curves decreased (**Fig S21D**). Although somewhat counterintuitive, this can be explained because although the number of clusters varied, the total fraction of points within all clusters remained constant. Because the centroid of each cluster was randomly chosen, increasing the number of clusters within the dataset shifted the overall distribution closer to random.

Having established these general trends, we then simulated two separate targets using PCCF curves. As noted in the main text, PCCF is similar to Ripley's G(r) except that the localizations for one species serve as the "seeds" for the concentric shells within which we count the number of localizations of the second species. The null hypothesis is that the second species is randomly distributed relative to the first species. For these simulations, we fixed the fraction clustered = 0.5, the number of clusters = 20, and the cluster diameter = 500 nm. We then simulated 5 different scenarios: 1) Two randomly distributed species (**Fig S21E**), 2) Species 1 clustered while species 2 is random (**Fig S21F**), 3) Species 1 and species 2 are both clustered, but with randomly distributed cluster centroids (**Fig S21G**), 4) Species 1 and species 2 are both clustered with overlapping cluster centroids (**Fig S21H**), and 5) Species 1 and species 2 are both clustered with overlapping cluster centroids, but the two species are partitioned into separate halves of each cluster (**Fig S21I**). We then computed the PCCF curve for each scenario and plotted results from ten independent simulated replicates (**Fig S21J**). As expected, The PCCF curves for condition (1), the two random distributions, was approximately 1 for all length scales (**Fig S21J** – black). Condition (2) where the first species is clustered and the second species is random also showed PCCF ≈ 1 for all length scales (**Fig S21J** – blue). This is because the normalization for the two-species PCCF is performed using the original distribution for species 1 and a random distribution for species 2, i.e., the null hypothesis is not that both species are randomly distributed, but rather that the second species is randomly distributed relative to the first species. As such, the normalized PCCF curves are insensitive to the spatial distribution of the first "seed" species. For condition (3) where both species are clustered, but the centroids of the clusters are randomly distributed, we observed a small increase in the average PCCF curve that started at the cluster diameter and progressively increased at smaller length scales (**Fig S21J** – cyan), although this trend was variable, sometimes increasing partitioning (PCCF <1) and sometimes increasing co-clustering (PCCF >1). In contrast, the PCCF curve for two species that share cluster centroids, but that are randomly distributed relative to each other within each cluster (condition (4)) showed a clear increase in the PCCF curve that started at the cluster diameter and then plateaued at short length scales (**Fig S21J** – magenta). Finally, two species of molecules that cluster together at the 500 nm cluster diameter, but then partition into discrete regions within each cluster (condition 5) showed a clear increase at the cluster diameter followed by a peak near half the cluster diameter (~250 nm) and eventual decrease at shorter length scales (**Fig S21J** – green).

Overall, these simulations provide an intuitive picture of the relationship between point statistics and potential characteristics of the individual target distributions. However, it is important to also note the limitations of this approach. Due to the inherent averaging in the computation, there is not a one-to-one relationship between a given G(r) or PCCF curve and a target distribution. Many different distributions could give rise to the same curve. Thus, we do not suggest these simulations and our experimental analysis imply that a given target or group of targets displays clearly delineated clusters or domains of a given size. Rather, we support a more limited interpretation based on the mathematical definition: G(r) and PCCF curves report the average enrichment of one target with itself or of one target with a second target at varying length scales. Thus, this analysis reports the propensity for two molecules to be found at a given proximity to each other. The shapes of these curves describe whether two targets tend to be enriched or depleted at given scales, which relate to potential organizing principles. For example, two molecules may be enriched at one-micron length scale and then either randomly distributed or partitioned into discrete nanodomains below this scale which would support liquid-like diffusion or solid-like nanodomain models respectively. However, the curves do not provide direct information about whether this organization exists at a specific location within the cell.
